# The creatine transporter SLC6A8 is an immune pressure-exposed metabolic vulnerability in acute myeloid leukemia

**DOI:** 10.64898/2026.11.18.767047

**Authors:** Petya Apostolova, Tatjana Gremelspacher, Anthony Sonrel, Hui Hao-Shen, Minsun Cha, Massimo Andreis, Hanke Gwendolyn Bauersachs, Jacob L Martin, David E Sanin, Johannes Tossounidis, Alejandro Gomez-Auli, Katarzyna Buczak, Lukas M Braun, Jonathan D Curtis, David O’Sullivan, Franziska M Uhl, Gerhard Mittler, Livius Penter, Judith B Zaugg, Robert Zeiser, Edward J Pearce, Joerg M Buescher, Joy Edwards-Hicks, Beth Kelly, Erika L Pearce

## Abstract

Chemotherapy imposes metabolic pressure that generates persisting tumor cell states, but whether these adaptations also shape immune evasion is poorly understood. Here, we identify creatine accumulation as a conserved feature of chemotherapy-persisting acute myeloid leukemia (AML) across mouse and human models. Stable-isotope tracing and genetic perturbation show that this is driven predominantly by SLC6A8-mediated creatine uptake rather than de novo synthesis. SLC6A8 was dispensable for AML proliferation and chemotherapy resistance but strongly promoted leukemia persistence under T cell pressure. This effect was independent of phosphocreatine generation and canonical energetic buffering. Instead, SLC6A8 deficiency increased cytokine-driven STAT1 signaling and chromatin remodeling, inducing the expression of antigen-presentation and co-stimulatory molecules, such as MHC-II, CD40, and CD80. Thus, creatine uptake via SLC6A8 is a non-canonical metabolic adaptation that restrains immune-driven remodeling and enables AML immune evasion. These findings link a therapy-induced metabolic adaptation to cytokine responsiveness and reveal SLC6A8 as a targetable vulnerability.

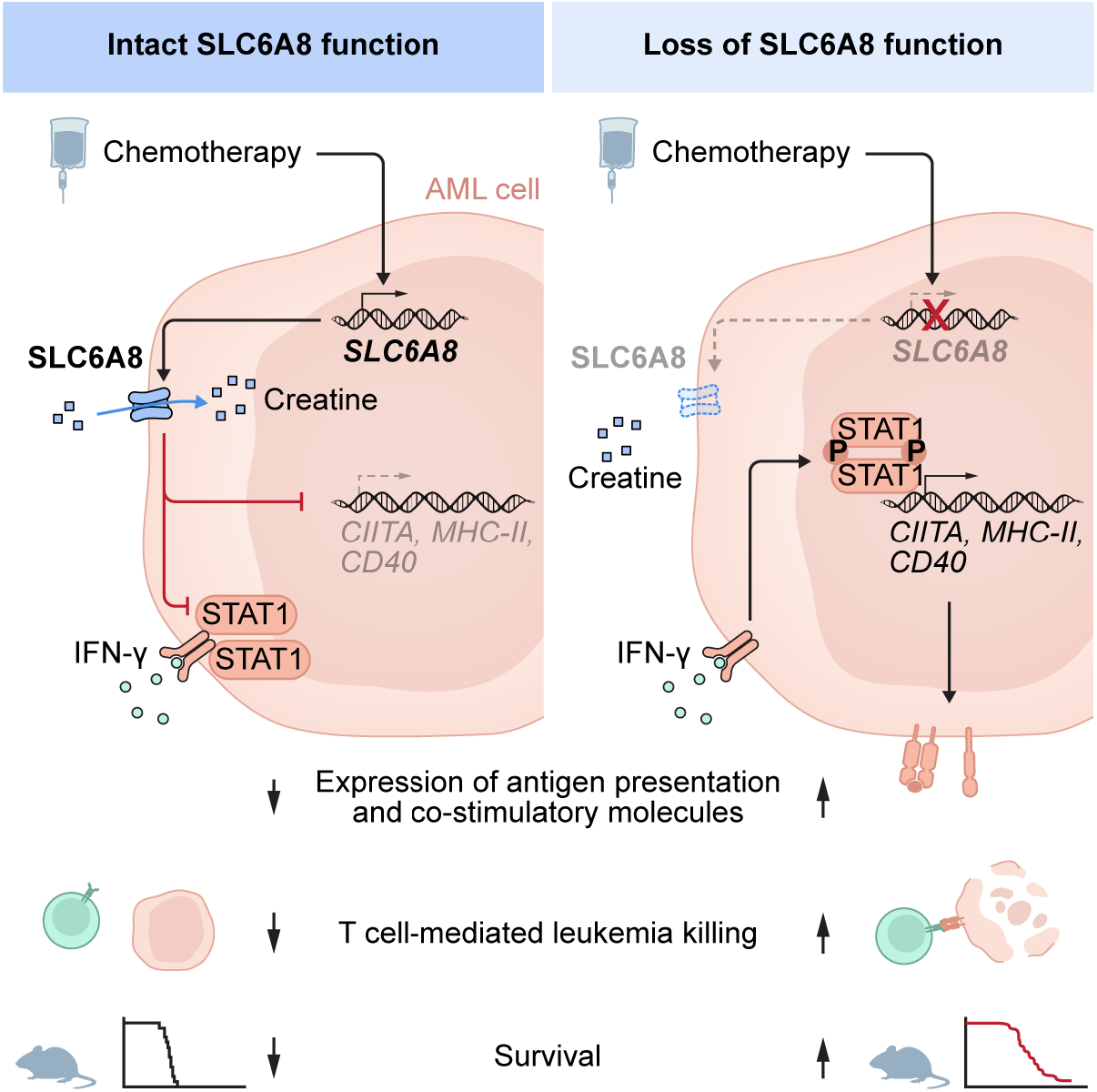
Graphical Abstract.

## Introduction

Cancer therapy imposes strong selective pressure that can drive the emergence of tumor cell states capable of surviving treatment^1–3^. Although acquired genetic alterations contribute to therapeutic resistance, non-genetic adaptations, including metabolic rewiring, also enable cancer cells to tolerate cytotoxic stress and persist after therapy^1,2,4,5^. Such therapy-persisting cells represent an important reservoir for relapse and disease progression^3,6^. However, cancer cells that survive chemotherapy must also evade immune-mediated elimination, particularly in settings in which anti-tumor immunity is therapeutically enhanced^7,8^. Whether chemotherapy-induced metabolic adaptations promote tumor immune evasion, and which specific molecular drivers underlie this process, remains incompletely understood.

Acute myeloid leukemia (AML) provides a clinically relevant model to address this question. AML is an aggressive hematologic malignancy characterized by impaired myeloid differentiation and accumulation of immature blasts in the bone marrow (BM), peripheral blood, and other tissues at the expense of normal hematopoiesis^9^. Although induction chemotherapy can induce remission, in many patients a subsequent allogeneic hematopoietic stem cell transplantation (allo-HSCT) is needed for durable disease control^10^. In this setting, donor-derived immune cells can eliminate residual leukemia cells through the graft-versus-leukemia (GVL) effect^11,12^, demonstrating that immune surveillance can provide potent anti-tumor activity. Nevertheless, relapse remains common and accounts for up to 50% of deaths in allo-HSCT recipients^13^, indicating that in many patients a subset of leukemic cells acquire or select for states that permit escape from both cytotoxic therapy and donor immune pressure. Current treatments for post-transplant relapse, including chemotherapy, donor lymphocyte infusions, and second allo-HSCT achieve limited long-term benefit with one-year survival of approximately 20%^14,15^. These poor outcomes underscore the need to identify tumor-intrinsic mechanisms that enable persistence and highlight AML relapse as an opportunity to study how tumor cells adapt at the interface of chemotherapy resistance and immune escape.

Metabolic rewiring is increasingly recognized as a central feature of chemotherapy resistance in AML^16–20^. Previous studies also indicate that AML can blunt the anti-tumor immune response by altering metabolite concentrations in the microenvironment^21^. For instance, AML-derived lactic acid suppresses cytotoxic CD8^+^ T cell function by blocking the transcription of metabolic genes and interfering with T cell glycolysis and proliferation^22^, and arginine catabolism via AML-expressed and secreted arginase II inhibits T cell proliferation and polarizes surrounding macrophages toward a tumor-promoting state^23,24^. These studies provide insights into how leukemia cells evade the immune response by impairing immune cell function through a metabolic remodeling of the microenvironment. However, whether chemotherapy induces metabolic states that drive cell-intrinsic resistance to immune-mediated killing and ultimately facilitate immune evasion is unknown.

## Results

### Creatine and glutathione accumulation are unifying features of chemotherapy-surviving AML cells

We hypothesized that chemotherapy-driven metabolic adaptations might contribute to an adaptive cell state associated with increased therapy resistance. We used untargeted high-performance liquid chromatography with high-resolution mass spectrometry (LC-MS)-based metabolomics to profile human HL-60 AML, mouse leukemia RMB-1, and primary murine AML cells (carrying *Mll-*PTD and *Flt3*-ITD mutations^25^). HL-60 and RMB-1 cells were exposed *in vitro* to the standard-of-care AML drugs cytarabine (CYT), daunorubicin (DAU), or azacytidine (AZA). The treatment doses were comparable to the concentrations reached in patient serum or leukemia cells^26–29^ and selected to induce comparable growth inhibition across treatments (**Fig. 1A, Fig. S1A-B**). After 24 h treatment, metabolites were extracted from persisting AML cells. In parallel, *Mll-*PTD *Flt3*-ITD AML-bearing mice were treated with AZA for 4 days, starting on day 14 after leukemia induction. Five days after the last injection, persisting CD117^+^ AML cells were enriched from the BM of animals for metabolomics analysis (**Fig. 1A**). AZA treatment significantly prolonged the survival of AML-bearing mice (**Fig. S1C**) and reduced the number of CD117^+^ AML cells in the BM (**Fig. S1D**), demonstrating that the selected treatment schedule induced selection pressure. Across all three models, principal component analysis performed on untargeted LC-MS metabolomics data showed clear treatment-associated separations, indicating that chemotherapy exposure induced substantial remodeling of the AML metabolome (**Fig. 1B-D**). Comparative analysis of >100 annotated polar metabolites further showed that each treatment altered a distinct, only partially overlapping metabolite subset (**Fig. 1E-G, Table S1-S3**). We compared the changes across models and treatments and identified that only two identified metabolites displayed concordant changes across all models: creatine and glutathione (**Fig. 1H, Fig. S1E**). Both metabolites were consistently enriched in chemotherapy-persisting AML cells (**Fig. 1I-K** and **Fig. S1F-H**). We also observed accumulation of phosphocreatine, the major downstream product of creatine which supports cellular energy homeostasis (**Fig. 1J-K**). To validate creatine accumulation as a unifying metabolic adaptation to chemotherapy in AML, we exposed primary patient samples and additional cell lines to AZA *in vitro* and observed consistently increased creatine abundance in chemotherapy-persisting cells (**Fig. 1L** and **Fig. S1I-J**). Independent support for this association was also evident when we analyzed a published dataset from an *in vivo* mouse MLL-AF9 AML model^19^, in which creatine abundance was elevated immediately after chemotherapy with CYT and doxorubicin, and, more pronounced, in the relapse setting (**Fig. 1M**).

**Figure 1.**
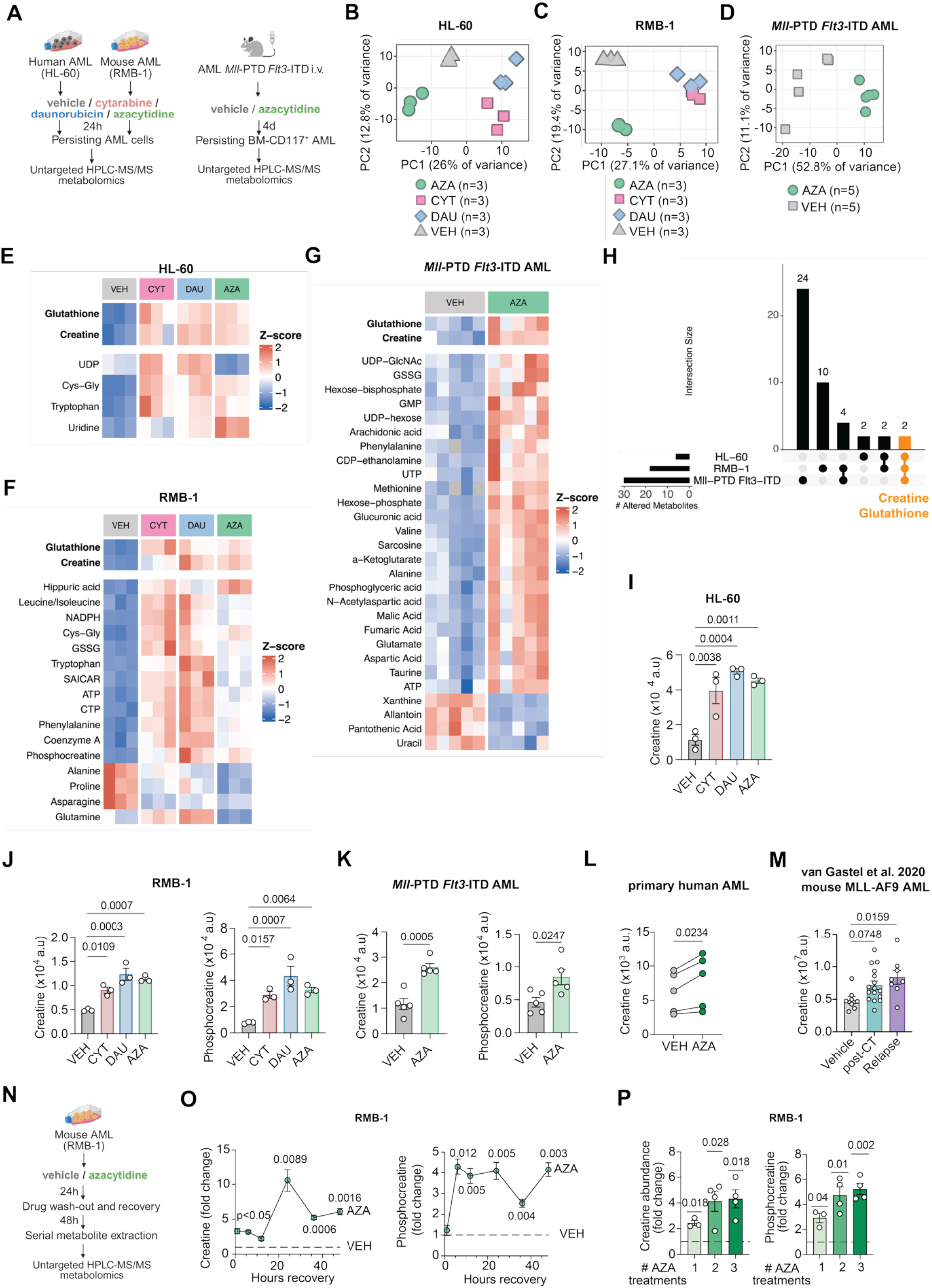
Chemotherapy-persisting AML cells accumulate intracellular creatine. (**A**) Left: Experimental setup for discovery metabolomics in HL-60 and RMB-1 cells *in vitro*. Right: Experimental setup for discovery metabolomics in *Mll-*PTD *Flt3*-ITD AML *in vivo*. Figure created with Biorender.com. **(B)-(D)** Principal component analysis (PCA) of non-targeted LC-MS metabolomics from chemotherapy-persisting HL-60 **(B)**, n=3/group, RMB-1 **(C)**, n=3/group, or *Mll-*PTD *Flt3*-ITD AML **(D)**, n=5/group. **(E)-(G)** Z-score of metabolite abundance for significantly changed metabolites across all treatments versus VEH in HL-60 and RMB-1 AML and for AZA against VEH in the *Mll-*PTD *Flt3*-ITD AML model. The top rows correspond to altered metabolites found in all models. The remaining rows were ordered by hierarchical clustering of Z-scored values. **(H)** UpSet plot of unique and overlapping significantly altered metabolites across models, highlighting in orange the metabolites altered in all models. **(I)** Creatine abundance by LC-MS in HL-60 cells exposed to the indicated treatments. Data are mean±SEM, n=3/group. **(J)** Creatine and phosphocreatine abundance by LC-MS in RMB-1 cells exposed to the indicated treatments. Data are mean±SEM, n=3/group. **(K)** Creatine and phosphocreatine abundance by LC-MS in CD117^+^ cells enriched from the BM of VEH– or AZA-treated *Mll-*PTD *Flt3*-ITD AML-bearing mice. Data are mean±SEM, n=5/group. **(L)** Creatine abundance by LC-MS in primary human AML cells treated with VEH or 5 µM AZA for 24h, n=5. **(M)** Creatine abundance by LC-MS in MLL-AF9-GFP cells sorted by flow cytometry from mice treated with vehicle (d18 after AML induction, n=8), cytarabine/doxorubicin chemotherapy (iCT, d18 after AML induction, n=16), or cytarabine/doxorubicin chemotherapy with 10 days post-treatment observation (relapse, d25 after AML induction, n=8) as measured by van Gastel et al^19^. Data are mean±SEM. **(N)** Experimental setup for metabolomics during chemotherapy recovery. Figure created with Biorender.com. **(O)** Time-resolved intracellular creatine and phosphocreatine abundance by LC-MS in AZA-persisting RMB-1 cells, relative to VEH-treated control at each time point. Data are mean±SEM, n=3-4/group. **(P)** Creatine and phosphocreatine abundance by LC-MS in RMB-1 cells exposed for 48 h to 2.5 µM AZA, followed by 6 days recovery without drug (1 cycle) for up to 3 cycles. Abundance was calculated relative to the respective vehicle control. Data are mean±SEM, n=3-4/group. **Abbreviations:** VEH: vehicle, CYT: cytarabine, DAU: daunorubicin; AZA: azacytidine. **Statistical analysis and data representation: (B)-(H)** Representative data from one of three independent experiments for HL-60 and RMB-1 and one of two independent experiments for the *Mll-*PTD *Flt3*-ITD AML model. **(B)-(D)** PCA was performed on log_2_-transformed and Pareto-scaled data (mean-centered, divided by √SD). **(E)-(H)** Statistical analysis was performed per metabolite using One-Way ANOVA (group as factor) followed by Tukey HSD post-hoc test for the HL-60 and RMB-1 data against VEH and with unpaired two-tailed t-test for the *Mll-*PTD *Flt3*-ITD AML model. In each model, p-values were adjusted using the Benjamini-Hochberg (BH) correction. **(I), (J)** Statistical analysis with One-way ANOVA followed by Dunnett’s multiple comparisons test. Representative data from one of three independent experiments. **(K)** Statistical analysis with unpaired two-tailed t-test. Representative data from one of two independent experiments. **(L)** Technical triplicates were run from each patient/treatment, and the average value was calculated. Statistical analysis with paired two-tailed t-test. (**M)** Statistical analysis with One-Way ANOVA followed by Dunnett’s multiple comparisons test. **(O)-(P)** Statistical analysis with One-Sample t-test. Representative data from one of two independent experiments.

Because glutathione’s role in cancer has been extensively studied, with well-established functions in buffering reactive oxygen species, detoxifying chemotherapy agents, and promoting therapy resistance across multiple malignancies including AML^20,30,31^, we focused on creatine and prioritized AZA for mechanistic studies, given its importance in both pre– and post-allo-HSCT settings^32^. We asked whether the creatine-high state persists beyond acute drug exposure and treated RMB-1 AML cells with AZA for 24h, followed by a drug wash-out and serial metabolite harvest over a recovery period of 48h (**Fig. 1N**). Intracellular creatine and phosphocreatine remained elevated during the recovery period, indicating that the adaptive metabolic state persists beyond acute drug exposure (**Fig. 1O**). Moreover, repeated cycles of AZA treatment led to progressively higher intracellular creatine and phosphocreatine abundance, consistent with reinforced metabolic adaptations under recurrent therapeutic stress (**Fig. 1P**). Supporting the clinical relevance of our findings, higher creatine abundance was also found in a published dataset of leukemic stem cells (LSCs) from patients with relapsed/refractory AML compared to LSCs from patients with untreated AML^16^.

### Chemotherapy-persisting AML cells accumulate creatine via SLC6A8-mediated uptake

Creatine can be acquired by cells via the solute carrier family 6 member 8 (SLC6A8) transporter or synthesized *de novo* from arginine and glycine via arginine:glycine amidinotransferase (AGAT, encoded by *GATM*) and guanidinoacetate N-methyltransferase (encoded by *GAMT*) (**Fig. 2A**). To assess how much of the intracellular creatine pool is derived from uptake, we performed stable-isotope tracing adding deuterated creatine (d_3_-creatine) to the media of vehicle– (VEH-) or AZA-treated AML cells (40 µM, corresponding to the concentration in human plasma, **Fig. 2B**). As expected, we observed that AZA-treated cells had higher intracellular creatine abundance compared to VEH counterparts, and most of this creatine was deuterated, indicating uptake from the media (**Fig. 2C**). Intracellular phosphocreatine abundance was also higher in AZA-compared to VEH-treated cells, with deuterated phosphocreatine comprising the major portion suggesting that most of phosphocreatine was made from the uptake-derived creatine (**Fig. 2C**). After 24 h of stable-isotope labeling, the fraction of d_3_-creatine increased to 84% in AZA-persisting cells compared to 72% in VEH-treated cells (**Fig. S2A**). Similarly, the proportion of labeled phosphocreatine was 55% in the VEH condition and 70% after AZA exposure (**Fig. S2A**). An even stronger predominance of creatine uptake was observed in human HL-60 AML cells, where ∼95% of intracellular creatine and phosphocreatine was deuterated (**Fig. 2D, Fig. S2B**). We also quantified the contribution of *de novo* synthesis to the creatine pool by stable-isotope tracing adding 100 µM U-^13^C-arginine (M+6) to the media of VEH– or AZA-pretreated AML cells. During creatine *de novo* synthesis, an amidino group from arginine is transferred to glycine by AGAT (encoded by *GATM,* rate-limiting step). This intermediate, guanidinoacetate (GAA), is further methylated to creatine by GAMT (**Fig. 2A**). This process results in one ^13^C atom from U-^13^C-arginine being transferred to creatine (M+1 labeling, **Fig. 2E**). RMB-1 AML readily took up labeled arginine (**Fig. 2F**, M+6 fraction of arginine), but only a minor fraction of creatine and phosphocreatine was labeled (**Fig. 2F**, M+1 fraction of creatine and phosphocreatine). The fraction of intracellular creatine and phosphocreatine derived from labeled arginine was only ∼15% and ∼30% of the total creatine and phosphocreatine, respectively, and was further diminished by AZA treatment (**Fig. S2C**). In HL-60 cells, the fractional contribution of labeled arginine to intracellular creatine and phosphocreatine was <1% and ∼3%, respectively (**Fig. 2G, Fig. S2D**). Supporting that AZA-persisting AML cells enrich creatine via uptake, we found lower creatine amounts in the media of AZA-persisting RMB-1 cells compared to VEH-treated controls and RPMI without cells (**Fig. 2H**). We assessed whether AZA treatment regulated *Slc6a8/SLC6A8* and *Gatm/GATM* expression. *Gatm*/*GATM* mRNA levels remained unchanged, while *Slc6a8*/*SLC6A8* transcripts were higher in AML cell lines and primary AML samples after AZA exposure (**Fig. 2I-K**).

**Figure 2.**
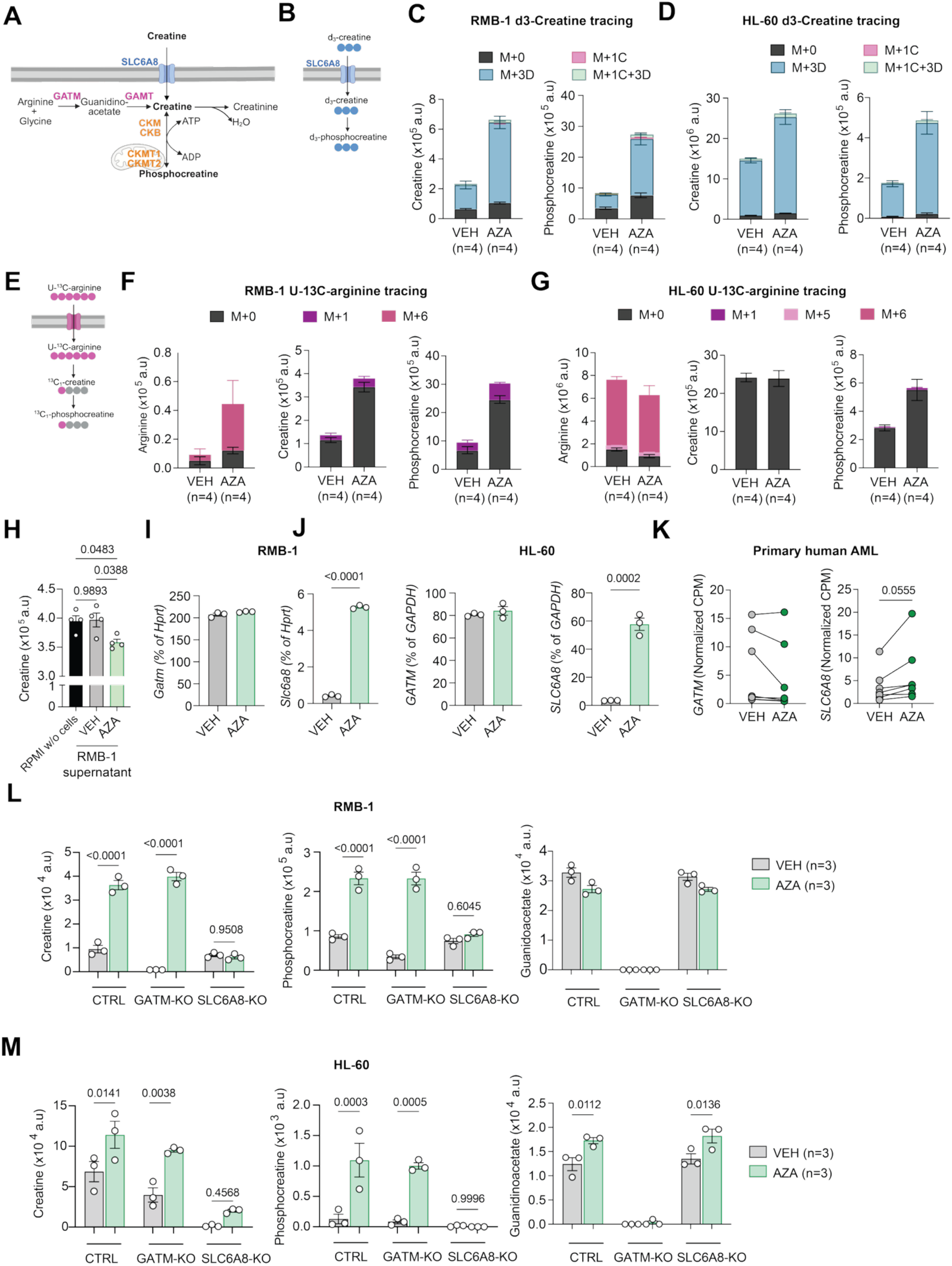
Chemotherapy-persisting AML cells accumulate creatine via SLC6A8-mediated uptake. (**A**) Overview of creatine metabolism. Figure created with Biorender.com. **(B)** Intracellular metabolism of d_3_-creatine upon environmental uptake. **(C)-(D)** d_3_-creatine tracing by LC-MS in RMB-1 (C) or HL-60 AML (D) pre-treated with vehicle (VEH) or azacytidine (AZA) for 24h. Labeling was performed for 24 h. Absolute abundance of creatine and phosphocreatine isotopes. M+0 denotes the proportion of unlabeled intracellular creatine, M+3D: d_3_-creatine, M+1C: creatine with one ^13^C-atom (natural isotope occurrence), M+1C+3D: d_3_-creatine with one ^13^C-atom. Data are mean±SEM, n=3-4/group. **(E)** Uptake and intracellular metabolism of U-^13^C-arginine for stable-isotope tracing in (F)-(G). **(F)-(G)** U-^13^C-arginine tracing by LC-MS in RMB-1 (F) or HL-60 AML (G) pre-treated with VEH or AZA for 24h. Labeling was performed for 24 h. Absolute abundance of arginine, creatine and phosphocreatine isotopes. M+0 denotes unlabeled metabolites, M+6: arginine with six ^13^C atoms, M+5: arginine with five ^13^C atoms, M+1: creatine and phosphocreatine with one ^13^C atom. Data are mean±SEM, n=4/group. **(H)** Creatine abundance in complete RPMI without cells or in the supernatant of VEH-/AZA-pretreated RMB-1 cells cultured for 24 h after drug removal. Data are mean±SEM, n=4/group. **(I)-(J)** *Gatm/GATM* and *Slc6a8/SLC6A8* mRNA by qPCR in RMB-1 **(I)** and HL-60 **(J)** cells treated with VEH or AZA for 24 h. Data are mean±SEM, n=3/group. *Hprt* was used as housekeeping gene for RMB-1 and *GAPDH* for HL-60. **(K)** *GATM* and *SLC6A8* mRNA by RNA-sequencing in primary human AML treated with VEH or 5 µM AZA for 24 h, n=7 patients. **(L)** RMB-1 AML cells underwent gene editing by CRISPR/Cas9 for *Gatm* (GATM-KO) or *Slc6a8* (SLC6A8-KO). Control (CTRL) cells were edited with a non-targeting control gRNA. Creatine, phosphocreatine, and guanidinoacetate abundance by LC-MS following treatment with VEH or AZA for 24 h. Data are mean±SEM, n=3/group. **(M)** HL-60 AML cells underwent gene editing by CRISPR/Cas9 for *GATM* (GATM-KO) or *SLC6A8* (SLC6A8-KO). Control (CTRL) cells were edited with a non-targeting control gRNA. Creatine, phosphocreatine, and guanidinoacetate abundance by LC-MS following treatment with VEH or AZA for 24 h. Data are mean±SEM, n=3/group. **Statistical analysis and data representation: (C)-(G)** Representative data from one of two independent experiments. **(H)** Statistical analysis by One-way ANOVA followed by Tukey’s multiple comparisons test. Representative data from one of two independent experiments. **(I)-(J)** Statistical analysis by unpaired two-tailed t-test. Representative data from one of three independent experiments. **(K)** Technical triplicates were run from each patient/treatment, and the average value was calculated. Statistical analysis with paired two-tailed t-test. **(L)-(M)** Statistical analysis with One-way ANOVA followed by Tukey’s multiple comparisons test. Representative data from one of two independent experiments.

To establish contribution of GATM and SLC6A8 to creatine accumulation in AML, we used CRISPR/Cas9 to delete *Gatm/GATM* or *Slc6a8/SLC6A8* in AML cells (GATM-KO and SLC6A8-KO). Non-targeting gRNA were controls. Successful deletion was verified by immunoblotting or Inference of CRISPR edits (ICE) (**Fig. S2E-F**). Notably, only perturbing *Slc6a8*/*SLC6A8* disrupted AZA-induced creatine and phosphocreatine accumulation measured both by LC-MS and by a colorimetric assay (**Fig. 2L-M, Fig. S2G**). While the de novo synthesis intermediate GAA was depleted in Gatm/GATM-KO cells, demonstrating that the *de novo* pathway was fully blocked (**Fig. 2L-M**), this did not abolish creatine and phosphocreatine accumulation in AZA-exposed cells. Together, these data show that AZA-persisting leukemia accumulate creatine via selective SLC6A8 upregulation.

### SLC6A8 is a selective metabolic vulnerability under immune pressure in AML

To define the functional role of SLC6A8 for leukemia progression, we first engrafted immunodeficient NSG mice with CTRL or SLC6A8-KO mCherry^+^ RMB-1 AML cells and treated them with VEH or AZA (**Fig. 3A**). Deleting SLC6A8 did not affect mouse survival (**Fig. 3B**) or BM or spleen leukemia burden measured by flow cytometry (**Fig. 3C-D**), irrespective of whether AZA treatment was administered or not. Given these *in vivo* data, we hypothesized that SLC6A8 targeting does not impair AML cell proliferation. Indeed, growth rates in standard culture conditions and the sensitivity of SLC6A8-deficient RMB-1 cells to chemotherapy treatment *in vitro* were not altered (**Fig. 3E, Fig. S3A**). Moreover, migration along a CXCL12 gradient, representing a major chemotactic stimulus for hematopoietic cells, was unchanged, suggesting that lack of functional creatine uptake did not impair AML migration to the BM (**Fig. 3F**). We next asked whether SLC6A8 plays a functional role in leukemia immune evasion. We injected AZA-pretreated CTRL or SLC6A8-KO RMB-1 cells into immunocompetent BALB/c mice undergoing allo-HSCT with transfer of C57BL/6 BM and CD4^+^/CD8^+^ T cells (**Fig. 3G**). Notably, mice receiving SLC6A8-deficient AML cells in combination with allogeneic T cells survived significantly longer than mice receiving CTRL AML cells (**Fig. 3H**). The longer survival was accompanied by a lower percentage and absolute number of mCherry^+^ AML cells in BM and spleen (**Fig. 3I-J**). In line with the enhanced *in vivo* elimination, SLC6A8-KO leukemia was more sensitive to allogeneic T cells in an *in vitro* cytolysis assay (**Fig. 3K**). These results demonstrate that SLC6A8 supports leukemia persistence in the presence of an allogeneic immune response, while it is dispensable in an immunodeficient environment.

**Figure 3.**
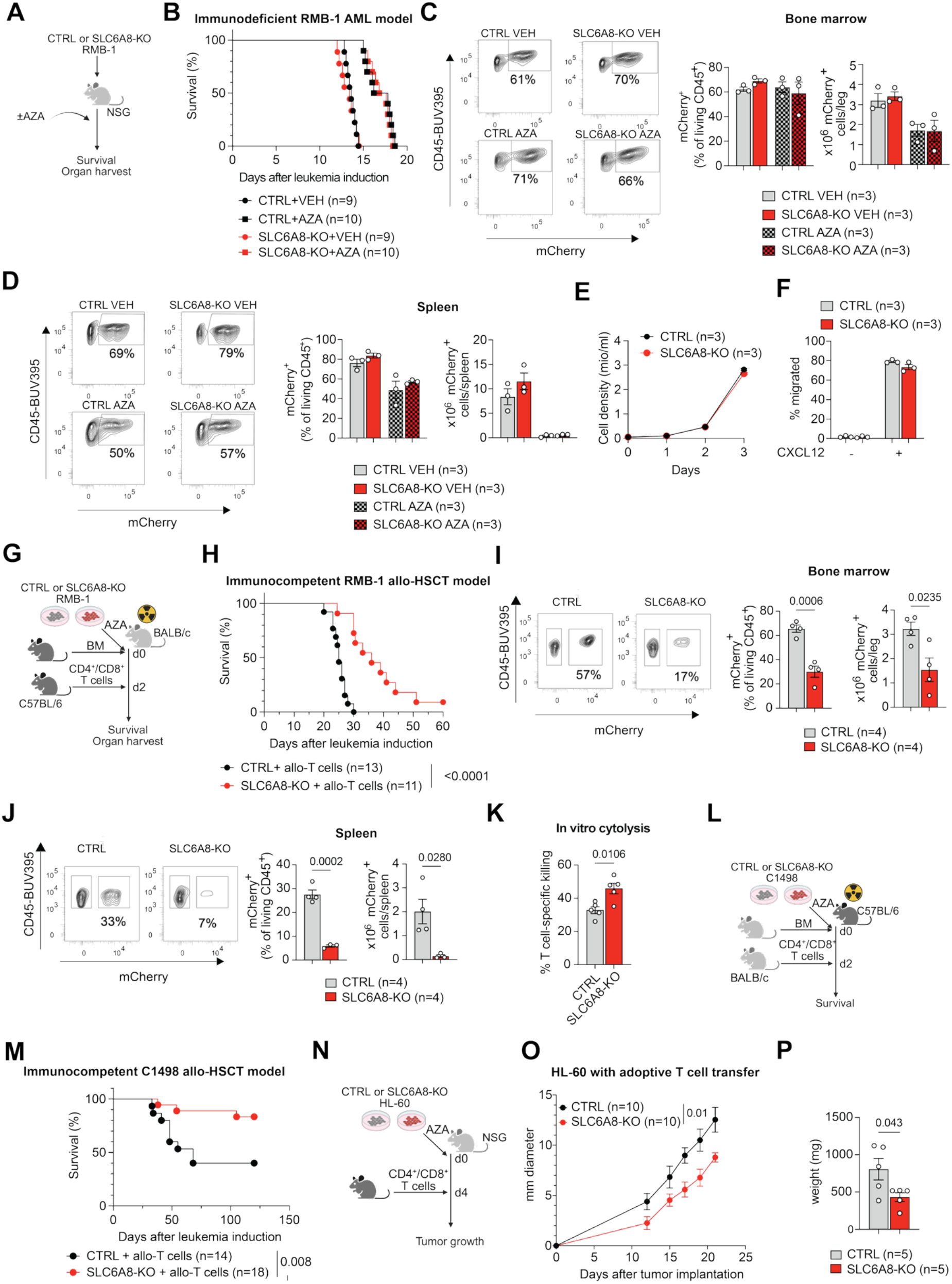
SLC6A8 is a selective metabolic vulnerability under immune pressure in AML. (**A**) Experimental workflow for (B)-(D), created with Biorender.com. **(B)** Survival of NSG mice engrafted with CTRL or SLC6A8-KO RMB-1 cells ± AZA treatment, n=9-10/group as indicated. **(C)** Left: representative flow cytometry contour plots from the BM of mice with AML. Middle and right: relative and absolute number of mCherry^+^ AML cells. Data are mean±SEM, n=3/group. **(D)** Left: representative flow cytometry contour plots from the spleen of mice with AML. Middle and right: relative and absolute number of mCherry^+^ AML cells. Data are mean±SEM, n=3/group. **(E)** Cell density of CTRL and SLC6A8-KO RMB-1 cells over a time course of 3 days. Data are mean±SEM, n=3/group. **(F)** Percentage migrated cells in a transwell migration assay ± a CXCL12 gradient. Data are mean±SEM, n=3/group. **(G)** Experimental workflow for (H)-(J), created with Biorender.com. **(H)** Survival of BALB/c mice engrafted with AZA-treated CTRL or SLC6A8-KO RMB-1 cells and allogeneic C57BL/6 BM and T cells (n=11-13/group as indicated). **(I)** Left: representative flow cytometry contour plots from the BM of mice with AML. Middle and right: relative and absolute number of mCherry^+^ AML cells. Data are mean±SEM, n=4/group. **(J)** Left: representative flow cytometry contour plots from the spleen of mice with AML. Middle and right: relative and absolute number of mCherry^+^ AML cells. Data are mean±SEM, n=3/group. **(K)** RMB-1 CTRL and SLC6A8-KO cells were treated with 5 µM AZA for 24 h, followed by co-culture with activated allogeneic (C57BL/6) CD3^+^ T cells in a 1:1 ratio for 24 h. Percentage of T cell-specific killing. Data are mean±SEM, n=5/group. **(L)** Experimental workflow for (M), created with Biorender.com. **(M)** Survival of C57BL/6 mice engrafted with AZA-treated CTRL or SLC6A8-KO C1498 cells and allogeneic BALB/c BM and T cells, n=14-18/group as indicated. **(N)** Experimental workflow for (O)-(P), created with Biorender.com. **(O)** Average tumor diameter over time. Data are mean±SEM, n=10/group. **(P)** Weight of explanted tumors on d22. Data are mean±SEM, n=5/group. **Statistical analysis and data representation**: **(B)** Statistical analysis with Mantel-Cox test (p>0.05). Pooled data from two independent experiments. **(C)-(D), (F)** Statistical analysis with Two-Way ANOVA with correction for multiple comparisons (adjusted p-values>0.05). Representative data from one of two independent experiments. **(H)** Statistical analysis with Mantel-Cox test. Pooled data from three independent experiments. **(I)-(K)** Statistical analysis with unpaired two-tailed t-test. Representative data from one of two or three independent experiments. **(M)** Statistical analysis with Mantel-Cox test. Pooled data from three independent experiments. **(O)** Statistical analysis with Two-Way ANOVA (p-value for effect of the experimental group indicated). Pooled data from two independent experiments. **(P)** Statistical analysis with unpaired two-tailed t-test. Representative data from one of two independent experiments.

To investigate whether SLC6A8 has a similar regulation and function in other AML models, we employed C1498 AML cells, a mouse AML cell line^33^ on a C57BL/6 background suitable for engraftment in an immunocompetent allo-HSCT model and the study of anti-tumor immune responses. Similarly to our observations in other AML cell lines, AZA induced *Slc6a8* expression and creatine accumulation in C1498 cells (**Fig. S3B-C**). SLC6A8 deletion through CRISPR-Cas9 (**Fig. S3D**) resulted in reduced intracellular creatine (**Fig. S3E**) but did not affect *in vitro* proliferation (**Fig. S3F**), chemotherapy sensitivity (**Fig. S3G**) or engraftment in immunodeficient mice (**Fig. S3H-I**), indicating that SLC6A8 was dispensable for *in vitro* and *in vivo* growth and chemotherapy persistence. Next, we injected AZA-pretreated C1498 CTRL or SLC6A8-KO cells alongside allogeneic BALB/c BM into irradiated C57BL/6 recipient mice. Two days later, BALB/c CD4^+^ and CD8^+^ T cells were transferred (**Fig. 3L**). Recipients of SLC6A8-KO AML cells survived significantly longer, indicating better immune control of leukemia (**Fig. 3M**). As a third AML model, we studied the previously generated HL-60 CTRL and SLC6A8-KO cells. SLC6A8 deficiency did not alter *in vitro* proliferation (**Fig. S3J**) or chemotherapy sensitivity (**Fig. S3K**). To study the anti-tumor immune response, we subcutaneously injected immunodeficient mice with HL-60 CTRL or SLC6A8-KO cells, followed by a transfer of C57BL/6 CD4^+^ and CD8^+^ T cells, and measured tumor size (**Fig. 3N**). SLC6A8-KO AML cells formed smaller tumors compared to CTRL, as shown by longitudinal size measurement (**Fig. 3O**) and tumor weight at terminal harvest (day 22, **Fig. 3P**). The data show that deleting SLC6A8 impairs leukemia progression under chemo-immune pressure in three independent AML models (summarized in **Table S4**), suggesting that chemotherapy-induced SLC6A8 expression protects AML from the anti-tumor immune response.

### SLC6A8 loss enhances T cell activation and requires combined CD4^+^ and CD8^+^ T cell pressure for leukemia control

Next, we characterized the immune compartment in CTRL and SLC6A8-KO leukemia-bearing mice (RMB-1 model with allo-HSCT, Fig. 3G). While the percentage and absolute number of allogeneic CD4^+^ and CD8^+^ T cells in the BM did not differ (**Fig. S4A-B**), or appeared slightly lower, we found higher expression of the T cell activation and memory marker CD44 on T cells of mice with SLC6A8-KO AML compared to CTRL, perhaps indicating that these cells had undergone a stronger TCR activation (**Fig. S4C-D**). To dissect the contributions of CD4^+^ and CD8^+^ subsets, we performed allo-HSCT with the transfer of only CD4^+^ T cells, only CD8^+^ cells, no T cells, or combined CD4^+^ and CD8^+^ T cells in BALB/c mice. To counteract engraftment of the non-transferred T cells from the allogeneic BALB/c BM, the respective depleting antibody was administered (e.g. mice receiving CD4^+^ T cells also received an anti-CD8 antibody, **Fig. S4E**). With this setup, we confirmed that CD4^+^ and CD8^+^ T cell depletion abolished survival differences in mice receiving CTRL and SLC6A8-KO cells (**Fig. S4F**). Notably, deleting SLC6A8 did not provide a survival advantage when only CD4^+^ T cells were transferred (**Fig. S5G**), while a small, but not statistically significant difference between the survival of CTRL and SLC6A8-KO AML-bearing mice was found when only CD8^+^ T cells were transferred (**Fig. S4H**). These data demonstrate that the full phenotypic disadvantage of SLC6A8-KO AML cells is only unmasked under combined CD4^+^ and CD8^+^ T cell pressure. Taken together, our data show that SLC6A8 is dispensable for AML proliferation, migration, chemotherapy resistance, and engraftment in immunocompromised mice, but its loss exposes leukemia to T cell-mediated killing *in vivo* and *in vitro*, and that this effect depends on the presence of both CD4^+^ and CD8^+^ T cell subsets.

### Creatine uptake is the major function of SLC6A8

SLC6A8 is a high-affinity sodium– and chloride-dependent transporter, transporting one creatine molecule across the plasma membrane alongside one chloride and two sodium ions^34,35^. Although its affinity for creatine is high^34^, potential other substrates may exist. The creatine precursor GAA has been described as a substrate for SLC6A8, albeit with a K_m_ value ∼10-fold greater than that for creatine^36^. To assess whether SLC6A8 deficiency limited the uptake of substrates other than creatine, potentially contributing to the observed functional phenotype, we analyzed the non-targeted LC-MS data from SLC6A8-KO and CTRL AML. Notably, SLC6A8-KO did not reduce intracellular GAA (**Fig. 2L-M**), ruling out reduced GAA uptake as the molecular link between SLC6A8 loss and improved T cell-mediated elimination of AML. Metabolomics identified 15 metabolites with significant changes upon SLC6A8-KO in RMB-1 leukemia cells ± AZA treatment (**Fig. S5A**). Of these, three displayed the pattern we expected to be directly linked to SLC6A8 function: they were enriched after AZA exposure in CTRL, but not in SLC6A8-KO cells. These metabolites were creatine, phosphocreatine, and phosphoglycerate (**Fig. S5A, B**). 2-/3-phosphoglycerate are glycolysis intermediates primarily linked to intracellular metabolism, although extracellular pools may exist^37^. Of note, phosphoglycerate was unchanged in SLC6A8-KO vs. CTRL HL-60 cells (**Fig. S5C**), demonstrating that SLC6A8 loss does not uniformly result in altered intracellular levels of this metabolite. Thus, the literature and our metabolomics data suggest that creatine uptake is the major metabolic function of SLC6A8.

### SLC6A8-mediated creatine uptake promotes AML persistence under immune pressure independently of creatine kinase-mediated energy buffering

One of the key intracellular functions of creatine is to maintain energy homeostasis^38^. The enzyme creatine kinase (CK) catalyzes the transfer of a phosphoryl group from ATP to creatine, yielding phosphocreatine and ADP (**Fig. 2A**). Creatine and phosphocreatine provide an intracellular phosphate-buffering system essential for maintaining ATP supply in tissues with high energy demands^38^. We found that phosphocreatine, like creatine, accumulated in chemotherapy-surviving AML cells (**Fig. 1J-K**, **Fig. 2L-M**). In line with higher phosphocreatine abundance, CK activity increased in AZA-persisting RMB-1 cells (**Fig. 4A**). CK has four isoforms: two cytosolic, CK muscle-type (CKM) and CK brain-type (CKB), and two mitochondrial, CK ubiquitous-type mitochondrial (CKMT1) and CK sarcomeric-type mitochondrial (CKMT2), all of which are encoded by different genes^39–41^. AZA treatment induced CK mRNA in RMB-1 and HL-60 cells (**Fig. 4B, Fig. S6A**). *Ckb/CKB* was the most abundant isoform and its protein level also increased (**Fig. 4C**). To define the contribution of individual CK isoforms to phosphocreatine generation, we deleted *Ckm*, *Ckb*, and *Ckmt1* (as *Ckmt2* expression was not detected) in RMB-1 cells by CRISPR/Cas9. We found that CKB-KO (**Fig. S6B**) had non-detectable CK activity in standard culture conditions and after AZA exposure (**Fig. 4D**). Consequently, no phosphocreatine accumulation was observed in CKB-KO cells (**Fig. 4E**). CK activity in response to AZA was also slightly reduced in CKM-KO cells, but this was not reflected by changes in phosphocreatine (**Fig. 4D-E**), underlining that CKB was the main isoform responsible for phosphocreatine generation in RMB-1 cells. *In vitro* cell proliferation and chemotherapy sensitivity were not changed between CKB-KO and CTRL RMB-1 cells (**Fig. S6C-D**). Next, we induced AML in immunodeficient NSG mice by transferring CTRL or CKB-KO AML cells, and performed AZA treatment for 5 days, starting on d7 after leukemia induction (**Fig. 4F**). Similarly to what we observed with SLC6A8-KO AML cells, disrupting CKB expression did not affect the survival of immunodeficient leukemia-bearing mice, either with or without AZA (**Fig. 4G)**. To test leukemia progression under immune pressure, we transferred AZA-pretreated CTRL and CKB-KO AML in BALB/c recipients undergoing allo-HSCT (**Fig. 4H**). Notably, mice receiving CKB-KO AML had neither improved survival nor reduced leukemia burden compared to mice receiving CTRL AML (**Fig. 4I, J**). Because CKB-KO cells have a defect in phosphocreatine generation, but not in SLC6A8-mediated creatine uptake, these data indicated that SLC6A8 affected leukemia survival under T cell pressure through a function independent of canonical energy buffering via phosphocreatine.

**Figure 4.**
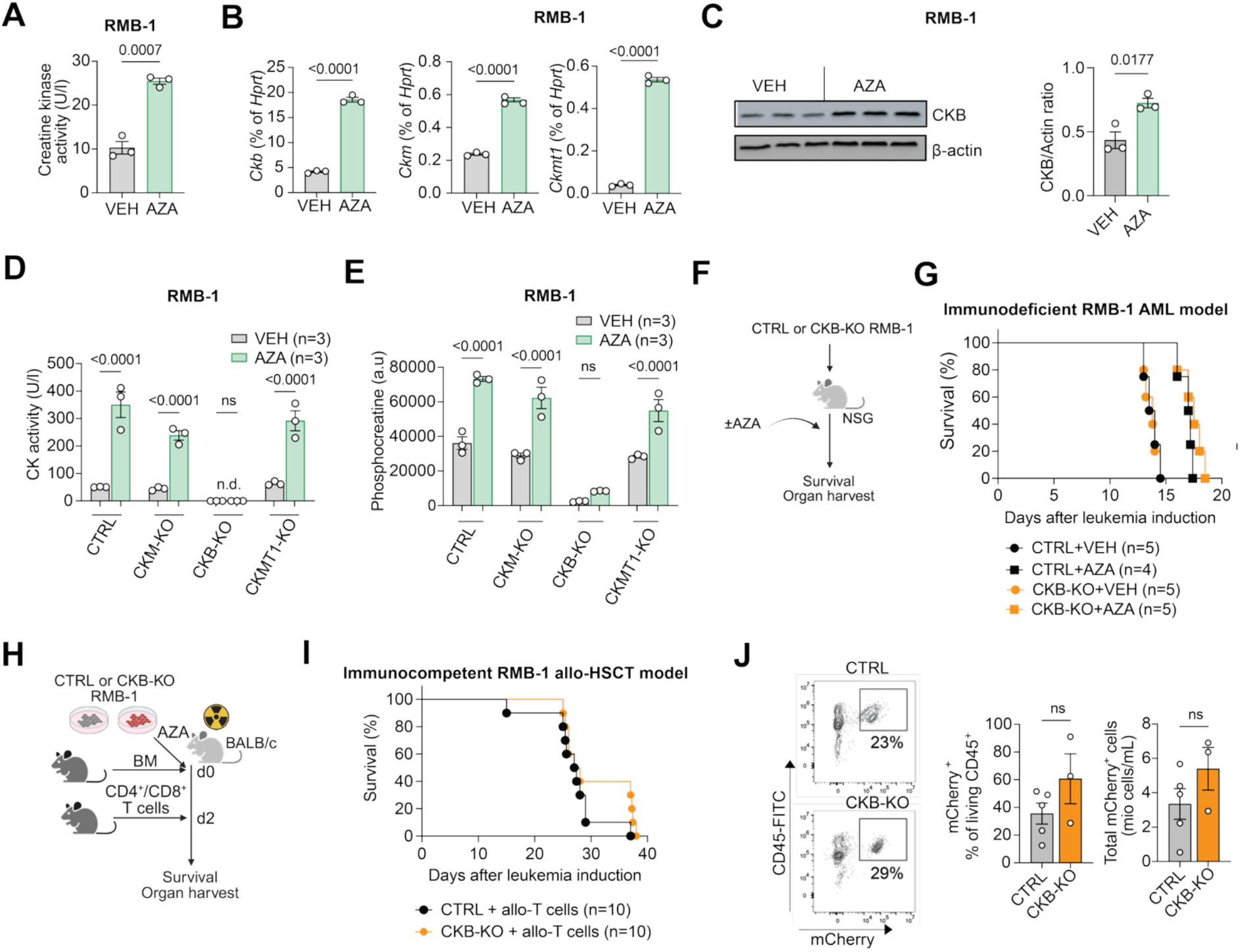
Creatine uptake promotes AML persistence under immune pressure through a phosphocreatine-independent mechanism. (**A**) Creatine kinase activity determined by an enzyme activity assay in RMB-1 leukemia treated with vehicle (VEH) or 2.5 µM azacytidine (AZA) for 24 h. Data are mean±SEM, n=3/group. **(B)** *Ckb*, *Ckm*, and *Ckmt1* mRNA by qPCR in RMB-1 cells treated with VEH or 2.5 µM AZA for 24 h. *Hprt* was used as housekeeping gene. Data are mean±SEM, n=3/group. **(C)** CKB protein expression by western blot in RMB-1 cells treated with VEH or 2.5 µM AZA for 24 h. Actin was used as loading control. Data are mean±SEM, n=3/group. **(D)** CKM-, CKB-, and CKMT1-KO RMB-1 cells were generated by CRISPR/Cas9 gene editing, treated with AZA 2.5 µM for 24h, and creatine kinase activity was determined by an enzyme activity assay. Data are mean±SEM, n=3/group. **(E)** Phosphocreatine abundance by LC-MS in RMB-1 cells treated as in (D). Data are mean±SEM, n=3/group. **(F)** Experimental workflow for (G), created with Biorender.com. **(G)** Survival of NSG mice engrafted with CTRL or CKB-KO RMB-1 cells ± AZA treatment, n=4-5/group as indicated. **(H)** Experimental workflow for (I)-(J), created with Biorender.com. **(I)** Survival of BALB/c mice engrafted with AZA-pretreated CTRL or CKB-KO RMB-1 cells and allogeneic C57BL/6 BM and T cells, n=10/group. **(J)** Left: representative flow cytometry contour plots from the BM of mice with AML. Right: relative and absolute number of mCherry^+^ AML cells. Data are mean±SEM, n=3-5/group. **Statistical analysis and data representation: (A)-(C)** Statistical analysis with unpaired two-tailed t-test. Representative data from one of two independent experiments. **(D)-(E)** Statistical analysis with One-Way ANOVA with Dunnett’s correction for multiple comparisons. Representative data from one of two independent experiments. **(G), (I)** Statistical analysis with Mantel-Cox test, p>0.05. Data from one (G) or two (I) independent experiments. **(J)** Statistical analysis with unpaired two-tailed t-test.

### SLC6A8 restrains immune pressure-induced inflammatory and antigen-presenting state remodeling in AML

We hypothesized that two potential mechanisms could explain how SLC6A8-mediated creatine uptake by leukemia cells may shape tumor-immune crosstalk. First, AZA-persisting AML cells might deplete creatine from the environment. Because creatine uptake is essential for maintaining the proper function of tumor-infiltrating lymphocytes^42^ and T cells during infection^43^, its depletion might impair anti-leukemia T cell immunity. Second, intracellular creatine abundance may drive cell-intrinsic AML resistance to T cell recognition or killing.

To test the first mechanism, we investigated whether supplementing creatine by feeding mice a creatine-rich diet could restore leukemia immune control. We performed allo-HSCT in mice engrafted with AZA-pretreated RMB-1 or *Mll*-PTD *Flt3*-ITD AML cells and fed them a standard diet or a 5% creatine-containing diet (**Fig. S7A**). Dietary creatine supplementation failed to improve survival and instead shortened survival in both AML models (**Fig. S7B-C**), arguing against the hypothesis that increased creatine availability would enhance immune-mediated leukemia control in these experimental settings.

To investigate how leukemia-intrinsic SLC6A8 influences the response to immune pressure, we compared the transcriptional states of CTRL and SLC6A8-deficient leukemia (RMB-1) under homeostatic conditions and following sequential exposure to AZA and allogeneic T cells (**Fig. 5A**). Analysis of differentially expressed genes (DEG, defined as FDR<0.05 and |log_2_ fold change| ≧ 0.5) identified 622 upregulated DEG and 436 downregulated DEG in SLC6A8-KO compared to CTRL RMB-1 cells under homeostatic conditions (VEH, **Fig. 5B**). When cells were exposed to AZA+T cells, there were 791 upregulated and 291 downregulated DEG (**Fig. 5B**). Combined AZA+T cell treatment elicited markedly different transcriptional remodeling at the pathway level in SLC6A8-KO compared with CTRL leukemia cells (**Fig. 5C**). Gene-set enrichment analysis (GSEA) identified increased inflammatory, IFN-γ-response, TNF/NF-κB and IL6/JAK/STAT3-associated programs in SLC6A8-deficient cells (**Fig. 5D-H**). To identify transcriptional regulators associated with this inflammatory state, we inferred transcription factor (TF) activity from coordinated expression of known target genes. Consistent with the pathway analysis, SLC6A8-deficient leukemia cells exposed to AZA+T cells showed increased inferred activity of multiple TFs belonging to the IRF, STAT, and NF-κB families (**Fig. S8A-B**).

**Figure 5.**
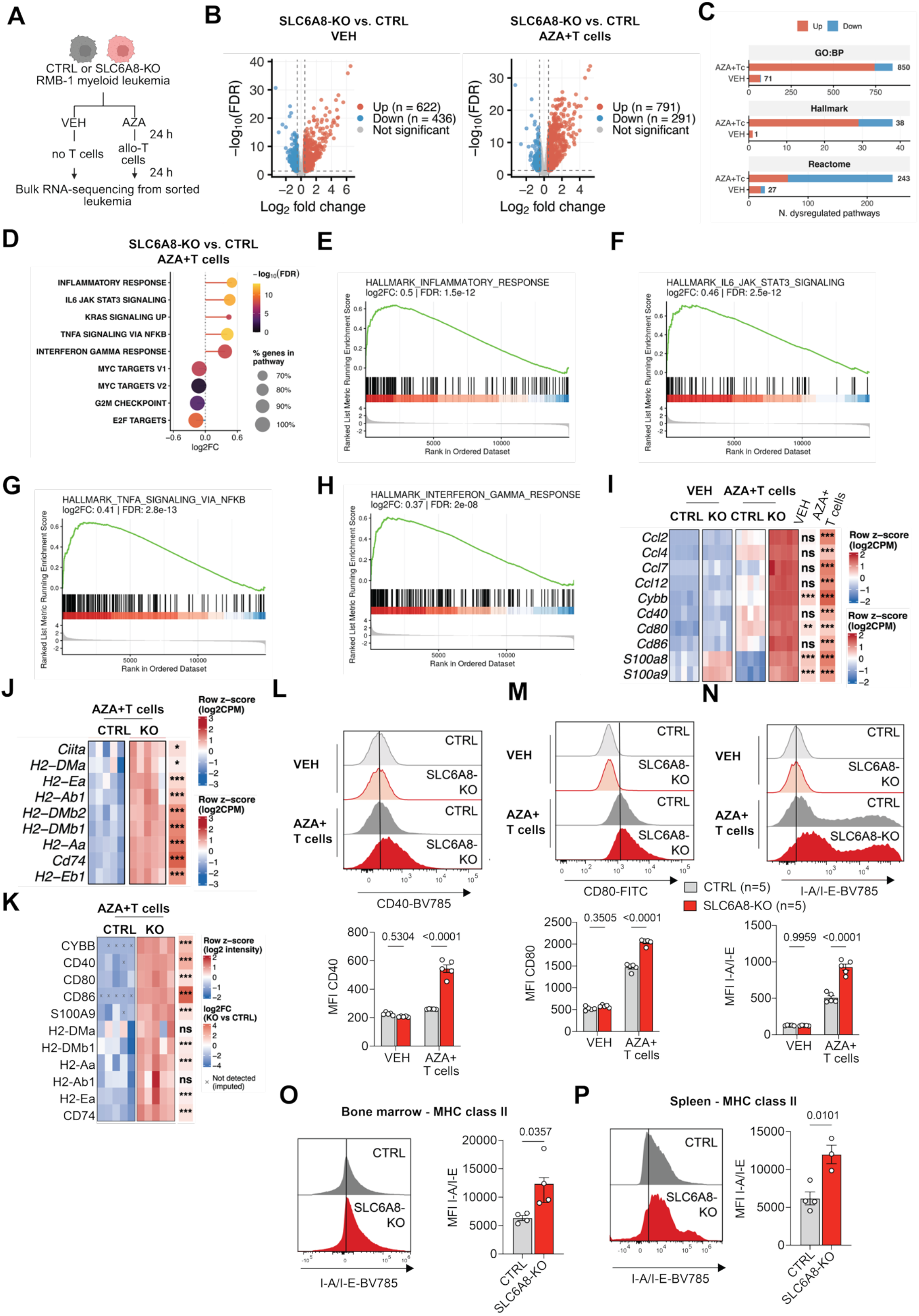
SLC6A8 deficiency enhances inflammatory and antigen-presentation programs under chemo-immune pressure. (**A**) SLC6A8-KO and CTRL RMB-1 were kept without treatment (VEH) or exposed sequentially to azacytidine (AZA, 2.5 µM) and activated allogeneic C57BL/6 CD3^+^ T cells (1:1 AML:T cell ratio) for 24 h each. Surviving leukemia cells were sorted by flow cytometry and submitted to transcriptomics analysis (B-J) or global proteomics (K), n=5/group. **(B)** Volcano plots of differential gene expression of SLC6A8-KO effect in (left) RMB-1 cells alone and (right) RMB-1 cells exposed to AZA+T cells. Genes meeting significance thresholds (FDR < 0.05, |log2FC| ≥ 0.5) are colored red (up-regulated in SLC6A8-KO) or blue (down-regulated in SLC6A8-KO); all other genes are shown in grey. Dashed lines indicate the significance thresholds. **(C)** Number of significantly dysregulated pathways from the indicated MSigDB collections for the SLC6A8 KO vs. CTRL comparisons, shown separately for VEH and AZA+T cells conditions. **(D)** Top 5 enriched and depleted Hallmark gene sets for KO vs CTRL comparison, ranked by log_2_FC. Dot size indicates the fraction of genes in the gene set present in the list of measured genes. **(E)-(H)** Enrichment plot of the indicated Hallmark gene sets for the SLC6A8-KO vs CTRL comparison under AZA+T cell pressure, identified by competitive gene set testing. **(I)** Expression heatmap of selected chemokine, myeloid and co-stimulatory genes across genotype and treatment groups. Values are log_2_ counts per million, scaled per gene (row z-score). Right annotation: log_2_ fold change of SLC6A8-KO versus CTRL cells under VEH and AZA+T cell conditions. Asterisks indicate the BH-adjusted p-value (* p < 0.05, ** p < 0.01, *** p < 0.001; ns, not significant). **(J)** Expression heatmap of selected genes related to MHC-II antigen presentation across the AZA+T cells groups. Values are log_2_ counts per million, scaled per gene (row z-score). Right annotation: log_2_ fold change of SLC6A8-KO versus CTRL cells under AZA+T cell conditions. Asterisks indicate the BH-adjusted p-value (* p < 0.05, ** p < 0.01, *** p < 0.001; ns, not significant). **(K)** Expression heatmap of selected proteins related to antigen presentation and T cell co-stimulation across the AZA+T cell groups. Values are log_2_ intensity, scaled per protein (row z-score). Crosses indicate imputed intensity, their values corresponding to imputation floor. Right annotation: log_2_ fold change of SLC6A8-KO versus CTRL cells under AZA+T cell conditions. Asterisks indicate the BH-adjusted p-value (* p < 0.05, ** p < 0.01, *** p < 0.001; ns, not significant). **(L)-(N)** Flow cytometry assessment of cell surface expression of CD40 (L), CD80 (M) and MHC-II (I-A/I-E, N) on SLC6A8-KO and CTRL RMB-1 cells cultured under the indicated conditions. Upper panels: representative flow cytometry histograms. Lower panels: quantification. Data are mean±SEM, n= 5/group. **(O)-(P)** Flow cytometry assessment of MHC class II (I-A/I-E) expression on mCherry^+^ RMB-1 AML cells in the BM (O) or spleen (P) of mice engrafted with CTRL or SLC6A8-KO AML and allogeneic C57BL/6 BM and T cells according to the scheme in Fig. 3G. Left: representative flow cytometry histograms. Right: quantification. Data are mean±SEM, n=4/group. **Statistical analysis and data representation: (B)-(H)** Statistical analysis of n=5 replicates/group as described in Methods. **(L)-(N)** Statistical analysis with Two-Way ANOVA with correction for multiple comparisons. Representative data from one of two independent experiments. **(O)-(P)** Statistical analysis with unpaired two-tailed t-test. Representative data from one of three independent experiments.

The transcriptional status of SLC6A8-KO leukemia exposed to chemo-immune pressure was further characterized by significantly increased expression of genes associated with immune activation, including the chemokines *Ccl2*, *Ccl7*, *Ccl12*, the co-stimulatory molecules *Cd40*, *Cd86*, *Cd80*, and the myeloid differentiation-associated alarmins *S100a8,* and *S100a9* (**Fig. 5I**). Because efficient T cell recognition additionally depends on antigen processing and presentation, we next examined this transcriptional program in greater detail. SLC6A8-deficient cells exposed to AZA+T cells showed significantly increased expression of multiple genes involved in antigen processing and presentation, including components of the major histocompatibility complex class II (MHC-II) pathway, such as *Cd74*, *H2-Aa*, *H2-Ab1*, *H2-Ea*, and the master regulator of MHC-II *Ciita* (**Fig. 5J**). These transcriptional signatures were reflected in altered protein abundance as determined by global proteomics (**Fig. 5K**). To investigate whether this also translated into altered expression at the leukemia cell surface, we next assessed expression of co-stimulatory molecules by flow cytometry. SLC6A8-KO RMB-1 cells showed markedly increased cell surface expression of CD40 and CD80 compared with CTRL cells following AZA+T cell exposure (**Fig. 5L-M**). This result suggested that loss of SLC6A8 promotes acquisition of a more immunostimulatory phenotype under chemo-immune pressure. Furthermore, expression of MHC-II (I-A/I-E) was higher on the surface of SLC6A8-KO compared to CTRL leukemia under chemo-immune pressure *in vitro* (**Fig. 5N**) and *in vivo* (**Fig. 5O-P**). This difference was confirmed in HL-60 AML exposed to activated allogeneic T cells *in vitro* (**Fig. S9A**). Notably, expression of MHC-I did not differ consistently between SLC6A8-KO and CTRL leukemia (**Fig. S9B-D**). Thus, the inflammatory transcriptional state associated with SLC6A8 deficiency is accompanied by increased expression of co-stimulatory and antigen-presentation molecules, suggesting an increased capacity of SLC6A8-deficient AML to engage allogeneic T cell responses.

To determine whether the inflammatory transcriptional phenotype associated with SLC6A8 loss was driven predominantly by chemotherapy, immune pressure, or their combination, we expanded our experimental setup to investigate the transcriptional responses of SLC6A8-KO and CTRL RMB-1 leukemia to AZA alone, allo-T cells alone, or the combination (**Fig. S10A**). We found that SLC6A8-KO cells showed higher transcription of genes belonging to some of the inflammation-related gene sets than CTRL cells (e.g., antigen presentation under allo-T cell pressure or cytokine signaling under AZA), but the full difference was unmasked only under combined AZA+T cell treatment (**Fig. S10B**). TF activity inference showed that exposure to allo-T cells, and particularly to the combination of AZA+T cells, led to higher inferred activity of STAT/IRF/NFkB TF in SLC6A8-KO compared to CTRL leukemia (**Fig. S10C**).

Together, these data support a model in which SLC6A8 limits the capacity of AML cells to undergo inflammatory and antigen-presenting state remodeling in response to combined chemo-immune pressure. Loss of SLC6A8 promotes surface expression of co-stimulatory molecules and MHC-II, shifting leukemia cells toward a more immunostimulatory phenotype. We propose that this more immunogenic state contributes to the enhanced susceptibility of SLC6A8-deficient leukemia to donor T cell-mediated elimination, associated with reduced leukemia burden and improved survival.

### IFN-γ drives inflammatory and antigen-presenting state remodeling in SLC6A8-deficient AML

The prominent enrichment of IFN-γ-response programs in SLC6A8-deficient leukemia exposed to allogeneic T cells prompted us to ask whether IFN-γ was sufficient to induce this inflammatory state independently of direct T cell contact. We therefore treated AZA-pre-exposed CTRL and SLC6A8-KO RMB-1 cells with recombinant IFN-γ for 24 h and performed bulk RNA-sequencing (**Fig. 6A**). We found that the transcriptional differences induced by recombinant IFN-γ showed a high concordance with those observed in SLC6A8-KO versus CTRL cells exposed to AZA+T cells, indicating that a single T cell-derived cytokine is sufficient to reproduce a substantial component of the immune pressure-associated phenotype (**Fig. 6B**). SLC6A8-deficient cells showed increased enrichment of inflammatory response, allograft rejection, TNF/NF-κB, and IL6/JAK/STAT3-associated gene sets (**Fig. 6C**). This was accompanied by increased expression of *Cybb, Ccl2*, *Ccl7*, *Ccl12*, *Cd40*, *Cd80*, *Cd86, S100a8, S100a9* and components of the MHC-II pathway (**Fig. 6D-E**). Transcriptional differences were reflected at the protein level as we detected higher surface CD40 and MHC-II in SLC6A8-KO AML exposed to AZA+IFN-γ (**Fig. 6F-G**). Notably, this was primarily observed upon combined chemo-cytokine pressure as neither AZA nor IFN-γ alone could elicit the full phenotype for CD40 (**Fig. 6F-G**). Signaling blockade with an anti-IFNγR1 antibody reversed the increased surface expression of CD40 and MHC-II in SLC6A8-deficient AML cells to a significant extent, confirming the importance of canonical IFN signaling (**Fig. 6F-G**).

**Figure 6.**
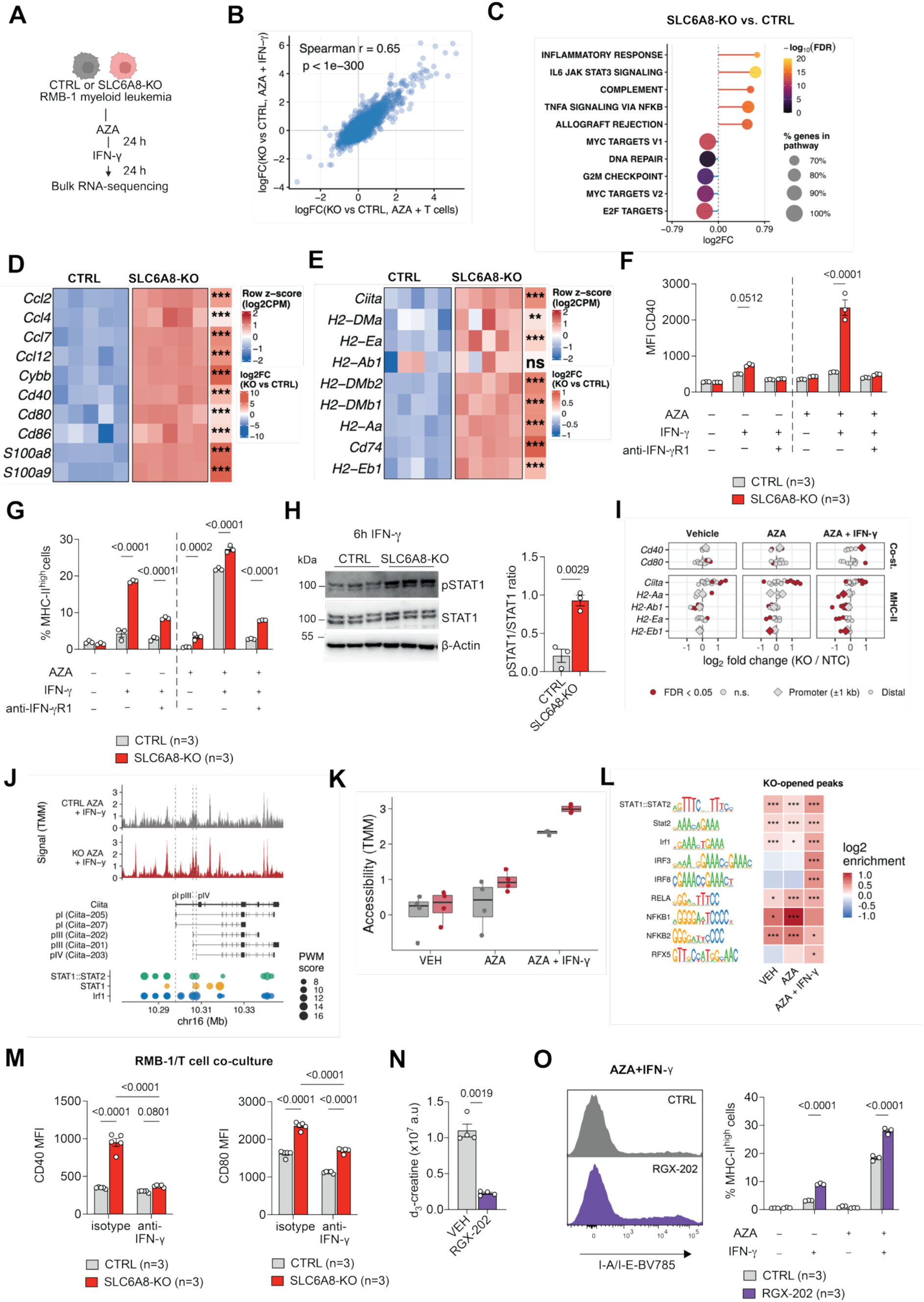
IFN-γ drives inflammatory and antigen-presenting state remodeling in SLC6A8-deficient AML. **(A)-(E)** SLC6A8-KO and CTRL RMB-1 leukemia cells were subjected for 24 h to 5 µM azacytidine (AZA) treatment, followed by 24 h with 10 ng/ml IFN-γ and bulk RNA-sequencing, n=5/group. **(A)** Experimental approach. **(B)** Correlation of the transcriptional response of AZA-exposed SLC6A8-KO to allogeneic T cells and IFN-γ, respectively. Spearman’s rank correlation coefficient and the associated p-value are indicated. **(C)** Top five enriched or depleted Hallmark gene sets for SLC6A8-KO vs. CTRL leukemia cells (FDR-wise, FDR<0.05), ranked by log_2_FC. Dot size indicates the fraction of genes in the gene set present in the list of measured genes. **(D)-(E)** Expression heatmap of selected chemokine, myeloid and co-stimulatory genes (D) and antigen presentation-associated genes (E) across genotype and treatment groups. Values are log_2_ counts per million, scaled per gene (row z-score). Right annotation: log_2_ fold change of SLC6A8-KO versus CTRL cells under AZA+IFN-γ conditions. Asterisks indicate the BH-adjusted p-value (* p < 0.05, ** p < 0.01, *** p < 0.001; ns, not significant). **(F)** SLC6A8-KO and CTRL RMB-1 leukemia cells were subjected for 48 h to the indicated treatments (5 µM AZA, 10 ng/ml IFN-γ) and cell surface expression of CD40 was measured by flow cytometry. Data are mean±SEM, n=3-4/group. **(G)** SLC6A8-KO and CTRL RMB-1 leukemia cells were subjected for 24 h to the indicated treatments (5 µM AZA, 10 ng/ml IFN-γ) and cell surface expression of MHC-II was measured by flow cytometry. Data are mean±SEM, n=3-4/group. **(H)** Western blot for pSTAT1, STAT1 and β-actin in CTRL or SLC6A8-KO RMB-1 cells treated for 24 h with AZA, followed by 6 h of 10 ng/ml IFN-γ. Right panel: densitometry quantification of pSTAT1/STAT1 ratio. Data are mean±SEM, n=3-4/group. **(I)-(L)** ATAC sequencing was performed from SLC6A8-KO and CTRL RMB-1 leukemia cells subjected for 24 h to vehicle (VEH) or 5 µM AZA treatment, followed by 24 h with 10 ng/ml IFN-γ in the indicated groups, n=3-4/group. **(I)** Chromatin accessibility difference of costimulatory and MHC-II loci in SLC6A8-KO vs CTRL RMB-1 cells. Each point is one consensus ATAC-seq peak within ±25 kb of the indicated gene, assigned to the gene with the nearest transcription start site. The x-axis shows the log_2_ fold change in accessibility (KO vs CTRL) within each treatment. Diamonds represent promoter peaks (TSS ± 1 kb), circles represent distal peaks. **(J)** Chromatin accessibility at the *Ciita* locus in SLC6A8-KO and CTRL RMB-1 cells under AZA+IFN-γ. Top: normalized (TMM) accessibility, shown as fragment coverage in 50-bp bins and averaged across biological replicates (CTRL, n=3; KO, n=4). Middle: genome model from GENCODE vM35, showing the gene body and the alternative promoters defined as transcription start sites (dashed lines) of isoform splicing. Bottom: predicted transcription factor binding sites within ATAC-seq consensus peaks with scores indicated as point size. **(K)** Normalized (TMM) accessibility of the *Ciita* pIV promoter in SLC6A8-KO and CTRL cells subjected to the indicated treatments. **(L)** Transcription factor motif enrichment in chromatin regions opened by SLC6A8-KO among treatments. Enrichment of JASPAR 2024 CORE vertebrate motifs with significantly increased accessibility in SLC6A8-KO vs CTRL. Sequence logos show the binding preference of each motif. Color indicates log_2_ enrichment of KO-opened peaks against all other peaks, and significance (BH-adjusted p-value) by asterisks; * adjusted p < 0.05, ** < 0.01, *** < 0.001. **(M)** SLC6A8-KO and CTRL RMB-1 cells were co-cultured with activated allogeneic T cells in a 1:1 ratio in the presence of 10 µg/ml anti-IFN-γ antibody or an isotype control. Cell surface expression of CD40 and CD80 was measured by flow cytometry. Data are mean±SEM, n=5/group. **(N)** RMB-1 leukemia cells were treated with vehicle or 10 mM RGX-202 for 24 h, followed by 24 h labeling with d_3_-creatine. Intracellular d_3_-creatine abundance by LC-MS. Data are mean±SEM, n=5/group. **(O)** RMB-1 leukemia cells were treated with VEH or 5 µM AZA ± 10 mM RGX-202 for 24 h. VEH or AZA was removed and cells were exposed to 10 ng/ml IFN-γ ± 10 mM RGX-202 for another 24 h. Expression of MHC-II by flow cytometry. Data are mean±SEM, n=3/group. **Statistical analysis and data representation: (B)-(C)** Statistical analysis of n=5 replicates/group as described in Methods. **(F)-(G)** Statistical analysis with Two-Way ANOVA with correction for multiple comparisons. Representative data from one of two independent experiments are shown. **(H)** Statistical analysis with unpaired two-tailed t-test. **(I)-(L)** Statistical analysis of n=3-4 replicates/group as described in Methods. **(L)** Statistical analysis with one-sided Fisher’s exact test, Benjamini–Hochberg-adjusted across a curated set of 37 motifs. **(M)** Statistical analysis with Two-Way ANOVA with correction for multiple comparisons. **(N)** Statistical analysis with unpaired two-tailed t-test. **(O)** Statistical analysis with Two-Way ANOVA with correction for multiple comparisons. Representative data from two independent experiments.

To understand the mechanism behind the enhanced IFN-γ response of RMB-1 cells, we first examined total cellular abundance of IFNGR1, the α chain of the IFN-γ receptor, and found no significant differences between CTRL and SLC6A8-KO cells (**Fig. S11A**). We next assessed signaling downstream of the IFN-γ receptor. STAT1 phosphorylation was increased in SLC6A8-KO compared with CTRL cells at 6 h after IFN-γ stimulation, indicating enhanced STAT1 activation in the absence of SLC6A8 (**Fig. 6H**). Because the magnitude of IFN-γ-driven transcriptional responses is also shaped by the accessibility of their regulatory elements, we next asked whether SLC6A8 loss altered chromatin accessibility at loci associated with the inflammatory and antigen-presenting state. ATAC-seq showed increased accessibility at regulatory elements associated with *Cd40, Cd80,* and *Ciita* in SLC6A8-deficient AML, whereas classical MHC-II loci (*H2-Aa*, *H2-Ab1*, *H2-Ea*, *H2-Eb1*) had comparatively limited differential accessibility, suggesting that enhanced MHC-II expression may be mediated predominantly through altered *Ciita* regulation (**Fig. 6I**). Notably, these accessibility differences at *Cd40* and *Ciita* emerged predominantly following combined AZA+IFN-γ exposure and were not evident at steady state or after AZA alone (**Fig. 6I-K, Fig. S11B**). At the *Ciita* locus, SLC6A8-deficient cells showed increased accessibility at a peak overlapping the IFN-γ-responsive *Ciita* promoter IV (pIV) following AZA+IFN-γ exposure (**Fig. 6J-K**). Motif analysis of this differentially accessible pIV peak identified predicted STAT– and IRF-family transcription factor binding motifs within the accessible region (**Fig. 6J**). Overall, regions gaining accessibility in SLC6A8-deficient cells were enriched for motifs recognized by inflammatory transcription factors, such as STAT, IRF, and NFκB-family motifs (**Fig. 6L**). Together, these findings suggest that loss of SLC6A8 enhances IFN-γ-induced STAT1 phosphorylation and the chromatin responsiveness of immune-regulatory programs, thereby promoting the expression of antigen-presenting and co-stimulatory molecules on the leukemia cell surface.

To determine to which extent IFN-γ also contributes to the phenotype during allo-T cell exposure, we neutralized IFN-γ during AZA+T cell coculture. IFN-γ blockade substantially reduced the induction of CD40 and CD80 in SLC6A8-KO leukemia cells and substantially diminished the difference between SLC6A8-KO and CTRL cells (**Fig. 6M**). These findings identify IFN-γ as a major mediator of the inflammatory and antigen-presenting state acquired by SLC6A8-deficient AML under allogeneic immune pressure.

Finally, we applied RGX-202 (3-guanidinopropionic acid), a creatine analog which acts as a pharmacological inhibitor of SLC6A8 by competing with creatine for the transporter^44^. We observed that RGX-202 substantially reduced the uptake of d_3_-creatine (**Fig. 6N**). In the presence of IFN-γ±AZA, RGX-202 substantially enhanced the expression of MHC-II, phenocopying the effects of genetic SLC6A8 deletion (**Fig. 6O**). Collectively, these findings identify SLC6A8 as a leukemia-intrinsic regulator that restrains IFN-γ-driven signaling and chromatin remodeling, thereby limiting acquisition of an inflammatory and antigen-presenting state.

### SLC6A8 expression in human AML is associated with erythroid differentiation state

Having identified SLC6A8 induction as an adaptive response to chemotherapy, we next asked whether baseline *SLC6A8* expression also varies across human AML and whether this variation is associated with specific disease or differentiation states, potentially identifying patients who may experience particular benefit from SLC6A8 blocking therapy. Using bulk transcriptomics from the BeatAML2.0 cohort^45^, we found that *SLC6A8* expression showed substantial inter-patient heterogeneity and was overall higher in AML than in healthy hematopoietic CD34^+^ cells (**Fig. 7A**). To identify transcriptional states associated with *SLC6A8* expression, we first sought genes whose expression covaried with *SLC6A8* across the BeatAML2.0 cohort, among samples with high blast content (>80%). This threshold was selected to minimize confounding introduced by different cell type composition of the samples, which consisted of blood or BM mononuclear cells. Because *SLC6A8* expression was itself associated with several clinical and technical variables, including prior MDS and *FLT3*-ITD status (**Fig. 7F-G**), simple gene-gene correlation would conflate *SLC6A8*-specific associations with these shared sources of variation. We therefore modeled *SLC6A8* abundance as a continuous predictor of genome-wide gene expression, which estimate, for each gene, the association with *SLC6A8* while adjusting for driver mutations status and technical effects. This framework can be viewed as a covariate-adjusted correlation analysis that additionally accounts for the mean-variance relationship commonly described in RNA-seq count data. Gene-wise associations with *SLC6A8* expression were assessed by linear modeling (*limma* model), adjusting for covariates (mutation status, cohort wave, etc). Strikingly, genes most strongly positively associated with *SLC6A8* were enriched for erythroid differentiation and heme metabolism, including *ALAS2*, *HBB*, *HBA1*, *HBA2* and *SLC4A1*. This association was independently observed in both BeatAML2.0 and TCGA-LAML^46^ datasets with n=191 and n=52 evaluable AML samples, respectively (**Fig. 7B** and **Fig. S12A**). Consistently, GSEA identified strong positive associations between *SLC6A8* expression and heme-related transcriptional programs in both cohorts (**Fig. 7C** and **Fig. S12B**). To quantify this relationship at the level of individual samples, we generated a consensus erythroid score from multiple erythroid-associated gene sets (see Methods). *SLC6A8* expression correlated strongly with this score in both BeatAML2.0 (r=0.51, p<1e-14, **Fig. 7D**) and TCGA-LAML (r=0.75, p<1e-10, **Fig. S12C**). Because bulk RNA-sequencing profiles reflect both leukemia-intrinsic transcriptional states and differences in cellular composition, we asked whether the association between *SLC6A8* expression and erythroid transcriptional programs could be explained by variable erythroid cell abundance, even after we had selected samples with high blast content. Importantly, adjustment for inferred erythroid content using CIBERSORTx^47^ did not abolish this association (**Fig. S12D-E**), arguing against high erythroid cell content as the sole explanation.

**Figure 7.**
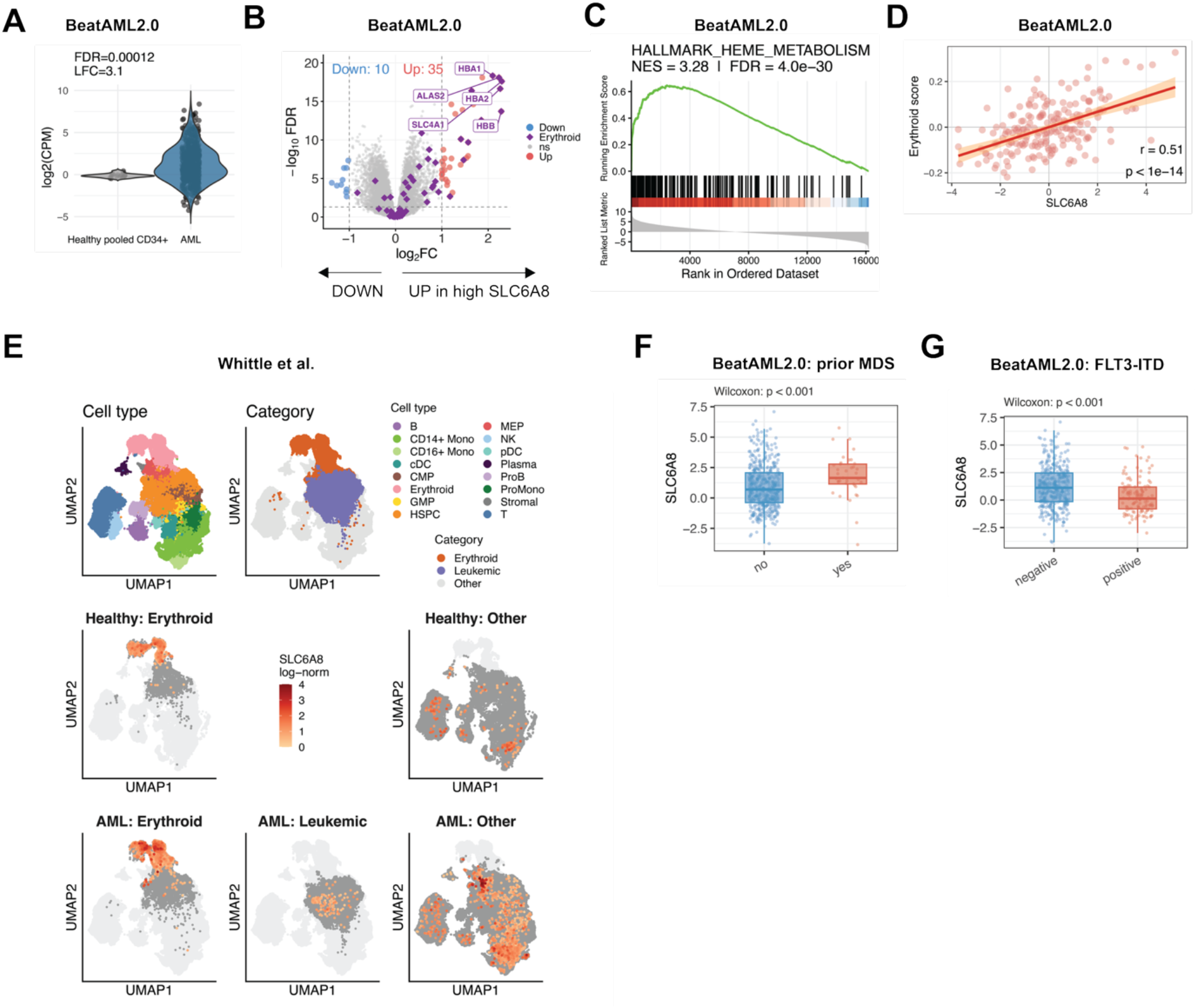
High SLC6A8 mRNA correlates with prior MDS and erythroid transcriptional signatures. (**A**) *SLC6A8* mRNA (log2CPM) normalized expression in healthy CD34^+^ cells and mononuclear cells from AML patients from the BeatAML2.0 cohort. **(B)** Volcano plot depicting genes correlated or anti-correlated with high *SLC6A8* expression in the BeatAML2.0 cohort, highlighting erythroid markers genes recovered from public databases (PanglaoDB, CellMarker 2.0). **(C)** Gene set enrichment analysis for the correlation between high *SLC6A8* expression and the Hallmark Heme metabolism gene set. **(D)** Correlation between *SLC6A8* mRNA and erythroid score in the BeatAML2.0 cohort. A linear fit is depicted to represent the trend. **(E)** UMAP of a single-cell AML atlas^48^ highlighting *SLC6A8* expression from primary AML samples and healthy donors. The UMAP are split by donor group (rows) and cell populations categories (columns). Dot colors depict the cell populations and the (non-) expression of *SLC6A8*; light grey, all cells of the donor group; dark grey, highlighted cells without detectable *SLC6A8*; coloured, *SLC6A8-*expressing cells. **(F)-(G)** *SLC6A8* expression in AML patients stratified according to (F) the occurrence of a prior MDS and (G) the *FLT3*-ITD mutational status, based on the BeatAML2.0 cohort. **Statistical analysis and data representation: (A)** Statistical analysis on 16 healthy vs 649 AML patient samples using linear modeling. **(B)-(D)** Statistical analysis on 191 samples testing transcriptomics-wise association with SLC6A8 via linear modeling, as described in Methods. **(E)** Descriptive representation, each dot showing one cell from the publicly available AML atlas (n=748,679 cells from 159 AML patients and 51 healthy donors), positioned using the authors’ integrated UMAP, also described in Methods. **(F)-(G)** Statistical analysis of 483 samples using two-sample Wilcoxon test.

To place these findings in the context of healthy and malignant hematopoiesis, we interrogated a published single-cell RNA-sequencing atlas of BM from healthy donors and AML patients^48^. *SLC6A8* was physiologically enriched in erythroid-lineage cells and was also detectable in a subset of malignant AML cells (**Fig. 7E**). Together, these data indicate that *SLC6A8* expression is linked to an erythroid-associated differentiation program that exists physiologically and is retained or adopted by subsets of AML. We next asked whether *SLC6A8* expression was associated with established clinical or molecular AML features. *SLC6A8* was not consistently associated with ELN 2017 risk category, common driver fusions, or *TP53* mutational status (**Fig. S12F-H**). In contrast, *SLC6A8* expression was higher in AML arising after a preceding myelodysplastic syndrome and lower in *FLT3*-ITD-mutant AML (**Fig. 7F-G**). Together, these findings place SLC6A8 at the intersection of adaptive and intrinsic AML biology: while chemotherapy induces *SLC6A8* as an acquired metabolic adaptation that promotes immune escape, baseline *SLC6A8* expression is shaped by leukemia cell state and differentiation programs in human AML.

## Discussion

In this study, we identify creatine accumulation via SLC6A8 as a conserved metabolic adaptation in AML cells surviving chemotherapy across multiple models and treatments. Our metabolomic profiling revealed that creatine and glutathione were universally elevated in chemotherapy-persisting AML. We prioritized creatine for mechanistic investigation as glutathione’s role in buffering ROS, detoxifying chemotherapy agents, and promoting therapy resistance is well-established across multiple malignancies^20,30,31^. In contrast, the functional significance of creatine accumulation in therapy-persisting leukemia cells was unknown, and the observation that creatine could be acquired through a single, druggable transporter (SLC6A8) presented an attractive opportunity to define a novel and potentially targetable pathway.

The consistency of creatine accumulation across diverse chemotherapy agents and AML models as well as in datasets from previous studies we explored^16,19^ suggests this represents a fundamental adaptive response rather than a drug-specific effect. The persistence of elevated intracellular creatine after drug withdrawal, and its progressive accumulation with serial treatment cycles, indicates that this metabolic state contributes to long-term adaptation rather than merely reflecting acute stress. Supporting clinical relevance, higher creatine levels were observed in relapsed/refractory AML patient samples compared to treatment-naïve disease^16^, and SLC6A8 expression correlated with prior myelodysplastic syndrome, a condition often treated with hypomethylating agents.

Using stable-isotope tracing and CRISPR/Cas9 genetic knockouts, we established that creatine accumulation occurs primarily through SLC6A8-mediated uptake rather than de novo synthesis. Notably, this finding may be model-specific, as a recent report showed that *FLT3*-ITD-mutant AML may rely more on de novo creatine synthesis, which was identified as a vulnerability in this subtype^49^. This is in line with our finding that *SLC6A8* expression was lower in patients with *FLT3*-ITD-mutant compared to *FLT3* unmutated AML, suggesting that therapeutic strategies targeting creatine metabolism may need to be tailored based on whether tumors depend on uptake versus synthesis.

Creatine metabolism has been implicated by multiple studies as an adaptation of tumors to potentially hostile environments. In colorectal cancer, metastatic cells exploit extracellular creatine/phosphocreatine metabolism to survive energetic stress during liver colonization^50^. SLC6A8 was required for uptake of extracellular phosphocreatine and metastatic outgrowth^50^. Pharmacological SLC6A8 inhibition with RGX-202 showed anti-tumor activity across colorectal cancer models limiting both primary tumor growth and metastasis^44^ and is currently in clinical trials. SLC6A8-dependent creatine uptake similarly supports prostate cancer progression^51^ and enables glioblastoma cells to survive within hypoxic niches, where tumor-associated myeloid cells provide an extracellular source of creatine^52^. Creatine accumulation was also observed in liver foci formed in a mouse AML model with C1498 cells, supporting ATP production to maintain energy homeostasis during infiltration, although the implications of this finding for the tumor-immune crosstalk were not investigated^53^. Importantly, creatine may support cancer growth and metastasis through both phosphocreatine-dependent and – independent mechanisms. While RGX-202 was implied to induce apoptosis by depleting intracellular phosphocreatine and ATP^44^, another study found that creatine enhances colorectal and breast cancer metastasis by upregulation of Snail and Slug expression via monopolar spindle 1 (MPS1)-activated Smad2 and Smad3 phosphorylation, independently of phosphocreatine generation^54^. In triple-negative breast cancer (TNBC), SLC6A8 was transcriptionally upregulated by p65/NF-κB and mediated creatine uptake which was essential for sustaining redox homeostasis under hypoxia and suppressing apoptosis^55^. At the same time, the effects of creatine appear to be highly context dependent. A recent study found that creatine can potently induce ferroptosis in colorectal cancer cells^56^. Creatine directly bound and inhibited extracellular signal-regulated kinase 2 (ERK2), preventing it from stabilizing the ferroptosis inhibitor ferroptosis suppressor protein 1 (FSP1)^56^.

In our study, while SLC6A8 had no effect on proliferation, chemotherapy sensitivity, or migration, its loss dramatically enhanced leukemia eradication under allogeneic T cell pressure across three independent models, demonstrating that SLC6A8 specifically promotes immune evasion rather than leukemia cell survival in the absence of immune pressure. Effector T cells and tumor-infiltrating lymphocytes depend on creatine uptake to sustain cytotoxic function^42,43^ and one of our hypotheses was that chemotherapy-persisting leukemia cells may deplete creatine from the leukemic environment, thus impairing T cell function. Dietary creatine supplementation worsened rather than improved survival in two mouse AML models, arguing against the hypothesis that increasing systemic creatine availability enhances leukemia immune control. Although these findings are compatible with a leukemia-intrinsic function of creatine uptake, the effects of supplementation on individual leukemia and immune cell populations remain to be defined.

The canonical metabolic role of creatine is to support cellular energy buffering through its reversible phosphorylation to phosphocreatine by creatine kinases^38,41^. Importantly, CKB deletion abolished phosphocreatine generation in RMB-1 leukemia without affecting immune-mediated clearance, indicating that the immune-regulatory function of SLC6A8-mediated creatine uptake is independent of canonical phosphocreatine-dependent energy buffering. Notably, the function of creatine in our study is distinct from what has previously been reported for EVI1-positive AML^57^. There, the authors discovered that oncogenic signaling through EVI1 (also known as MECOM) induced the expression of CKMT1, and targeting CKMT1 reduced the viability and promoted apoptosis of EVI1-positive cell lines *in vitro* and *in vivo* by disrupting mitochondrial respiration and ATP production^57^. Mechanistically, inhibition of the CK pathway in EVI1-positive cells blocked GSK3/WNT signaling^58^. With the development of covalent CK inhibitors, targeting this pathway may become feasible in a translational context^59^. Our data are distinct from these studies in several aspects. First, we observe SLC6A8-mediated regulation of inflammatory cell states in models with distinct genetic drivers. Second, this regulation is independent of the generation of phosphocreatine and cannot be targeted by inhibiting CK as demonstrated by the fact that CK deficiency did not phenocopy SLC6A8 deficiency.

Mechanistically, loss of SLC6A8 induced the expression of pro-inflammatory and antigen-presentation-related genes in leukemia cells under chemo-immune pressure. IFN-γ was sufficient to reproduce a substantial component of this signature and was a major mediator of the phenotype induced by allogeneic T cells as shown by experiments with direct exposure to or neutralization of IFN-γ. SLC6A8 regulated cytokine responsiveness by restricting signaling downstream of the IFN-γ receptor, as we found enhanced STAT1 phosphorylation in SLC6A8-KO cells exposed to IFN-γ. This finding is in line with a study by Ji et al. which demonstrated that creatine inhibited receptor proximal IFN-γ-JAK-STAT1 TF signaling to suppress pro-inflammatory macrophage remodeling^60^. In the same study, creatine enhanced IL-4-induced macrophage polarization by promoting chromatin remodeling, thus supporting IL-4-STAT6-driven arginase 1 expression^60^. We found chromatin remodeling in SLC6A8-KO cells compatible with higher accessibility at the *Cd40* and *Ciita* loci. While some differences in chromatin appeared to be present at steady-state, they were particularly evident under combined chemo– and cytokine pressure.

Changes in cellular redox state represent one potential link between creatine accumulation and inflammatory transcriptional responses. In hypoxic TNBC, SLC6A8-mediated creatine accumulation decreased mitochondrial activity and intracellular reactive oxygen species thereby promoting tumor cell survival by stabilizing redox homeostasis^55^. In a model of endometriosis, creatine supplementation reduced cellular iron concentrations, mitigating oxidative stress and lipid peroxidation^61^. The co-enrichment of creatine alongside glutathione, the cell’s principal antioxidant, as the only two metabolites universally elevated in chemotherapy-persisting AML suggests that both may serve complementary roles in buffering oxidative stress.

Notably, the effects of creatine on tumor immunity are strongly dependent on which cells use the creatine. SLC6A8-mediated creatine uptake and its phosphorylation by CKB supports the anti-tumor function of CD8^+^ T cells, and creatine supplementation enhanced tumor control and cooperated with PD-1/PD-L1 blockade in murine tumor models of melanoma^42,43^. For AML, extracellular vesicles released by NPM-1-mutant AML cells were shown to transfer miR-19a-3p to CD8^+^ T cells, suppressing SLC6A8 expression, creatine uptake and ATP production, thereby impairing T cell function^62^.

Our findings have translational implications. The SLC6A8 inhibitor RGX-202 is in clinical development, with preclinical data demonstrating anti-proliferative effects in solid tumors^44^. Our work suggests that in AML, the therapeutic benefit of SLC6A8 inhibition may derive not from direct cytotoxicity but from rendering leukemia cells more susceptible to immune attack. Because SLC6A8 is dispensable for leukemia growth in the absence of immune pressure, targeting this pathway could selectively sensitize persisting AML cells to immune-mediated eradication without impairing baseline leukemia cell fitness. This positions SLC6A8 blockade as a strategy to enhance GVL responses following allo-HSCT or to synergize with donor lymphocyte infusions. In addition, we found that leukemias with erythroid transcriptional programs have higher SLC6A8 expression, perhaps identifying a subset of patients that may benefit from SLC6A8 inhibition. More broadly, our work establishes a paradigm in which chemotherapy-induced metabolic adaptations not only promote therapy resistance but also actively reprogram tumor-immune interactions, suggesting that interrogating the metabolic states of therapy-persisting cells may reveal additional targetable vulnerabilities at the interface of cancer metabolism and immunology.

## Limitations of the study

The molecular link between SLC6A8 and enhanced STAT1 phosphorylation remains to be defined. While we demonstrated the importance of SLC6A8 in the allo-HSCT setting, its relevance to other immunotherapy contexts such as checkpoint inhibition, bispecific antibodies, or CAR-T cell therapy has not been tested. Furthermore, testing the *in vivo* effects of pharmacological inhibition of SLC6A8 with RGX-202 or more potent inhibitors will be important. The effects of SLC6A8 inhibition on normal hematopoietic stem cells, T cells, and other immune populations will require particularly careful evaluation, since creatine metabolism supports T cell function in other contexts^42,43^.

## Acknowledgements

We thank H. Christofk, D. Green, and M. Febbraio for helpful input, Michael Mitterer and Fabian Hässler for technical support, and Michal Stanczak for help with technical troubleshooting. The graphical abstract was created by Michal Rössler. ATAC-sequencing was performed at the Genomics Facility Basel, operated by the University of Basel and the Department of Biosystems Science and Engineering (D-BSSE), ETH Zurich. Calculations were performed at sciCORE (http://scicore.unibas.ch/) scientific computing center at University of Basel, with support by the SIB – Swiss Institute of Bioinformatics.

This study was funded by the Swiss National Science Foundation – project funding 10.001.317 to P.A., the Deutsche Forschungsgemeinschaft (DFG, German Research Foundation) – project 492259164 to P.A., the Novartis Foundation for Medical-Biological Research – project 24C246 to P.A., the Olga Mayenfisch foundation to P.A.. the Max Planck Society to E.L.P., NIH R01AI156274 to E.L.P, the Bloomberg∼Kimmel Institute, and a Johns Hopkins University Bloomberg Distinguished Professorship to E.L.P. P.A. further receives support by the European Union (ERC, LIFE, 101301617), the European Hematology Association (Bilateral Collaborative Research Grant 2024), the ISREC Foundation, the Jacques and Gloria Gossweiler Foundation, and the Deutsche Forschungsgemeinschaft (DFG, German Research Foundation) – CRC 1479, P05. Views and opinions expressed are however those of the author(s) only and do not necessarily reflect those of the European Union or the European Research Council. Neither the European Union nor the granting authority can be held responsible for them. L.P. is a participant in the German Society of Internal medicine (DGIM) Advanced Clinician Scientist Program (ASCP), and is supported by the American Society of Hematology (ASH) Global Research Award, the Max-Eder program of the German Cancer Aid (Deutsche Krebshilfe), the Else Kröner-Fresenius-Stiftung (2023_EKEA.102), the DKMS John Hansen Research Grant, research funding from the German José Carreras Leukemia Foundation (16 R/2026), and the Brigitte und Dr. Konstanze Wegener-Stiftung [Project #133 A 2024 II (7)].

---

**Figure S1.**
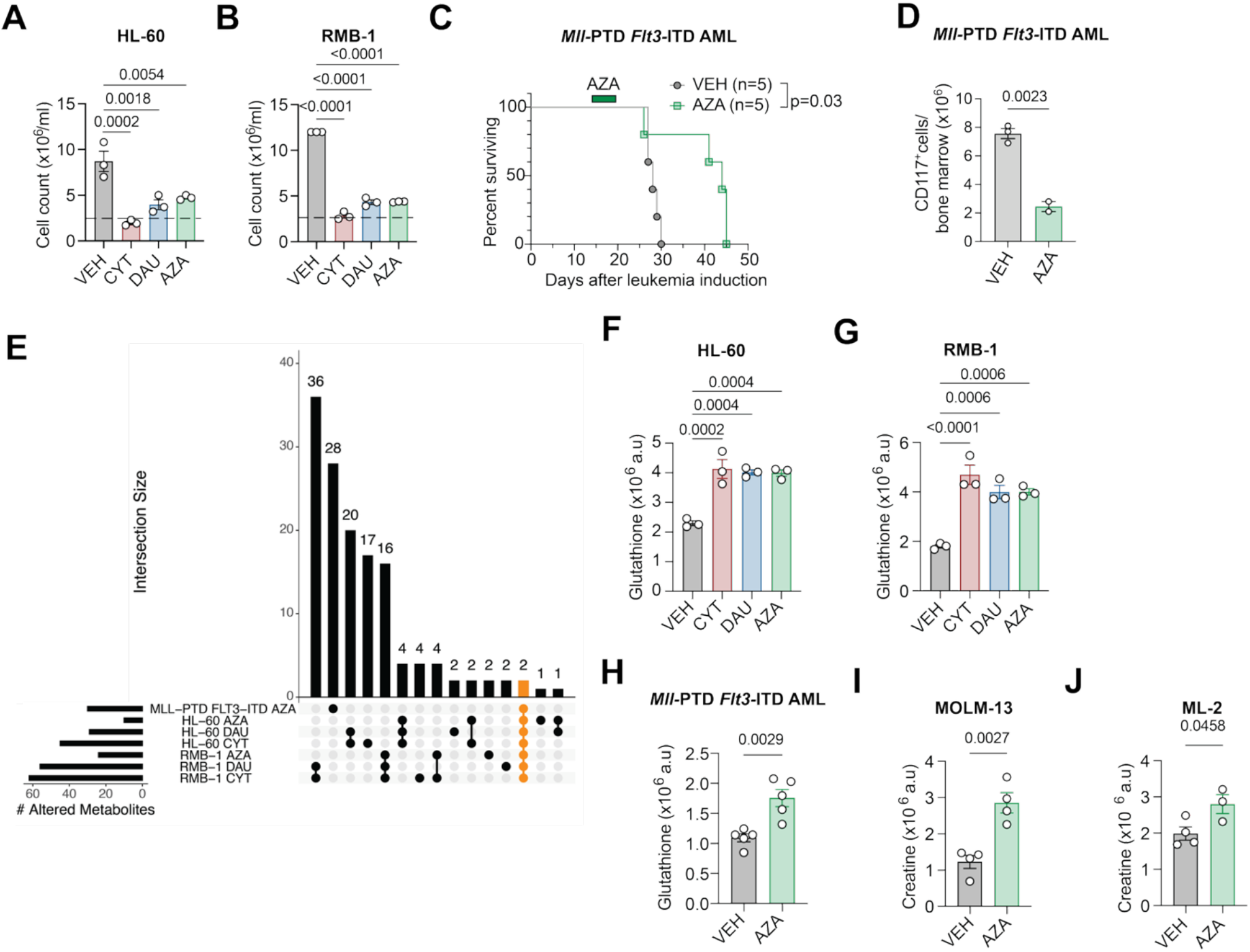
Chemotherapy selection pressure induces metabolic adaptations in AML. (**A**) HL-60 living cell numbers after 24 h exposure to the indicated treatments. Data are mean±SEM, n=3/group. **(B)** RMB-1 living cell numbers after 24 h exposure to the indicated treatments. Data are mean±SEM, n=3/group. **(C)** Survival of *Mll*-PTD *Flt3*-ITD AML-bearing mice treated with 5 mg/kg BW AZA or VEH at the indicated time point, n=5/group. **(D)** Number of CD117^+^ cells isolated from the BM of *Mll*-PTD *Flt3*-ITD AML-bearing mice treated as in (C). Data are mean±SEM, n=2-3/group. **(E)** Upset plot depicting the overlap of significantly regulated metabolites across treatments and models. For RMB-1 and HL-60 models, the values indicate the number of altered metabolites detected using a one-way ANOVA, and the differences between VEH and each condition, detected using post-hoc Tukey HSD tests. **(F)** Glutathione abundance by LC-MS in HL-60 cells exposed to the indicated treatments. Data are mean±SEM, n=3/group. **(G)** Glutathione abundance by LC-MS in RMB-1 cells exposed to the indicated treatments. Data are mean±SEM, n=3/group. **(H)** Glutathione abundance by LC-MS in CD117^+^ cells enriched from the BM of VEH– or AZA-treated *Mll-*PTD *Flt3*-ITD AML-bearing mice. Data are mean±SEM, n=5/group. **(I)-(J)** Creatine abundance by LC-MS in MOLM-13 and ML-2 cells treated with vehicle (VEH) or azacytidine (AZA) for 24 h. Data are mean±SEM, n=4/group. **Abbreviations:** VEH: vehicle, CYT: cytarabine, DAU: daunorubicin; AZA: azacytidine. **Statistical analysis and data representation: (A), (B)** Statistical analysis with One-way ANOVA followed by Dunnett’s multiple comparisons test. Representative data from one of three independent experiments. **(C)** Statistical analysis with Mantel-Cox test. **(D)** Statistical analysis with unpaired two-tailed t-test. Representative data from one of two independent experiments. **(F), (G)** Statistical analysis with One-way ANOVA followed by Dunnett’s multiple comparisons test. Representative data from one of three independent experiments. **(H)-(J)** Statistical analysis with unpaired two-tailed t-test. Representative data from one of two independent experiments.

**Figure S2.**
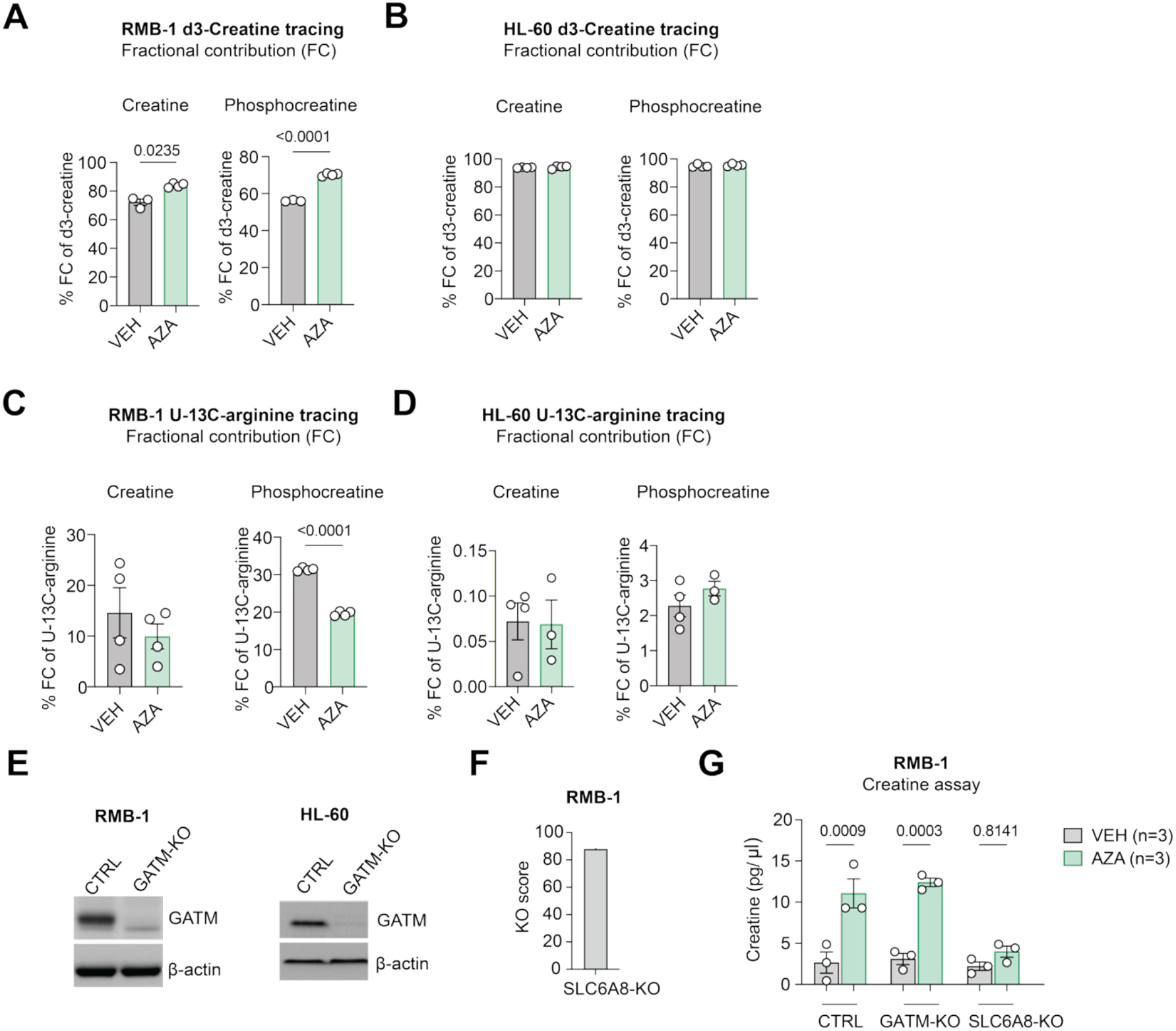
Uptake rather than de novo synthesis is the main source of intracellular creatine in AML. **(A)-(B)** d_3_-creatine tracing by LC-MS in RMB-1 (A) or HL-60 AML (B) pre-treated with vehicle (VEH) or azacytidine (AZA) for 24h. Labeling was performed for 24 h. Fractional contribution of d_3_-creatine to creatine and phosphocreatine. Data are mean±SEM, n=3-4/group. **(C)-(D)** U-^13^C-arginine tracing by LC-MS in RMB-1 (C) or HL-60 AML (D) pre-treated with vehicle (VEH) or azacytidine (AZA) for 24h. Labeling was performed for 24 h. Fractional contribution of U-^13^C-arginine to creatine and phosphocreatine. Data are mean±SEM, n=4/group. **(E)** GATM protein abundance in vehicle (VEH)– and azacytidine (AZA)-treated RMB-1 and HL-60 cells by western blotting with β-actin used as loading control. **(F)** KO score for SLC6A8 derived from inference of CRISPR edits analysis of gene-edited RMB-1 cells. **(G)** Creatine concentration by a colorimetric assay in control, GATM-KO and SLC6A8-KO RMB-1 cells treated with VEH or AZA for 24h. **Statistical analysis and data representation: (A)-(D)** Statistical analysis with unpaired two-tailed t-test (n=3-4/group). **(G)** Statistical analysis with One-Way ANOVA with correction for multiple comparisons.

**Figure S3.**
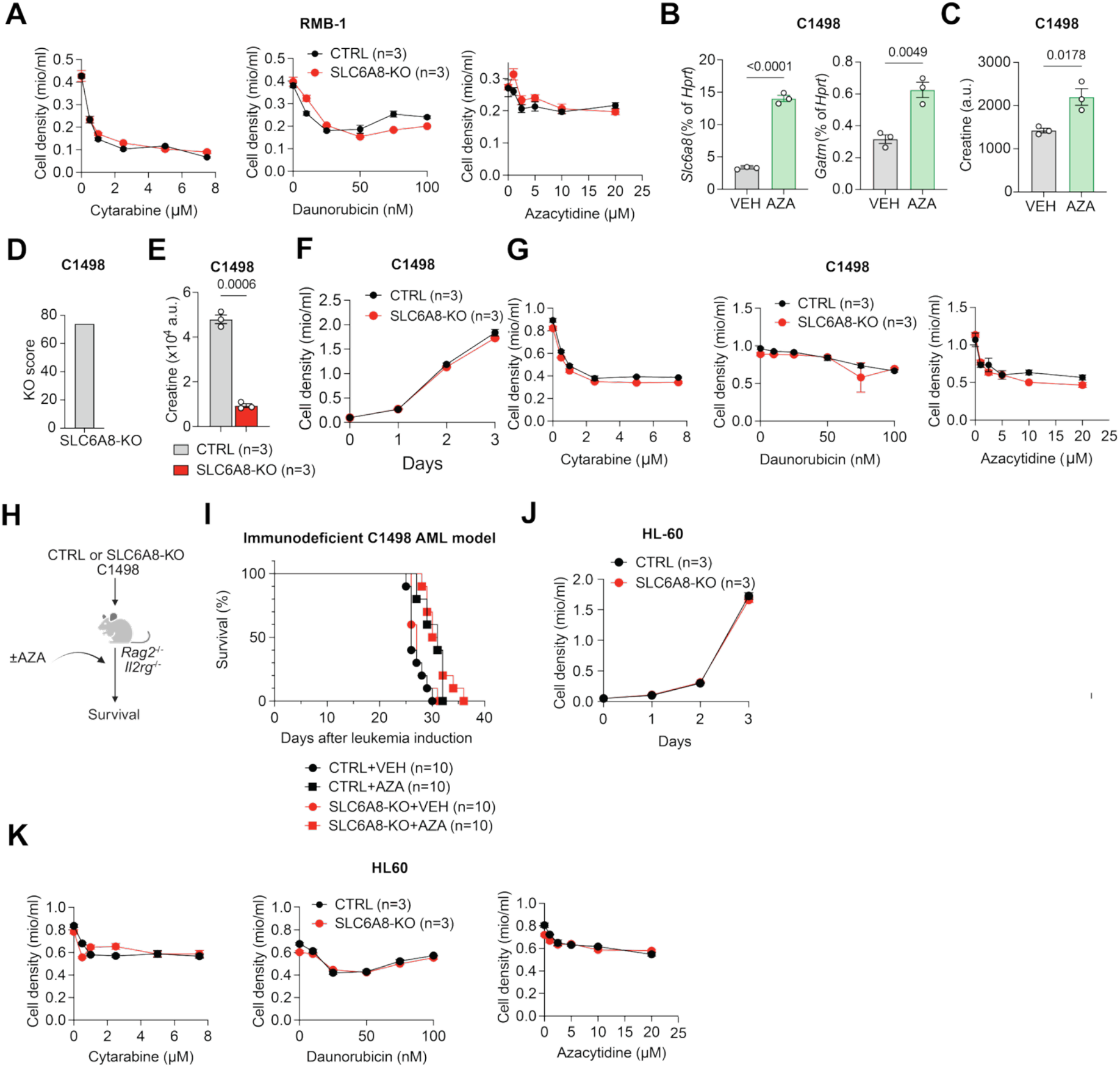
Knockout of SLC6A8 does not affect leukemia cell proliferation or chemotherapy sensitivity. (**A**) Cell density of CTRL and SLC6A8-KO RMB-1 cells treated with the indicated doses of cytarabine, daunorubicin or azacytidine (AZA) for 24 h. Data are mean±SEM, n=3/group. **(B)-(C)** *Gatm* and *Slc6a8* mRNA by qPCR **(B)** and creatine abundance by HPLC-MS/MS **(C)** in C1498 cells treated with vehicle (VEH) or AZA for 24 h. *Hprt* was used as housekeeping gene. Data are mean±SEM, n=3/group. **(D)** *Slc6a8* knockout score in C1498 AML cells calculated with inferene of CRISPR edits. **(E)** Creatine abundance in CTRL and SLC6A8-KO C1498 cells. Data are mean±SEM, n=3/group. **(F)** Cell density of CTRL and SLC6A8-KO C1498 cells over a time course of 3 days. Data are mean±SEM, n=3/group. **(G)** Cell density of CTRL and SLC6A8-KO C1498 cells treated with the indicated doses of cytarabine, daunorubicin or AZA for 24 h. Data are mean±SEM, n=3/group. **(H)** Experimental workflow for (I), created with Biorender.com. **(I)** Survival of immunodeficient *Rag2^-/-^ Il2rg^-/-^* mice engrafted with CTRL or SLC6A8-KO C1498 cells ± AZA treatment, n=10/group. **(J)** Cell density of CTRL and SLC6A8-KO HL-60 cells over a time course of 3 days. Data are mean±SEM, n=3/group. **(K)** Cell density of CTRL and SLC6A8-KO HL-60 cells treated with the indicated doses of cytarabine, daunorubicin, or AZA for 24 h. Data are mean±SEM, n=3/group. **Statistical analysis and data representation: (A), (F), (G), (J), (K)** Representative data from one of two to three independent experiments. **(B), (C), (E)** Statistical analysis with two-tailed unpaired t-test. Representative data from one of two independent experiments. **(I)** Statistical analysis with Mantel-Cox test. Pooled data from two independent experiments.

**Figure S4.**
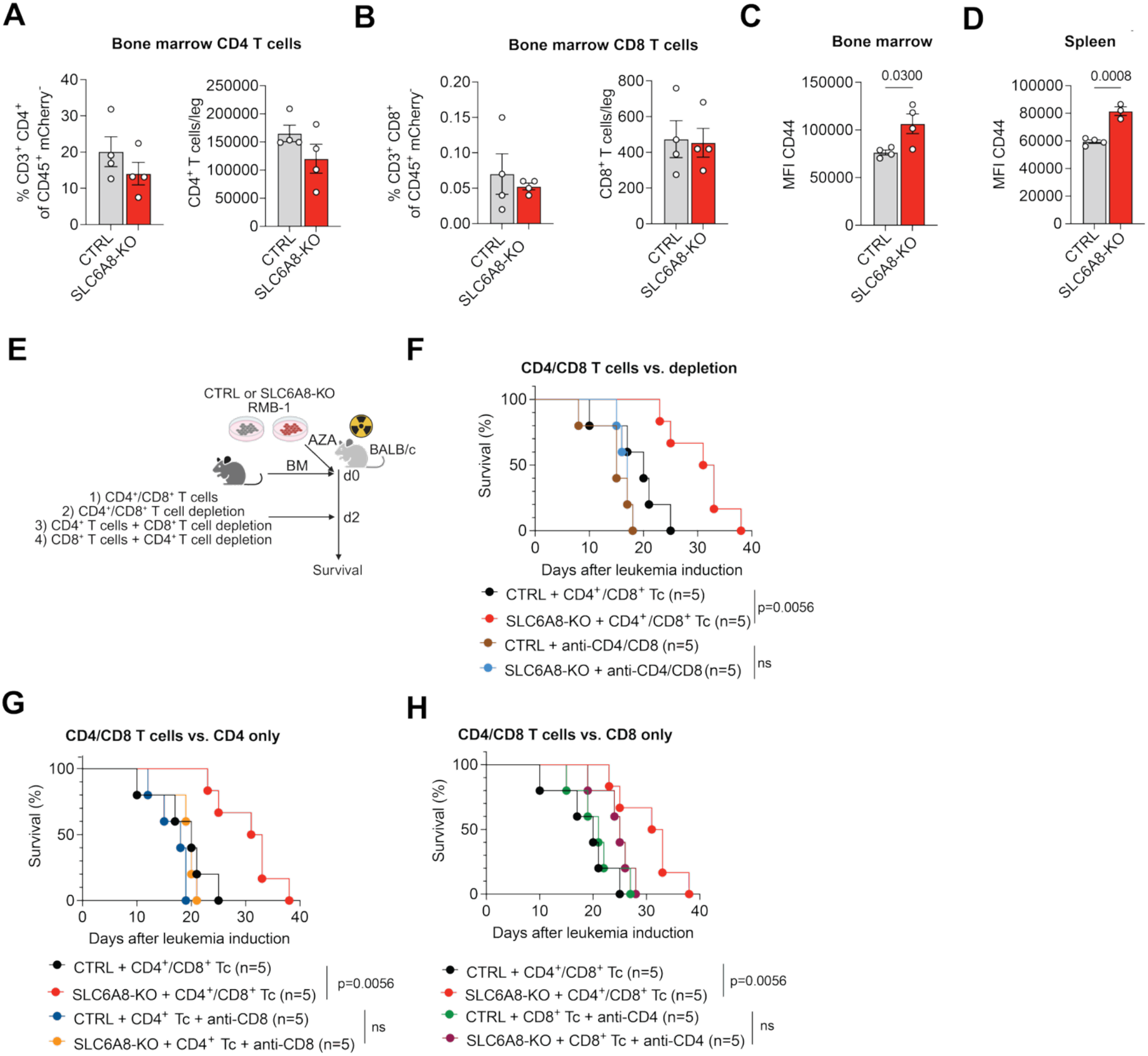
SLC6A8 loss sensitizes AML to coordinated CD4^+^ and CD8^+^ T cell pressure. (A)-(B) Percentage and absolute number of CD4^+^ (A) or CD8^+^ (B) T cells in the BM of mice engrafted with azacytidine (AZA)-treated SLC6A8-KO or CTRL RMB-1 and allogeneic T cells (as in Fig. 3G). Data are mean±SEM, n=4/group. **(C)-(D)** CD44 expression on CD3^+^ T cells from the BM or spleen. Data are mean±SEM, n=4/group. **(E)** Experimental workflow for (F)-(H), created with Biorender.com. **(F)** Survival of BALB/c mice engrafted with CTRL or SLC6A8-KO RMB-1 cells. Mice received allogeneic CD4^+^/CD8^+^ T cells and isotype control antibodies or no T cell transfer and anti-CD4^+^/anti-CD8^+^ depleting antibodies, n=5/group. **(G)** Survival of BALB/c mice engrafted with CTRL or SLC6A8-KO RMB-1 cells. Mice received allogeneic CD4^+^/CD8^+^ T cells and isotype control antibodies or CD4^+^ T cell transfer and anti-CD8^+^ depleting antibody, n=5/group. **(H)** Survival of BALB/c mice engrafted with CTRL or SLC6A8-KO RMB-1 cells. Mice received allogeneic CD4^+^/CD8^+^ T cells and isotype control antibodies or CD8^+^ T cell transfer and anti-CD4^+^ depleting antibody, n=5/group. **Statistical analysis and data representation: (A)-(D)** Statistical analysis with unpaired two-tailed t-test. Representative data from one of three independent experiments. **(F)-(H)** Statistical analysis with Mantel-Cox test. All groups were tested in the same experiment and the split in three panels is to improve readability. The CTRL+CD4^+^/CD8^+^ Tc and SLC6A8-KO + CD4^+^/CD8^+^ Tc groups are identical in the three graphs.

**Figure S5.**
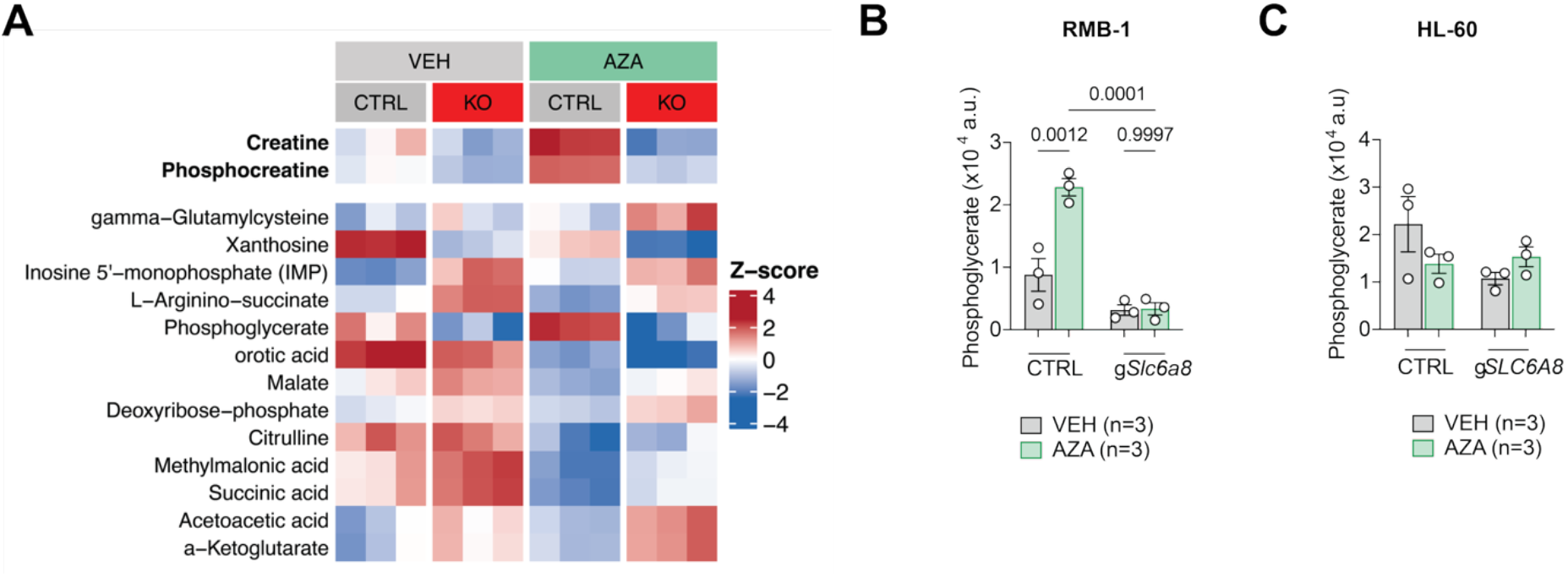
Metabolomics of SLC6A8-KO and CTRL leukemia support creatine uptake as the most likely function of SLC6A8. **(A)** Z-score depicting differentially abundant metabolites in SLC6A8-KO vs. CTRL RMB-1 cells ± azacytidine (AZA) treatment. Metabolites with FDR<0.05 for an effect of the knockout are displayed. **(B)-(C)** Phosphoglycerate abundance in SLC6A8-KO and CTRL RMB-1 (B) and HL-60 (C) AML ± AZA treatment. **Statistical analysis and data representation: (A)** Analysis of n=3 independent replicates as described in Methods. **(B)-(C)** Statistical analysis with One-Way ANOVA with correction for multiple comparisons.

**Figure S6.**
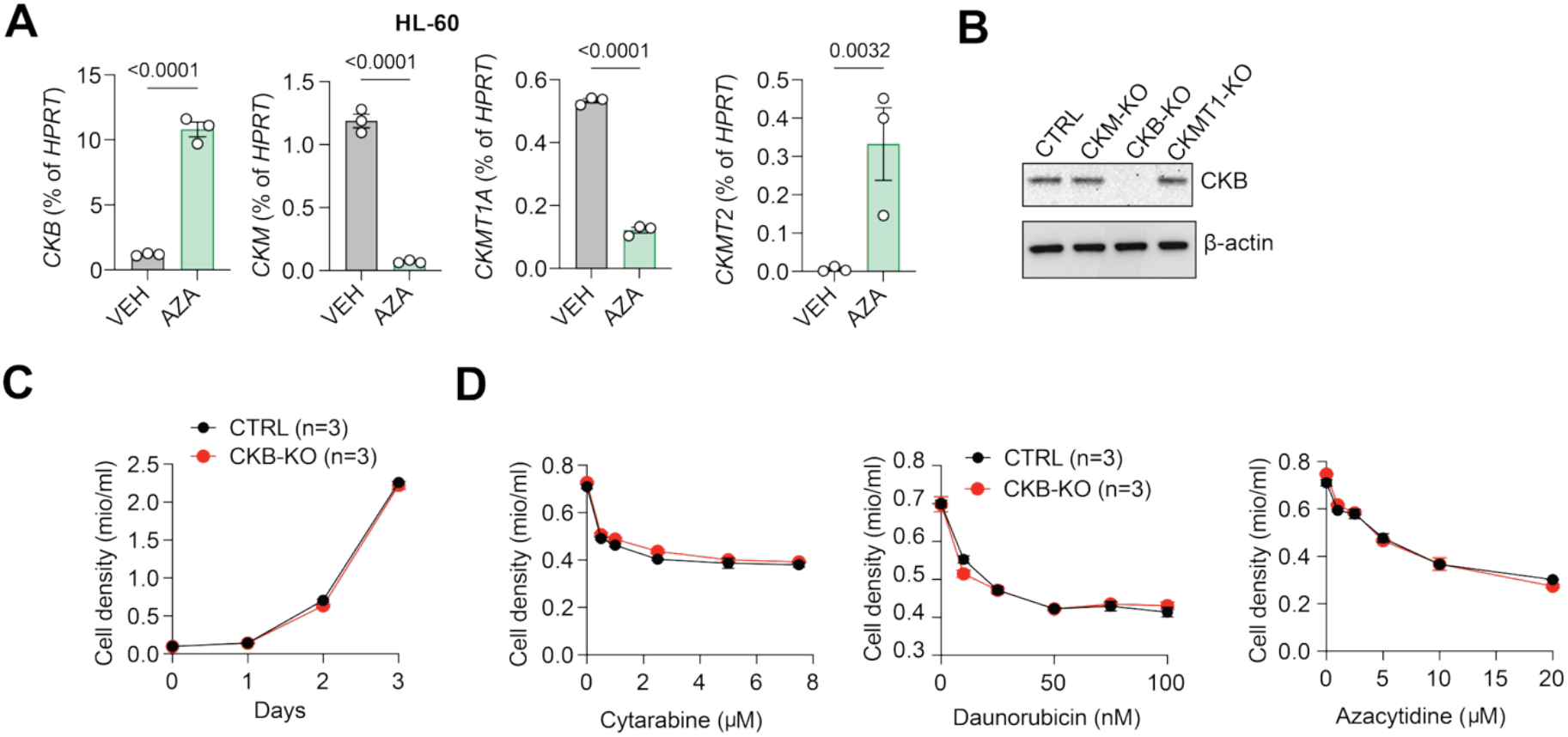
Creatine kinase expression is induced by chemotherapy and is dispensable for leukemia growth and chemotherapy sensitivity. **(A)** *CKM, CKB, CKMT1A, and CKMT2* mRNA by qPCR in HL-60 cells treated with vehicle (VEH) or 5 µM azacytidine (AZA) for 24 h. *HPRT* was used as housekeeping gene. Data are mean±SEM, n=3/group. **(B)** CKB protein expression by western blot in CTRL and genetically modified RMB-1 cells (CKM-KO, CKB-KO, and CKMT1-KO). **(C)** Cell density of CTRL and CKB-KO RMB-1 cells over a time course of 3 days. Data are mean±SEM, n=3/group. **(D)** Cell density of CTRL and CKB-KO RMB-1 cells treated with the indicated doses of cytarabine, daunorubicin, or azacytidine for 24 h. Data are mean±SEM, n=3/group. **Statistical analysis and data representation: (A)** Statistical analysis with unpaired two-tailed t-test. Representative data from one of two independent experiments. **(C)-(D)** Representative data from one of two independent experiments.

**Figure S7.**
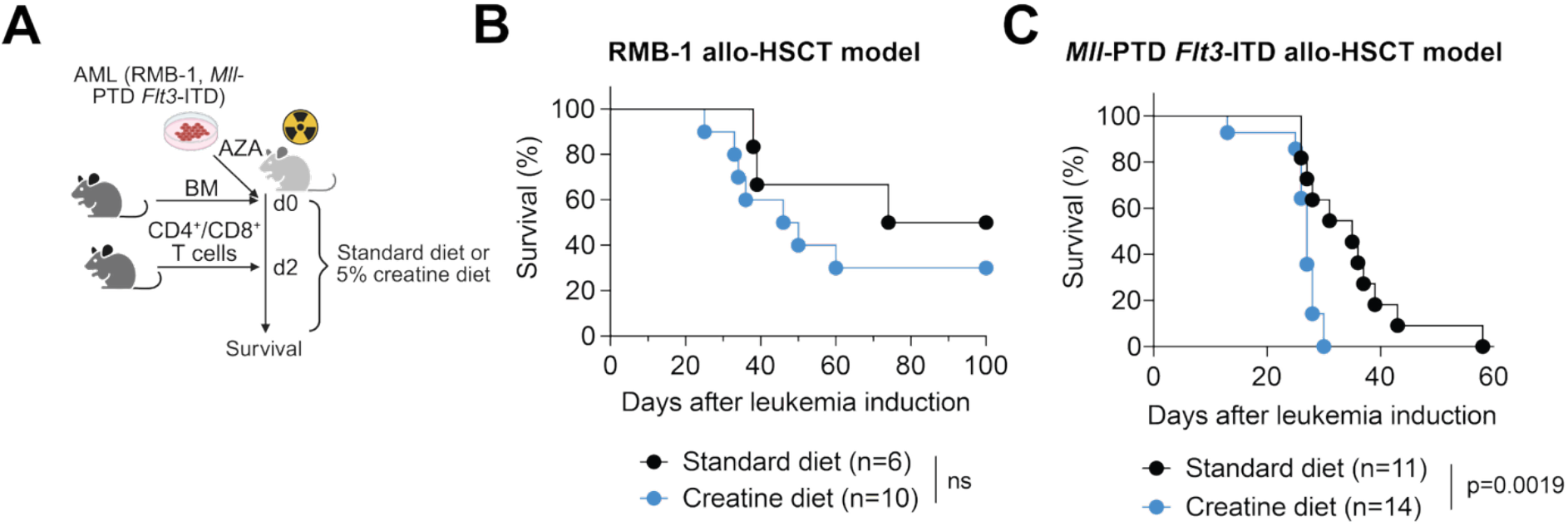
Creatine supplementation promotes leukemia progression. **(A)** Experimental model for (B) and (C), created with Biorender.com. **(B)** Survival of mice engrafted with AZA-pretreated RMB-1 AML cells and allogeneic C57BL/6 BM and T cells and kept on standard diet or 5% creatine diet, n=6-10/group as indicated. **(C)** Survival of mice engrafted with *Mll*-PTD *Flt3*-ITD AML cells and allogeneic BALB/c BM and T cells and kept on standard diet or 5% creatine diet, n=11-14/group as indicated. **Statistical analysis and data representation: (B)-(C)** Statistical analysis with Mantel-Cox test. Pooled data from three independent experiments.

**Figure S8.**
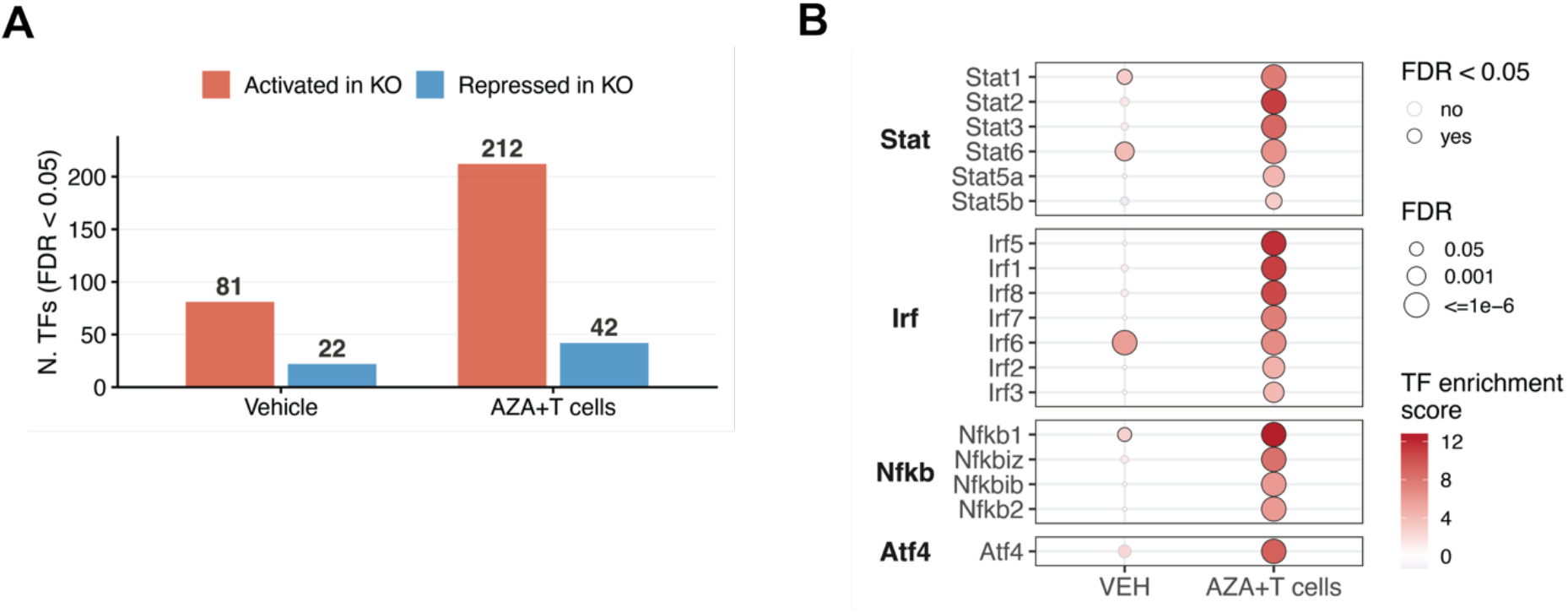
SLC6A8-deficient leukemia displays rewired inflammatory transcriptional signatures under chemo-immune pressure. **(A)** Number of transcription factors (TFs) with significantly altered inferred activity in SLC6A8-KO relative to CTRL cells, n=5/group, derived from the differential expression statistics using the CollecTRI regulon network and decoupleR TF inference. TFs with (adjusted) p < 0.05 are counted as activated (left) or repressed (right) in KO, separately for VEH and AZA+T cells conditions. **(B)** Enrichment statistics of four main TF families related to the inflammatory response. Only TFs enriched at least in the AZA+T cells condition are displayed and ranked by average TF enrichment score within each family. The dots depict the enrichment score computed by decoupleR (red colour scale) as well as their BH-corrected p-values (dot size).

**Figure S9.**
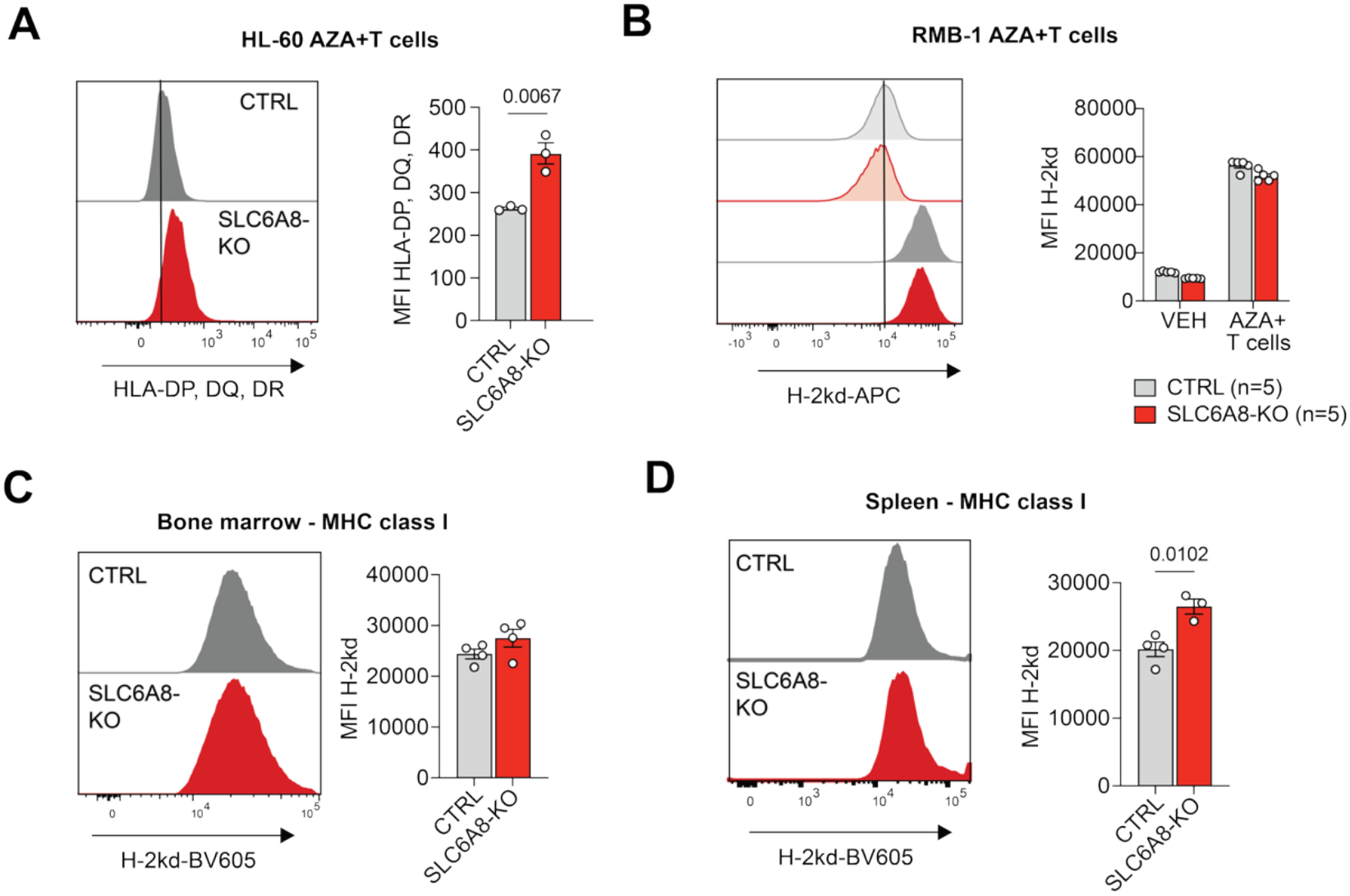
Loss of SLC6A8 enhances HLA class II expression upon exposure to allogeneic T cells. **(A)** Flow cytometry assessment of HLA class II expression on HL-60 AML cells exposed to allogeneic T cells *in vitro*. Left: representative flow cytometry histograms. Right: quantification. Data are mean±SEM, n=3/group. **(B)** Flow cytometry assessment of cell surface expression of MHC-I (H-2kd) on SLC6A8-KO and CTRL RMB-1 cells cultured *in vitro* under the indicated conditions. Upper panels: representative flow cytometry histograms. Lower panels: quantification. Data are mean±SEM, n=5/condition. **(C)-(D)** Flow cytometry assessment of MHC-I (H-2kd) expression on mCherry^+^ RMB-1 AML cells in the BM (C) or spleen (D) of mice engrafted with CTRL or SLC6A8-KO AML and allogeneic C57BL/6 BM and T cells according to the scheme in Fig. 3G. Left: representative flow cytometry histograms. Right: quantification. Data are mean±SEM, n=4/group. **Statistical analysis and data representation: (A)** Statistical analysis with unpaired two-tailed t-test. Representative data from one of three independent experiments. **(B)** Statistical analysis with Two-way ANOVA with correction for multiple comparisons. Representative data from one of two independent experiments. **(C)-(D)** Statistical analysis with unpaired two-tailed t-test. Representative data from one of three independent experiments.

**Figure S10.**
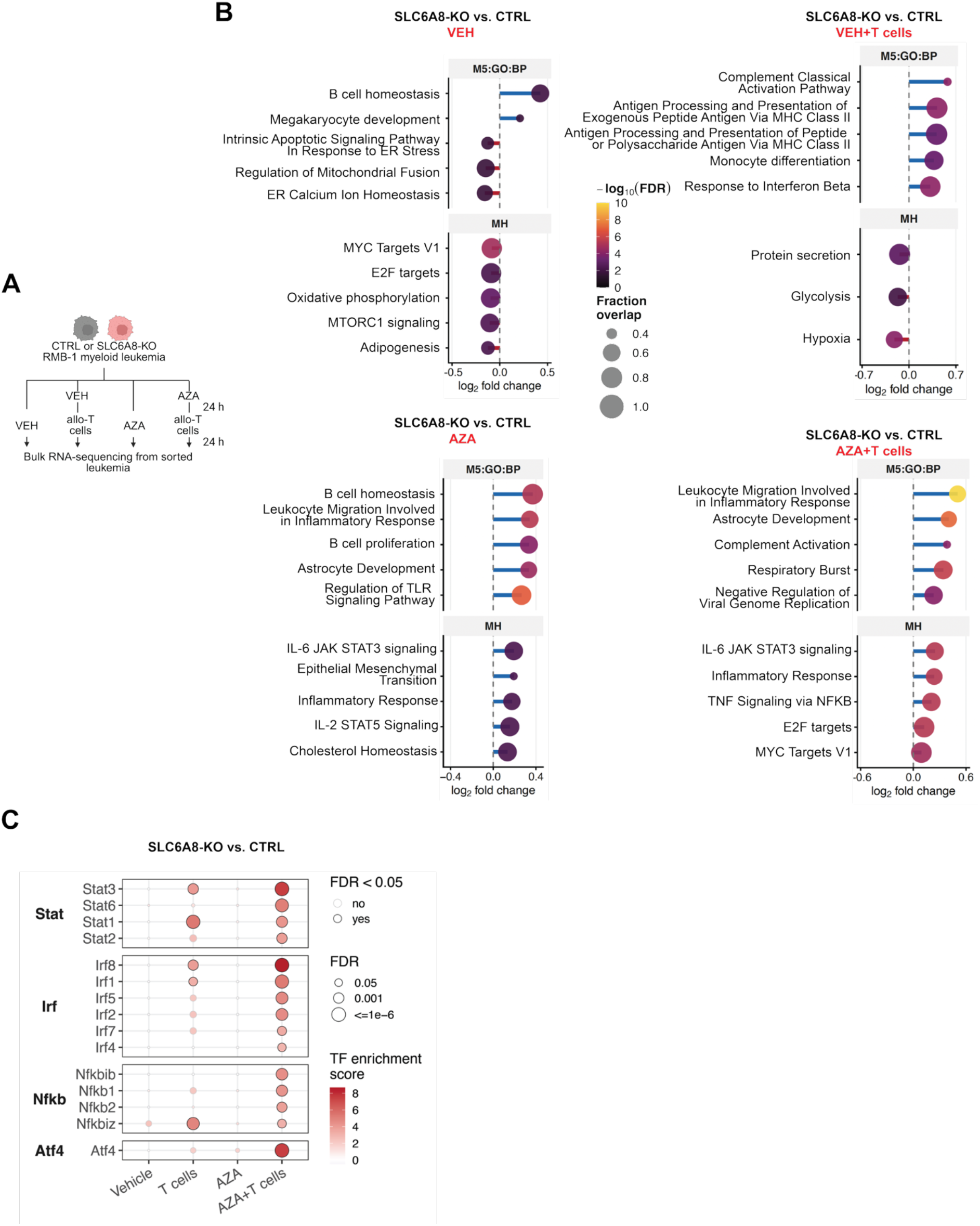
Immune pressure is the major driver of inflammatory transcriptional remodeling in SLC6A8-KO leukemia. **(A)** SLC6A8-KO and CTRL RMB-1 leukemia were treated as follows: 24 h with vehicle (VEH) or 2.5 µM azacytidine (AZA) followed by 24 h culture without or with activated allogeneic C57BL/6 CD3^+^ T cells (AML:T cell ratio 1:1). Surviving leukemia cells were sorted by flow cytometry and subjected to transcriptomics analysis, n=5/group. **(B)** Top 5 enriched and depleted Hallmark gene sets for KO vs. CTRL comparison for each treatment, ranked by log_2_FC. Dot size indicates the fraction of genes in the gene set present in the list of measured genes. **(C)** Enrichment statistics of four main TF families related to the IFN/inflammatory response in the VEH, T cells, AZA and AZA+T cells conditions. Only TFs significantly enriched in at least one of the T cell conditions (T cells or AZA+T cells) are displayed, ranked by TF enrichment score in the AZA+T cells condition within each family. Colour fill shows the enrichment score computed by decoupleR and dot size the BH-corrected p-value. **Statistical analysis and data representation:** Statistical analysis of n=5 replicates/group as described in Methods.

**Figure S11.**
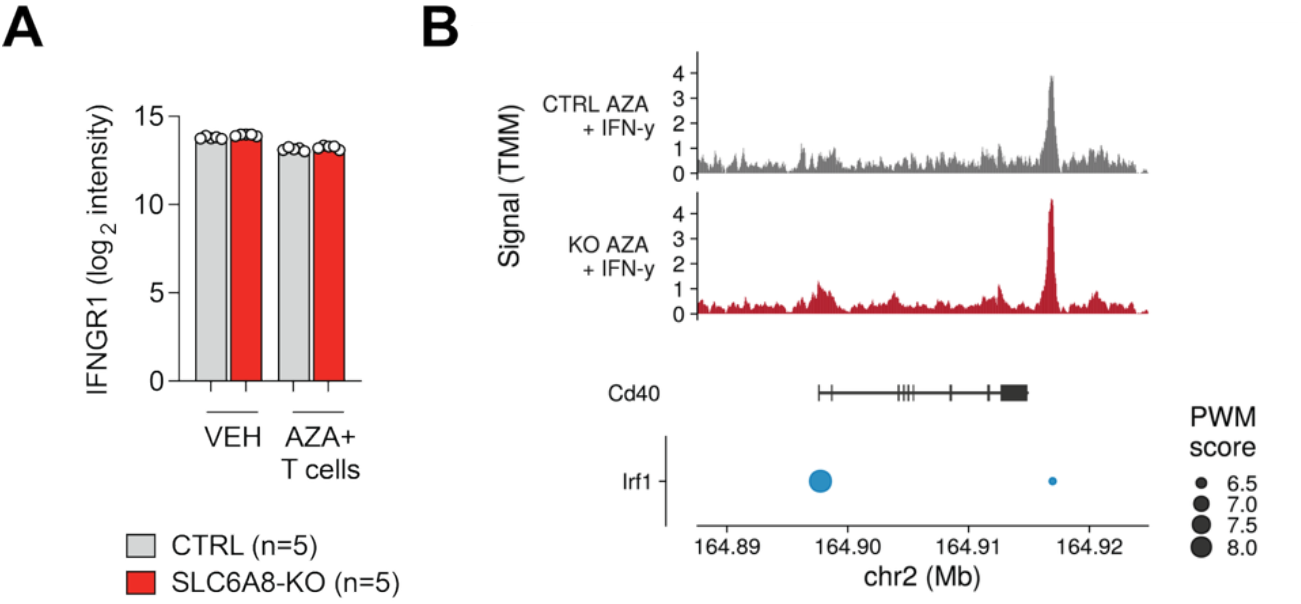
Loss of SLC6A8 alters chromatin accessibility at specific inflammatory loci. **(A)** Total protein abundance of IFNGR1 in RMB-1 CTRL and SLC6A8-KO cells exposed to the indicated conditions. Data are mean ± SEM, n=5/group. **(B)** Chromatin accessibility at the *Cd40* locus in SLC6A8-KO and CTRL RMB-1 cells under AZA+IFN-γ. Top: normalized (TMM) accessibility, shown as fragment coverage in 50-bp bins and averaged across biological replicates (CTRL, n=3; KO, n=4). Middle: genome model from GENCODE vM35, showing the gene body. Bottom: predicted transcription factor binding sites within ATAC-seq consensus peaks with scores indicated as point size.

**Figure S12.**
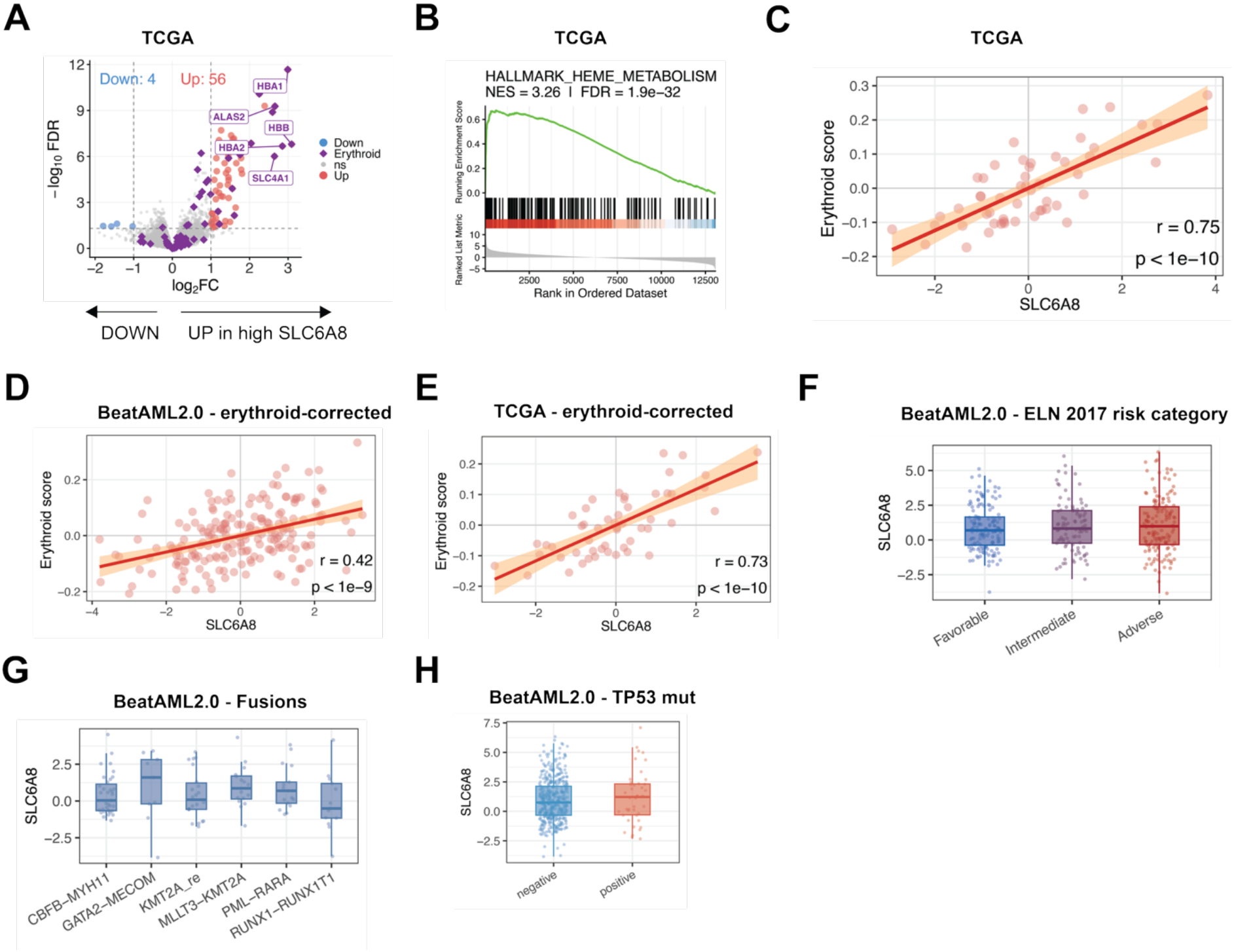
High SLC6A8 mRNA correlates with erythroid transcriptional signatures in the TCGA-LAML cohort. **(A)** Volcano plot depicting genes correlated or anti-correlated with high *SLC6A8* expression in the TCGA cohort, highlighting erythroid markers genes recovered from public databases (PanglaoDB, CellMarker 2.0). **(B)** Gene set enrichment analysis for the correlation between high *SLC6A8* expression and the Hallmark Heme metabolism gene set. **(C)** Correlation between *SLC6A8* mRNA and erythroid score in the TCGA cohort. A linear fit is depicted to represent the trend. **(D)** Correlation between *SLC6A8* mRNA and erythroid score in the BeatAML2.0 cohort, adjusted for the estimated erythroid composition in the samples. A linear fit is depicted to represent the trend. **(E)** Correlation between *SLC6A8* mRNA and erythroid score in the TCGA cohort, adjusted for the estimated erythroid composition in the samples. A linear fit is depicted to represent the trend. **(F)-(H)** *SLC6A8* expression in AML patients stratified according to (F) to the ELN 2017 risk category, (G) the presence of driver fusions and (H) the TP53 mutational status, based on the BeatAML2.0 cohort. **Statistical analysis and data representation: (A)-(C), (E)** Statistical analysis on 52 samples testing transcriptomics-wise association with SLC6A8 via linear modeling, as described in Methods. **(D)** Statistical analysis on 191 samples testing transcriptomics-wise association with SLC6A8 via linear modeling. **(F)-(H)** Statistical analysis of 483 samples using two-sample Wilcoxon test (TP53 mutation) and Kruskal-Wallis rank sum tests (ELN2017 and Fusions), none returning significant p-value.

## Materials and Methods

### Cell lines

RMB-1 mCherry^+^, HL-60, ML-2, and MOLM-13 cell lines were a gift from Robert Zeiser. C1498 cells were purchased from ATCC. Human cell lines were authenticated by short tandem repeat analysis. Cells were tested for mycoplasma on a regular basis. Cells were cultured, unless otherwise specified, in RPMI 1640 media (Invitrogen or Capricorn), supplemented with 10% fetal calf serum (Gibco or PAN), 4 mM L-glutamine, and 100 U/ml penicillin-streptomycin, in a humidified incubator at 37°C, atmospheric oxygen supplemented with 5% CO_2_. Cell counting was performed manually with a Neubauer’s chamber or automatically on a Vi-CELL BLU cell viability analyzer (Beckman Coulter).

### Chemicals and treatments

Cytarabine (CYT), daunorubicin (DAU), and azacytidine (AZA) were purchased from SelleckChemicals or Sigma. All chemicals were dissolved in DMSO for a stock solution which was further diluted in cell culture media so that the final concentration of DMSO in the culture did not exceed 0.2%. DMSO was used as vehicle. Recombinant IFN-γ was purchased from ThermoFisher Scientific. The anti-IFN-γ blocking antibody (clone XMG1.2) and rat IgG1 isotype control were purchased from Leinco Technologies. The anti-IFN-γR1 antibody (clone W20017H) was purchased from Biolegend. RGX-202 was purchased from MedChemExpress and prepared freshly in cell culture medium.

### Cell culture treatments

For discovery metabolomics and subsequent experiments, HL-60 cells were plated at a concentration of 0.25 million/ml and treated with vehicle (VEH), 0.75 µM CYT, 50 nM DAU, or 5 µM AZA for 24 h. RMB-1 cells were plated at a concentration of 0.25 million/ml and treated with VEH, 1 µM CYT, 20 nM DAU, or 2.5-5 µM AZA for 24 h.

For the recovery metabolomics experiment (Fig. 1N-O), RMB-1 cells were plated at a concentration of 0.25 million/ml and treated with VEH or 2.5 µM AZA for 24 h, followed by drug wash-out and culture in standard media for up to 48 h. Cell pellets were collected at 1 h, 6 h, 12 h, 24 h, 36 h and 48 h after the end of AZA treatment for metabolic analysis. Cell-free supernatants were collected at 24 h and 48 h after the end of AZA treatment for metabolic analysis.

For the serial treatment experiment (Fig. 1P), RMB-1 cells were plated at a concentration of 0.25 million/ml and treated with vehicle or 2.5 µM AZA for 48 h. Following a drug wash-out, the cells were rested for 6 days in standard media. The sequence of treatment and rest corresponded to one cycle of AZA treatment. This treatment was repeated for a total of 3 cycles. Cell pellets were preserved for metabolite extraction immediately after the end of treatment.

For creatine abundance measurement, MOLM-13 cells were treated with 3 µM AZA for 24 h and ML-2 cells were treated with 4 µM AZA for 24 h. C1498 cells were treated with 5 µM AZA for 24 h. All concentrations were determined to induce ∼50% growth inhibition.

For determining the effects of chemotherapy on CTRL and SLC6A8-KO AML cells, cells were plated at a concentration of 0.5 million/ml in 96-well-plates and incubated with the indicated drug concentrations for 24 h. Cell count at the end of treatment was determined on a Vi-CELL BLU cell viability analyzer (Beckman Coulter).

IFN-γ was used at 10 ng/ml unless otherwise stated. Anti-IFN-γ and isotype control antibodies were used at 10 µg/ml. The anti-IFN-γR1 antibody was used at 1 µg/ml. RGX-202 was used at 10 mM.

### Primary patient samples

Patient samples were collected and cryopreserved at the University Hospital Basel, Switzerland. All patients signed written informed consent, and the study was approved by the responsible authorities (Ethikkommission Nordwest-und Zentralschweiz, approval ID 2025-01775) and registered at clinicaltrials.gov (NCT07189117). Peripheral blood and BM mononuclear cells were separated from whole blood and BM aspirates using density gradient centrifugation and cryopreserved in 90% fetal calf serum with 10% DMSO. Samples were stored in liquid nitrogen. Information on demographics and disease characteristics are included in **Table S5**.

For RNA-sequencing and HPLC-MS/MS, samples with >80% blast percentage were selected. Samples were thawed, counted and equal cell numbers were cultured in RPMI 1640 media (Capricorn), supplemented with 20% fetal calf serum (PAN), 4 mM L-glutamine, and 100 U/ml penicillin-streptomycin, in a humidified incubator at 37°C, atmospheric oxygen supplemented with 5% CO_2_. Treatment with vehicle (DMSO) or 5 µM AZA was performed. After 24h, cells were counted and 0.1-0.4 million cells were harvested for RNA-sequencing and metabolite extraction.

### Mice

C57BL/6J (RRID: IMSR_JAX:000664), BALB/c (RRID: IMSR_JAX:000651), NOD.Cg-*Prkdc^scid^ Il2rg^tm1Wjl^*/SzJ (NSG, RRID: IMSR_JAX:005557), and C;129S4-*Rag2^tm1.1Flv^ Il2rg^tm1.1Flv^*/J (RRID: IMSR_JAX:014593) were purchased from the Jackson Laboratory or from Janvier and bred at the Johns Hopkins University and the University of Basel. Male and female mice (8-14 weeks old, age– and sex-matched between experimental conditions) were used in all experiments. All mice were maintained in the animal facilities at the Johns Hopkins University and the University of Basel under specific-pathogen-free conditions and following institutional animal use and care guidelines. Mice were exposed to a 14 h/10 h light/dark cycle and fed ad libitum. The room temperature and humidity were maintained and monitored. The animal protocols were approved by the Johns Hopkins University Animal Care and Use Committee (research protocol MO21M453) or the cantonal veterinary authorities Basel-Stadt, Switzerland (licence 36714_3233).

### Bone marrow and splenocyte harvest for analysis or *in vivo* transfer

For BM harvest, the hind limbs of donor mice were removed, the epiphyses from femur and tibia were cut and the BM was flushed using a 2 ml syringe with a 26G needle. The cells were resuspended to form a single-cell suspension. After centrifugation, the cells were resuspended in PBS, counted, and adjusted to the desired cell concentration. For *in vivo* transfer, 5 million BM cells were injected intravenously in each recipient.

Spleens from donor mice were harvested and homogenized, followed by an erythrocyte lysis step. After centrifugation, the cells were resuspended to form a single-cell suspension for terminal analysis or T cell enrichment. Where indicated, the desired T cell fraction was enriched with the MoJoSort kits (Biolegend) according to the manufacturer’s instructions. After centrifugation, the cells were resuspended in PBS, counted, and adjusted to the desired cell concentration for intravenous injection.

### AML and allo-HSCT models

For discovery metabolomics, C57BL/6 mice were subjected to total body irradiation with 5 Gy, followed by a tail vein injection of 1 million *Mll-*PTD *Flt3*-ITD AML cells^25^. Starting on day 14 after leukemia induction, mice received intraperitoneal injections of 5 mg/kg BW AZA for four consecutive days. Five days after the last azacytidine injection, the hind limbs were harvested, and CD117^+^ cells were enriched using the EasySep™ Mouse CD117 (cKIT) Positive Selection Kit (Stemcell) according to the manufacturer’s instructions.

For RMB-1 AML induction in NSG mice, recipients received a tail vein injection of 10,000 CTRL, SLC6A8-KO or CKB-KO RMB-1 mCherry^+^ cells on d0. Starting from d7, recipients received an intraperitoneal injection of 1.5 mg/kg BW AZA or an equal volume of vehicle for 5 days. Survival was monitored or organ harvest for leukemia burden assessment was performed.

For RMB-1 AML with allo-HSCT, CTRL, SLC6A8-KO, or CKB-KO RMB-1 mCherry^+^ cells were pre-treated with 2.5 µM AZA for 24h. Recipient BALB/c mice received total body irradiation with 2×4.5 Gy on d0, followed by a tail vein injection of 5 million C57BL/6 BM cells and 25,000 AZA-pretreated RMB-1 mCherry^+^ cells on the same day. Two days later, recipient mice received a tail vein injection of 500,000 CD4^+^/CD8^+^ T cells enriched from the spleens and lymph nodes of C57BL/6 donors. The enrichment of CD4^+^ and CD8^+^ T cells was performed separately using the corresponding MoJoSort kit (Biolegend) and cells were mixed in a 1:1 ratio before injection. Survival was monitored or organ harvest for leukemia burden assessment and immune phenotyping was performed.

For RMB-1 AML with allo-HSCT and T cell depletion, the same setup was used. Mice were divided into four groups: group 1 – CD4^+^ and CD8^+^ T cell transfer + IgG2a and IgG2b isotype control; group 2 – no T cell transfer + anti-CD4 + anti-CD8 depleting antibodies; group 3 – CD4^+^ T cell transfer + IgG2b isotype control antibody + anti-CD8 depleting antibody; group 4 – CD8^+^ T cell transfer + IgG2a isotype control antibody + anti-CD4 depleting antibody. The following antibodies were used (all from BioXCell): *InVivo*MAb anti-mouse CD4 (clone GK1.5, #BE0003-1), *InVivo*MAb anti-mouse CD8a (clone 53-6.7, #BE0004-1), *InVivoMAb* rat IgG2b isotype control, anti-keyhole limpet hemocyanin (clone LTF-2, #BE0090), *InVivoMAb* rat IgG2a isotype control, anti-trinitrophenol (clone 2A3, #BE0089). 0.2 mg of each were applied via intraperitoneal injection on d-1, d1, d8 and after that weekly until the end of the experiment, and survival was monitored.

For C1498 AML induction in *Rag2^-/-^ Il2rg^-/-^* mice, recipients received a tail vein injection of 250,000 C1498 cells on d0. Starting from d7, recipients received an intraperitoneal injection of 1.5 mg/kg BW AZA or an equal volume of vehicle for 5 days, and survival was monitored.

For C1498 AML with allo-HSCT, CTRL or SLC6A8-KO C1498 cells were pre-treated with 5 µM AZA for 24 h. Recipient C57BL/6 mice received total body irradiation with 2×5 Gy on d0, followed by a tail vein injection of 5 million BALB/c BM cells and 250,000 AZA-pretreated C1498 cells on the same day. Two days later, recipient mice received a tail vein injection of 500,000 CD4^+^/CD8^+^ T cells enriched from the spleens and lymph nodes of BALB/c donors. The enrichment of CD4^+^ and CD8^+^ T cells was performed separately using the corresponding MoJoSort kit (Biolegend) and cells were mixed in a 1:1 ratio before injection. Survival was monitored.

For HL-60 AML with T cell transfer, CTRL or SLC6A8-KO HL-60 cells were pre-treated with 5 µM AZA for 24 h. NSG mice received a subcutaneous injection in the right flank with 1 million cells. Four days later, mice received a tail vein injection of 1.5 million C57BL/6 CD4^+^ and CD8^+^ T cells. The enrichment of CD4^+^ and CD8^+^ T cells was performed separately using the corresponding MoJoSort kit (Biolegend) and cells were mixed in a 1:1 ratio before injection. Tumor sizes were measured with callipers at the indicated time points. The maximal allowed tumor size was 20 mm in diameter and animals were humanely euthanized if this size was reached, or if there was tumor ulceration or bleeding. On day 22, mice were euthanized and tumors were excised and weighed.

For RMB-1 AML with allo-HSCT and creatine-supplemented diet, RMB-1 mCherry^+^ cells were pre-treated with 2.5 µM AZA for 24h. Recipient BALB/c mice received total body irradiation with 2×4.5 Gy on d0, followed by a tail vein injection of 5 million C57BL/6 BM cells and 25,000 AZA-pretreated RMB-1 mCherry^+^ cells on the same day. Two days later, recipient mice received a tail vein injection of 500,000 CD4^+^/CD8^+^ T cells enriched from the spleens and lymph nodes of C57BL/6 donors. The enrichment of CD4^+^ and CD8^+^ T cells was performed separately using the corresponding MoJoSort kit (Biolegend) and cells were mixed in a 1:1 ratio before injection. Mice were kept on AIN-93G Growing Rodent Diet With 5% creatine monohydrate or standard AIN-93G Growing Rodent Diet (Research Diets, Inc.) *ad libitum*. Survival was monitored.

For *Mll*-PTD *Flt3*-ITD AML with allo-HSCT and creatine-supplemented diet, splenocytes harvested from leukemic mice ^25^ were treated with 2.5 µM AZA for 4h. Recipient C57BL/6 mice received total body irradiation with 2×5 Gy on d0, followed by a tail vein injection of 5 million BALB/c BM cells and 50,000 *Mll*-PTD *Flt3*-ITD splenocytes. Two days later, recipient mice received a tail vein injection of 500,000 CD4^+^/CD8^+^ T cells enriched from the spleens and lymph nodes of BALB/c donors. The enrichment of CD4^+^ and CD8^+^ T cells was performed separately using the corresponding MoJoSort kit (Biolegend) and cells were mixed in a 1:1 ratio before injection. Mice were kept on AIN-93G Growing Rodent Diet With 5% creatine monohydrate or standard AIN-93G Growing Rodent Diet (Research Diets, Inc.) *ad libitum*. Survival was monitored.

### T cell isolation and activation

Mouse T cell isolation and activation was performed as previously described^63^. In brief, spleens from healthy mice were dissected, and T cells were enriched using the MojoSort Mouse CD3 T Cell Isolation Kit (Biolegend, 480024) according to the manufacturer’s instructions. T cells were activated using 5 µg/ml plate-bound anti-CD3 (InVivoMab anti-mouse CD3, BioXCell, BE0002), 0.5 µg/ml soluble anti-CD28 (InVivoMab anti-mouse CD28, BioXcell, BE0015), and 100 U/ml human IL-2 (Peprotech) in RPMI 1640 media supplemented with 10% fetal calf serum (Gibco or PAN), 4 mM L-glutamine, 100 U/ml penicillin-streptomycin, and 55 µM β-mercaptoethanol. T cells were activated for 48 h in a humidified incubator at 37 °C, atmospheric oxygen supplemented with 5% CO_2_, and then re-plated every 24h in media supplemented with 100 U/ml human IL-2.

### Leukemia-T cell co-cultures

For leukemia-T cell co-cultures, RMB-1 mCherry^+^ leukemia cells were pretreated with 2.5-5 µM AZA as indicated or VEH for 24 h, following incubation with activated C57BL/6 T cells at a 1:1 ratio. Co-culture was performed for up to 24 h, after which leukemia and T cell phenotypes were assessed by flow cytometry. Percentage T cell specific killing was calculated as:

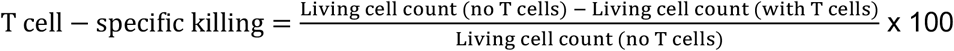

Where indicated, surviving leukemia cells were sorted by flow cytometry on a CytoFlex SRT Cell Sorter.

### Stable isotope tracing

For d_3_-creatine tracing, cells were incubated in RPMI 1640 Medium for SILAC (ThermoFisher Scientific), supplemented with dialyzed fetal calf serum (Gibco), 4 mM L-glutamine, 100 U/ml penicillin-streptomycin, 40 mg/l L-lysine hydrochloride (MedChemExpress), 100 µM L-arginine hydrochloride (MedChemExpress), and 40 µM d_3_-creatine (MedChemExpress). For U-^13^C-arginine tracing, cells were incubated in RPMI 1640 Medium for SILAC (ThermoFisher Scientific), supplemented with dialyzed fetal calf serum (Gibco), 4 mM L-glutamine, 100 U/ml penicillin-streptomycin, 40 mg/l L-lysine hydrochloride (MedChemExpress), 40 µM creatine monohydrate (Sigma), and 100 µM U-^13^C-arginine hydrochloride (MedChemExpress). After 24 h, cells were harvested and metabolites were extracted for HPLC-MS/MS.

### Metabolite extraction

For metabolite extraction from cells, 0.1-1 million cells were centrifuged for 5 min at 500 g, 4°C. The supernatant was removed and the pellet was washed with ice-cold PBS and centrifuged for a further 5 min 500 g, 4°C. Following removal of the supernatant, samples were extracted in 100-200 µl ice-cold extraction solution (methanol:acetonitrile:water 50:30:20 % v/v). Samples were immediately stored at –80°C. Before measurement, samples were centrifuged for 10 min at maximum speed, 4°C, and the clear supernatant was transferred into new 1.5 ml Eppendorf tubes. Up to 50 µl of sample was added to a plastic LC-MS vial or plate or 96-well-plate and subjected to LC-MS analysis.

For metabolite extraction from cell-free media supernatants, 500 µl supernatant was mixed with 500 µl methanol and 500 µl chloroform, vortexed for 1 min and centrifuged for 10 min at maximum speed, 4°C. The upper phase (containing polar metabolites) was transferred to a plastic LC-MS vial or 96-well-plate and subjected to LC-MS analysis.

### HPLC-MS/MS metabolomics

Chromatographic separation was performed on an Agilent 1290 Infinity II UHPLC inline using a Waters Atlantis Premier BEH Z-HILIC VanGuard FIT column (2.1 mm x 100 mm, 1.7 µm particles) or a Hilicon iHILIC(P) classic column (100 x 2.1 mm, 5 µm particles). The solvent gradient was 95% buffer B (90:10 acetonitrile:buffer A) to 20% buffer A (20 mM ammonium carbonate + 5 µM medronic acid in water). Flow rate was 150 µL/min. Column temperature was 40°C and autosampler temperature was 4°C. Injection volume was 2 µl. The LC system was coupled to a Bruker Impact II QTOF-MS or Bruker timsTOF-Pro mass spectrometer equipped with an ionBooster electrospray ionization (ESI) source. The mass spectrometer was operated in both positive and negative modes. Scan range was from 20 to 1050 Da. Mass calibration was performed at the beginning of each run.

Data were acquired with the Compass Hystar software. Data processing including detection, deconvolution, and identification of features was carried out using Metaboscape and TASQ software (versions 2022 – 2024b; Bruker). The following parameters were used for metabolite annotation: m/z matching with ±3ppm, milliSigma matching within 300, confirmation of critical metabolites (creatine, phosphocreatine, glutathione) with pure standards measured on the same instrument.

### Creatine assay

Creatine concentrations were determined where indicated using a creatine assay kit (MAK494, Sigma). 2 million cells were lysed in 100 µl Creatine Assay Buffer, centrifuged at 13,000 g for 10 min at 4°C and processed further according to the manufacturer’s instructions. Absorbance was read at 570 nm.

### Creatine kinase activity assay

Creatine kinase activity was determined using a creatine kinase activity assay kit (MAK116, Sigma). 3.5 million cells were lysed in 40 µl ice-cold potassium phosphate 50 mM buffer, pH 7.5, centrifuged at 10,000 g for 15 min at 4°C and processed further according to the manufacturer’s instructions. Absorbance was read at 340 nm after 20 min (initial) and 40 min (final) incubation at 37°C.

### RNA extraction and reverse transcription

RNA was extracted with the miRNeasy extraction kit (Qiagen) or the Quick-RNA Miniprep Kit (Zymo Research) according to the manufacturer’s instructions. RNA concentration measurements were performed with Qubit or Nanodrop. 200–2000 ng RNA underwent reverse transcription using the High-Capacity cDNA Reverse Transcription Kit (ThermoFisher Scientific) according to the manufacturer’s instructions, with RNA amounts being kept constant across samples belonging to the same experiment. RNA and cDNA were stored at –80°C.

### Bulk RNA-sequencing

250,000 – 1,000,000 cells were preserved in Zymo DNA/RNA shield and further processed by Plasmidsaurus. Following automated extraction, 500 ng RNA were used for library preparation. Bulk RNA-sequencing was performed by Plasmidsaurus using Illumina Sequencing Technology with custom analysis and annotation. Between 6-13M unique reads per sample were acquired.

### Bulk ATAC-sequencing

Chromatin accessibility profiling was performed using a modified ATAC-seq protocol^64^. For adapter annealing, 22.5 µl each of ME and R1N (or R2N) oligonucleotide (all 100 µM) were combined with 5 µl 10x annealing buffer (100 mM Tris-HCl pH 7.5, 500 mM NaCl, 10 mM EDTA), incubated in a thermocycler (95 °C for 3 min, cooled to 25 °C at 0.1 °C/s) and diluted with an equal volume of 100 % glycerol to a final adapter concentration of 22.5 µM. Tn5 assembly was performed by mixing 25 µl R1N:ME, 25 µl R2N:ME and 50 µl Tn5 (1 mg/ml in 25 mM Tris-HCl pH 7.5, 100 mM NaCl, 0.1 mM EDTA, 1 mM DTT, 0.1% IGEPAL CA-630, ∼84 % (v/v) glycerol) followed by incubation at room temperature for 45 min. Assembled Tn5 was stored at –20 °C and its activity was confirmed before use.

Cells were pelleted at 300 x g for 5 min at 4 °C and lysed for 2 min in cold resuspension buffer (RB, 10 mM Tris-HCl pH 7.5, 10 mM NaCl, 3 mM MgCl2, 0.1 % Tween-20, 1 % BSA) containing 0.025 % IGEPAL CA-630 and 0.01 % Digitonin. Lysis was stopped by addition of RB, nuclei were pelleted at 500 x g for 5 min at 4 °C, resuspended in 1% BSA in PBS and adjusted to 10,000 nuclei/µl. For each condition, 5 µl nuclei suspension (50,000 nuclei) and 5 µl assembled Tn5 were added to 40 µl transposition buffer (38.8 mM Tris-acetate, 77.6 mM potassium acetate, 11.8 mM magnesium acetate, 18.8 % dimethylformamide, 1x protease inhibitor cocktail) and incubated for 60 min at 37 °C shaking at 500 rpm. DNA was purified using the MinElute Kit and eluted in 21 µl EB. Transposed DNA was amplified in a 50 µl reaction containing NEBNext High-Fidelity 2x PCR Master Mix and 2 µM each of barcoded Nextera-compatible i5 and i7 index primers^64^. Thermocycling was 72 °C for 5 min and 98 °C for 30 s, followed by 6 cycles of 98 °C for 10 s, 65 °C for 30 s, and 72 °C for 1 min. To determine the number of additional cycles, a 2.5 µl aliquot of the pre-amplified reaction was analyzed by qPCR with SYBR Gold, and the number of cycles needed to reach a quarter of maximum fluorescence was added to the remaining reaction. Accordingly, libraries received 0 or 2 additional cycles (6 or 8 cycles in total). Libraries were purified with 1.2x SPRIselect beads and eluted in 16 µl EB.

ATAC-seq libraries were quality-controlled using the Fragment Analyzer High Sensitivity NGS Fragment Analysis Kit (Agilent). Libraries were pooled and subjected to a double-sided size-selection cleanup (1.2×-0.5×) using MAGFLO NGS magnetic beads (INTEGRA Biosciences). The final library pool was sequenced on an AVITI24 instrument (Element Biosciences) using an AVITI 2×75 Sequencing Kit with a Cloudbreak FS High Output flow cell in paired-end 2 × 51 bp mode.

### Proteomics

Cells were lysed with buffer containing 100 mM TEAB pH=8.5 / 5% SDS / 10 mM TCEP by 10 min heating at 95°C followed by 10 min sonication on a Pixul® system (Active Motif) using following settings: Pulse[N]: 50; PRF[kHz]: 1; burst rate[Hz]:20. Protein extracts were alkylated using 20 mM iodoacetamide at 25° C in the dark for 30 min. Lysates were digested, and desalted using STRAP cartridges (Protifi). In brief, samples were acidified by addition of 10% TFA (1:10) and then S-trap buffer (90% methanol, 100 mM TEAB pH 7.1) was added to the samples (6:1). Samples were briefly vortexed and loaded onto S-trapTM micro spin-columns (Protifi) and centrifuged for 1 min at 4000 g. Flow-through was discarded and spin-columns were then washed 3 times with 150 µL of S-trap buffer (each time samples were centrifuged for 1 min at 4000 g and flow-through was removed). S-trap columns were then moved to the clean tubes and 20 µL of digestion buffer (50 mM TEAB pH 8.0) and trypsin (at 1:25 enzyme to protein ratio) were added to the samples. Digestion was allowed to proceed for 1h at 47 °C. After, 40 µL of digestion buffer was added to the samples and the peptides were collected by centrifugation at 4000 g for 1 minute. To increase the recovery, S-trap columns were washed with 40 µL of 0.2% formic acid in water (400g, 1 min) and 35 µL of 0.2% formic acid in 50% acetonitrile. Eluted peptides were dried under vacuum and stored at –20 °C until further processing.

Dried peptides were resuspended in 0.1% aqueous formic acid and subjected to LC–MS/MS analysis using an Astral Zoom Mass Spectrometer fitted with a Vanquish Neo (both Thermo Fisher Scientific) and a custom-made column heater set to 60°C. Peptides were resolved using a RP-HPLC column (75 um × 30 cm) packed in-house with C18 resin (ReproSil Saphir 100 C18, 1.5 um resin; Dr. Maisch GmbH) at a flow rate of 0.2 ul/min. The following gradient was used for peptide separation: from 2 % B to 25 % B over 17 min to 35 % B over 3 min to 95 % B over 0.5 min followed by 5 min at 95 % B to 2 % B over 0.5 min. Buffer A was 0.1 % aqueous formic acid and buffer B was 80 % acetonitrile, 0.1 % formic acid.

The mass spectrometer was operated in DIA mode. MS1 scans were acquired in the Orbitrap in profile mode at a resolution of 240,000 FWHM, a scan range from 380 to 980 m/z, normalized AGC target set to 500 % and a maximum ion injection time of 30 ms. MS2 scans were acquired in the Astral analyzer over a precursor mass range of 380 to 980, an isolation window of 6 m/z without window overlap, maximum injection time of 7 ms, a scan range from 150 to 2000 m/z, normalized AGC target of 800 % with window placement optimization enabled. Peptides were fragmented by HCD (Higher-energy collisional dissociation) with collision energy set to 25 %, Pre-Accumulation was enabled and one microscan was acquired for each spectrum.

The acquired raw files were searched using SpectroNaut (v21.0, directDIA workflow, default settings) against a murine database (downloaded from Uniprot on 20250926) using the following search criteria: full tryptic specificity was required (cleavage after lysine or arginine residues, unless followed by proline); 3 missed cleavages were allowed; carbamidomethylation (C) was set as fixed modification; oxidation (M), N-acetylation (N-term). Quantitative fragment group data (FG.MS2RawQuantity) was exported from Spectronaut and analyzed using ProteoFlux v1.10.4. (https://zenodo.org/records/21267281). To meet additional assumptions (normality and homoscedasticity) underlying the use of linear regression models and t-Tests, intensity signals were transformed from the linear to the log-scale. Data was normalised by median_equalization, imputed using the LC-ConMed method and p-values for differential expression analysis were calculated using a limma-based empirical Bayes framework. The resulting per protein and condition comparison p-values were adjusted for multiple testing using the Benjamini-Hochberg method.

### Quantitative real-time PCR

Quantitative real-time PCR was performed in 96-well or 384-well plates using the iTaq Universal Probes kit (Biorad) or the TaqMan Fast Advanced Master Mix (Applied Biosystems) and commercially validated TaqMan Real-Time PCR Assays (Applied Biosystems). The following assays were used: mouse *Gatm* (Mm00491879_m1), mouse *Slc6a8* (Mm00506023_m1), mouse *Hprt* (Mm03024075_m1), mouse *Ckb* (Mm04213225_s1), mouse *Ckm* (Mm01321487_m1), mouse *Ckmt1* (Mm00438221_m1), human *GATM* (Hs00155208_m1), human *SLC6A8* (Hs00940515_m1), human *GAPDH* (Hs02786624_g1), human *HPRT* (Hs02800695_m1), human *CKM* (Hs00176490_m1), human *CKB* (Hs00176484_m1), human *CKMT1B* (Hs00179727_m1), human *CKMT2* (Hs00176502_m1). Analysis was performed on a QuantStudio 5 instrument (Applied Biosystems) or the ViiA 7 Real-Time PCR System. Quantification was performed using the ΔCt method expressing the expression of the gene of interest as percentage of the expression of the housekeeping gene (*Hprt* for mouse, *GAPDH* or *HPRT* for human).

### Western blot

For western blot analysis, cells were lysed in 1X cell lysis buffer (Cell Signaling Technology), containing 1X Protease/ Phosphatase Inhibitor Cocktail (Cell Signaling). Samples were frozen and thawed, then centrifuged at 13,000 g for 10 min at 4°C. Cleared protein lysate was denatured with LDS loading buffer (1X loading dye, 1 mM DTT) for 5 min at 95°C.

For phospho-STAT1 (pSTAT1) and total STAT1 immunoblotting, cells were lysed directly in 2× sample buffer containing Bolt LDS Sample Buffer (Thermo Fisher Scientific, cat. no. B0007) and 5% β-mercaptoethanol. Samples were immediately heated at 95°C for 8 min and centrifuged twice for 20 min at 4°C. The clarified supernatants were collected for analysis.

Protein samples were separated on precast 4-12% Bis-Tris protein gels (Merck). Proteins were transferred onto nitrocellulose membranes using a Trans-Blot Turbo Transfer System (Biorad) according to the manufacturer’s protocols. Membranes were blocked for 1 h at room temperature with 5% w/v milk powder or bovine serum albumin (BSA) in Tris-buffered saline with 0.1% v/v Tween-20 (TBS-T), then incubated with primary antibodies in 5% w/v blotting buffer in TBS-T overnight at 4°C, with shaking. All primary antibody incubations were followed by at least three washes with TBS-T, then incubation with secondary horseradish peroxidase-conjugated antibody (goat anti-rabbit IgG 1:10,000 (BioConcept, 7074) in 5% w/v blotting buffer, for 1 h at room temperature. After a further three washes with TBS-T, membranes were incubated for 10 min with Clarity Western ECL Substrate (BioRad) or SuperSignal™ West Femto Maximum Sensitivity Substrate (ThermoFisher), and visualized using a Vilber Fusion FX Western Blot Imager.

Antibodies used were: GATM (abcam, #ab228937), CKB (abcam, #ab92452), Phospho-STAT1 (Tyr701) (Cell Signaling Technology, clone D4A7, #7649), STAT1 (Cell Signaling Technology, #9172), β-actin (Cell Signaling Technology, clone 13E5, #4970L). All primary antibodies were used at a dilution of 1:1000. Densitometry was performed using ImageJ.

### CRISPR/Cas9 gene editing

CRISPR/Cas9 gene editing was performed by nucleofection or lentiviral transduction. For nucleofection, 2 million cells were electroporated with gRNA-Cas9 ribonucleoproteins (RNPs) using the 4D-Nucleofector system (Lonza). RNPs were prepared by complexing 180 pmol of gRNAs (IDT) with 60 pmol of Cas9 nuclease (IDT). Electroporation was performed in P4 Primary Cell solution using the program CM137. The nucleotide sequences for gRNA are listed in **Table S6**. A mix of three different gRNA was used.

To generate stable edited cell lines, gRNA were cloned into the lentiCRISPR v2 with puromycin resistance plasmid backbone. In brief, gRNA oligonucleotides were ordered from IDT according to the following format: GGAAAGGACGAAACACC*(20-21-bp protospacer)*GTTTTAGAGCTAGAAATAGCAA. The protospacer sequences are listed in **Table S7**. LentiCRISPR v2 was cut with BsmBI overnight at 37°C, ran on an agarose gel, and the 12,988 bp band was extracted. 0.04 pmol linearized lentiCRISPR v2 and 0.2 µmol gRNA oligonucleotide were assembled using the NEBuilder HiFi DNA Assembly kit at 50°C for 60 minutes. Plasmids were transformed into Stable Competent *E. coli* cells (New England Biolabs) via heat shock using 2 µl of the assembled product. Successfully transformed bacteria were selected by growth in LB broth with ampicillin, and plasmid DNA was extracted using a Qiagen MiniPrep kit. Lentiviral particles were produced using HEK293T cells. HEK293T cells were cultured at a density of 0.8×10^6^ cells/ml in 1.92 ml of Reduced Serum Medium (OPTI-MEM) with GlutaMAX Supplement (Invitrogen), supplemented with 5% FCS, 1 mM sodium pyruvate, and 1× MEM nonessential amino acid in a 6-well plate for one night before transfection with the plasmid of interest. On the day of transfection, the transfection mixture was prepared by making tubes A and B as follows: tube A was prepared with 210 µl OPTI-MEM (no serum), 1.79 µg transfer plasmid, 1.28 µg psPAX2 packaging plasmid, 0.55 µg pMD2.G VSV-G envelope expressing plasmid and 6.19 µl p3000 reagent (Thermo Fisher Scientific), and tube B with 210 µl Opti-MEM (no serum) and 7.04 µl Lipofectamine 3000 (Thermo Fisher Scientific). Tubes A and B were vortexed, combined, and incubated at room temperature for 15 min. 850 µl of media was removed from each well of HEK293T cells, and after 15 min, the transfection mixture was added to the appropriate wells. HEK293T cells were incubated at 37°C for 6 hours, at which point the media was removed and replaced with 2 ml OPTI-MEM (5% FCS, 1 mM sodium pyruvate, 1× MEM nonessential amino acid) supplemented with 1:500 ViralBoost VB100 from Alstem Bio. The culture media was collected 24 h later, fresh media was added to the cells, and this media was collected after a further 24 h and combined with the previously collected media. This was filtered using a 0.45 µm filter and 1/3 volume Lenti-X concentrator (Takara Bio) was added. This was mixed by inversion, incubated at 4°C overnight, then centrifuged at 1500 g, 4°C, for 45 min. The supernatant was removed and the pellet of virus was resuspended in OPTI-MEM (without supplements) at 1/10 – 1/100 of the original volume, before freezing at –80°C (for storage) or addition to cells. For lentiviral transduction, 2×10^6^ AML cells were plated in 2 ml media (RPMI 1640 with supplements) in a 6-well-plate with the addition of 8 µg/ml polybrene and the virus. Spinoculation was performed at 2000 g and 30°C for 2 h. Puromycin selection (2 µg/ml) was started 48 h after transduction. Where indicated, single cells were sorted by flow cytometry.

### Inference of CRISPR edits

For determination of CRISPR/Cas9 gene editing efficiency with Inference of CRISPR edits (ICE), DNA from CTRL and SLC6A8-KO cells was extracted using the Quick-DNA MiniPrep (Zymo Research). The gene region of interest was amplified by PCR using the NEBNext Ultra II Q5 Master mix (New England Biolabs) according to the manufacturer’s instructions. Forward and reverse primers specific for each gRNA were designed and are listed in **Table S7**. The PCR product was run on a 1% agarose gel and the band corresponding to the expected size of PCR product was extracted with MinElute Gel Extraction Kit (Qiagen). Sanger sequencing was performed by MicroSynth using gRNA-specific primers (listed in **Table S7**). Analysis and calculation of the knockout score was done at https://ice.editco.bio/#/ by comparing the sequences of CTRL and SLC6A8-KO cells.

### Flow cytometry

Single cell suspensions were plated in 96-well plates (0.1-0.5 million cells/well) and centrifuged at 1300 rpm for 5 min at 4°C. Fc block (1:100) was performed in PBS with 2% FCS and 1 mM EDTA for 10 min at 4°C. Cell surface staining was performed in PBS with 2% FCS and 1 mM EDTA for 20 min at 4°C and dead cells were excluded using the LIVE/DEAD Fixable Dead Cell Stain kits (ThermoFisher Scientific). Antibodies were diluted 1:200 unless otherwise stated. If acquisition was not performed immediately, samples were fixed with BD Cytofix/Cytoperm Fixation and Permeabilisation solution or Fixation Buffer (Biolegend) for 20 min at RT and stored at 4°C. Flow cytometry analyses were performed on the FACSymphony, LSR Fortessa (BD Biosciences) or Cytek Northern Lights, and then analyzed with FlowJo 10 Software (TreeStar). The following antibodies were used for cell surface staining: anti-mouse CD45-Brilliant Ultra Violet 395 (BD, clone 30-F11, cat #564279), anti-mouse H-2kd-Brilliant Violet 605 (Biolegend, clone SF1-1.1, cat #116631), anti-mouse I-A/I-E-Brilliant Violet 785 (Biolegend, clone M5/114.15.2, cat #107645), anti-mouse-CD40-Brilliant Violet 785 (Biolegend, clone 3/23, cat #124645), anti-mouse CD80-FITC (Biolegend, clone 16-10A1, cat #104706), anti-mouse-CD3-PE-Cy7 (Biolegend, clone 17A2, cat #100220), anti-mouse CD3-APC/Fire810 (Biolegend, clone 17A2, cat #100268), anti-mouse CD4-APC-Cy7 (Biolegend, clone GK1.5, cat #100414), anti-mouse CD4-Brilliant Violet 650 (Biolegend, clone RM4-5, cat #100545), anti-mouse CD8-PE-Cy7 (Biolegend, clone 53-6.7, cat #100722), anti-mouse CD8-FITC (Biolegend, clone 53-6.7, cat #100706), anti-mouse/human CD44-PE-Cy7 (Biolegend, clone IM7, cat # 103020), anti-human HLA-DP,DQ, DR-PE (Biolegend, clone Tü39, cat #361715).

### Migration assay

To study migration along a CXCL12 gradient, 600 µl RPMI 1640 media ± 50 ng/ml CXCL12 were plated into the lower chamber of a 24-well transwell plate (Corning, pore size 5 µm). Two-hundred µl RMB-1 cell suspension (1×10^6^ cells/ml) in RPMI 1640 media without chemokine addition were plated into the insert. The plate was incubated in a humidified incubator at 37°C, atmospheric oxygen supplemented with 5% CO_2_ for 6 h. After that, the number of migrated and non-migrated cells was determined on a Cytek Aurora flow cytometer acquiring a pre-determined equal volume from each sample. The percentage migrated cells was calculated as % = (migrated cells) / (migrated+non-migrated cells).

## Data analysis

Statistical analysis was performed using Excel (Microsoft) or Prism 10 software (GraphPad) unless otherwise indicated. Data distribution was assumed to be normal, but this was not formally tested. Results are represented as the mean ± s.e.m. Comparisons of two groups were calculated using an unpaired two-tailed t-test; when one group was used for normalization and thus all values in this group were equal (for example, 0 or 1), a one-sample t-test was applied. Comparisons of more than two groups were calculated using ordinary one-way ANOVA or two-way ANOVA with the indicated appropriate test for multiple comparisons.

## Metabolomics analysis

The untargeted and targeted LC-MS metabolomics data were loaded and analyzed using R version 4.5.2^65^. For untargeted data, raw metabolite peak intensities were organized and analyzed per model, separately. The features with no measured values were removed and the missing intensity values were imputed. The type of imputation was chosen based on the type of missingness vs. mean feature intensity observed for each model. For RMB-1 and HL-60, the missingness was independent of the mean feature intensity, which indicates that the missingness was likely not related to the limit of detection (LOD). KNN imputation was performed on both models, since it proved efficient in this case scenario and on metabolomics data in general^66,67^. In contrast, the *Mll-*PTD *Flt3*-ITD AML model showed an inflation of missing values at low intensity, indicating missing non at random (MNAR) values. The values of this model were imputed using the half-minimum imputation method, as recommended by a previous benchmark^68^. The imputed data were then log transformed and scaled using the Pareto scaling^69^. PCA was applied on the transposed matrices of the untargeted metabolomics data.

For targeted data, for the RMB-1 and HL-60 models, which comprised four treatments group (VEH, CYT, DAU and AZA), each with 3 biological replicates, differential abundance was tested using a one-way ANOVA applied independently to each metabolite. Specific differences in comparison to the vehicle group were assessed using Tukey HSD post-hoc tests. For the *Mll-*PTD *Flt3*-ITD AML model, which involved two groups, a two-sample t-test was used on each metabolite. In all models, p-values were adjusted for multiple testing using the Benjamini-Hochberg false discovery rate (FDR) procedure. Adjusted p-values were interpreted using a 0.05 significance threshold. The metabolite abundances were visualized using the *ComplexHeatmap* R package^70^, displaying the row-wise Z-score. Hits overlap was displayed using the *UpSetR* R package^71^.

### Bulk RNA-seq analysis

We ran three independent bulk RNA-seq experiments on murine RMB-1 cells carrying an SLC6A8 knockout (KO) or control (CTRL) assigned to different treatment groups:

– Experiment 1: CTRL and KO cells treated with vehicle, T cells, AZA or AZA+T cells (8 groups, n=5/group)
– Experiment 2: CTRL and KO cells treated with vehicle or AZA+T cells (4 groups, n=5/group)
– Experiment 3: CTRL and KO cells treated with AZA+IFN-γ (2 groups, n=5/group) Plasmidsaurus service performed library preparation, sequencing and primary data processing; reads were trimmed for poly(X) tails and low-quality 3’ bases with fastp^72^ (minimum Phred 15, minimum length 50 bp) and aligned to the mouse reference genome (GRCm39) with STAR^73^. PCR duplicates were collapsed by UMI and alignment position with UMICollapse^74^. Strand-specific gene-level counts were generated with featureCounts^75^, assigning reads over exons and 3’ UTRs to Ensembl gene IDs.

We performed downstream analyses on the gene-level count matrices provided by Plasmidsaurus in R (v4.5.2) with edgeR^76^ (v4.8.2) and limma-voom^77^ (v3.66.0). Genes with zero counts were filtered out, followed by lowly expressed gene filtering using edgeR’s filterByExpr method and using the experiment’s design matrix and default settings. For each experiment, we fitted a negative binomial generalized linear model with one coefficient per genotype – treatment group. Within each treatment condition, we compared KO against CTRL with quasi-likelihood F-tests. Multiple testing was adjusted using the Benjamini-Hochberg procedure.

We performed competitive gene set testing using the CAMERA^78^ method on all genes retained after filtering, using the gene-wise statistics from each of the KO vs CTRL contrast. Gene sets came from the mouse collections of MSigDB v2025.1 and were tested using a 15-200 genes limit and corrected p-values with the Benjamini-Hochberg procedure within each collection and contrast. For each gene set effect size, we report the average log2fold change of member genes, signed by the effect direction returned by CAMERA.

To identify transcriptional regulators associated with the altered transcriptomics changes observed in all three experiments of the RMB-1 model, we inferred TF activity from known TF-target gene relationships. TF activity was inferred based on the input of the differential expression results of each contrast using the decoupleR R package (v2.16)^79^ and the univariate linear model (ULM) method. As recommended by the authors, the input of decoupleR was the gene-wise log2FC multiplied by the associated –log(10) p-value of the differential gene expression. We used the mouse CollecTRI regulon network^80^ as prior knowledge of TF-regulon associations, covering 1066 TFs and 6035 target genes and containing modes of regulation (activation, inhibition) based on known relationships of TFs with activated/ suppressed genes. Since STAT1 is not present in CollecTRI and since it plays a major role in tumor development and immune evasion, we complemented the prior knowledge network using ChIP-Atlas (v3.0)^81^ and SIGNOR (v4.0)^82^. For ChIP-Atlas, we first retrieved genes with STAT1 ChIP-seq peak within +/− 1 kb of the TSS and scored them by their average MACS2 –log10 q-value across all experiments involving STAT1 in the database. From the ranked score distribution, we inferred the (knee) inflection point and kept genes scoring above this point, yielding 663 targets. Importantly, ChIP-seq experiments do not tell whether a TF activates or represses a gene. For this reason, we assigned a default +1 (activation) mode of regulation to these genes. For SIGNOR, we retrieved signed STAT1 interactions for mouse STAT1 (Uniprot P42225) and kept only interactions where STAT1 regulates a target’s transcription, yielding 13 targets. Based on this extended TF-regulon network, we estimated TF activity using decoupleR’s method. We tested only TFs with at least five targets among the genes in the data. The t-value of the slope was used as the activity score and p-values were corrected using the Benjamini-Hochberg procedure.

### ATAC-seq analysis

Raw reads were processed using the nf-core/atacseq pipeline v2.1.2^83^ run with Nextflow v24.10.4^84^ and Singularity containers on the HPC of the University of Basel. Reads were aligned to the mouse GRCm39 primary assembly with GENCODE vM35 annotation^85^. The ENCODE mm10 blacklist v2^83^ was lifted over to GRCm39 and supplied to the pipeline. A consensus peak set was built with the DiffBind R package^86^ from the per-replicate MACS2 narrow peaks. A peak was retained if it was called in at least two replicates of at least one experimental group (condition x treatment). Consensus peaks were re-centered on their summits and resized to 401 bp windows (summit +/− 200 bp). Fragments overlapping each window were counted from the filtered, blacklist-removed, cleaned BAM files, yielding ∼106,000 regions. One sample (CTRL condition, AZA+IFN-γ treatment, replicate 2) was excluded from all downstream analysis since it presented major QC deviations from all other samples and from its own experimental group. The final dataset comprised 23 samples (three replicates for CTRL AZA+IFN-γ, four for all other groups).

Differential accessibility was tested using the edgeR R package. Low-count regions were removed and normalization factors were computed with the trimmed mean of M-values method (TMM)^87^ on the reads in the consensus peaks. A quasi-likelihood negative binomial generalized linear model was fitted with a group-means design strategy and using the robust estimation of the quasi-likelihood dispersions from edgeR. KO versus CTRL was tested within each treatment (VEH, AZA and AZA+IFN-γ) using quasi-likelihood F-tests, and p-values were adjusted with the Benjamini-Hochberg method. Consensus regions were annotated with the ChIPseeker R package^88^ using a transcript database built from the same GENCODE vM35 GTF used for alignment. Each region was assigned to its nearest gene, genomic feature class and distance to a transcription start site (TSS). The logFC of regions related to candidate genes (co-stimulatory genes and MHC class II genes) were reported and annotated as either promoter or distal regions.

Motif enrichment analysis was performed using the monaLisa R package^89^ and using position weight matrices (PWMs) from the JASPAR 2024 CORE vertebrate collection^90^. A curated set of transcription factor families was used for the rest of the motif analysis; STAT, IRF, NF-κB/REL, RFX, CREB1/ATF1, NF-Y and USF. To compare motif enrichment between treatments, a class-based analysis was run separately for each of the three KO versus CTRL contrasts (VEH, AZA and AZA+IFN-γ). Using the differential abundance results, regions were labelled as gained in KO (FDR < 0.05, log2FC > 0), lost in KO (FDR < 0.05, log2FC < 0) or unchanged (all remaining regions). Sequences of 200 bp centered on the peak summit were scanned with the PWMs of the curated families only. The enrichment was thus computed using the three classes (Up, down, NS), each one being tested against the union of the two other classes with background sequences weighted to match the foreground GC and k-mer composition. P-values were adjusted with the Benjamini-Hochberg method over the curated set of motifs and motifs with adjusted p-value<0.05 were considered enriched, reported as log2 enrichment for the gained and lost classes in each contrast.

The alternative *Ciita* promoters were defined directly from the GENCODE vM35 annotation. Transcripts sharing the common second exon were grouped, and their first-exon TSSs were clustered into promoter groups. This recovered promoter I (*Ciita*-205/207 transcript isoforms), promoter III (*Ciita*-201/202 transcript isoforms) and promoter IV (Ciita-203 transcript isoform). To describe motif content at individual loci of interest (*Ciita* and *Cd40*), consensus regions within +/− 25kb of these genes were scanned with the curated PWMs described above using the monaLisa R package, using a minimum score of 6. Motif occurrences per region are reported in locus view, alongside the region’s accessibility signal and gene bodies. To generate the locus views, per-sample coverage was computed from the filtered BAM files and scaled by the edgeR’s TMM factor so that tracks match the normalization used for the differential testing. Normalized coverage was average per group and plotted together with gene models from the GENCODE annotation.

### BeatAML2 and TCGA analysis

Clinical data, raw and normalized expression of the BeatAML2.0 were acquired from the biodev page related to the second BeatAML manuscript^45^ (https://biodev.github.io/BeatAML2/). Samples were filtered to retain only those with high level of blast composition (BM blast percentage >80%) to reduce potential source of confounding due to sample composition heterogeneity. For consistency, both cohorts were preprocessed and analyzed in a similar fashion. Genes with insufficient expression were removed using *edgeR*’s *filterByExpr* method^76^, retaining transcripts with a minimum count of 10 in at least as many samples as the smallest group and a minimum total count of 15 across all samples (BeatAML2.0: 16190/ 63676 genes retained; TCGA 13067/19944 genes retained). log_2_CPM normalization was applied to the filtered dataset. Samples with systematic missing covariate data (*FLT3*-ITD, *NPM1*, *RUNX1*, *ASXL1*, *TP53*, cohort wave, blast percentage, or specimen type) were excluded. Sample-level quality control including principal component analysis and pairwise sample correlation clustering on the top 2000 most variable genes were performed to screen for batch effects and outlier structure. Differential gene expression analysis was performed on both cohort (BeatAML2.0; n=191, TCGA; n=52 samples) using normalized (log_2_CPM, prior count = 2) *SLC6A8* expression as a continuous predictor in a linear model adjusting for covariates identified in each cohort. Overall, these covariates consisted in mutation status (*FLT3*-ITD, *NPM1*, *RUNX1*, *ASXL1*, *TP53*), clinical confounding (percentage blasts), and for BeatAML2.0, sequencing cohort wave and sample specimen type. Mean-variance modeling was performed using the *voom* method (*limma* R package), followed by linear model fitting (*lmFit*) and empirical Bayes moderation of variance (*eBayes*). Differential expression statistics coefficients were extracted and filtered based on Benjamini-Hochberg corrected p-values (FDR<0.05). In such model, the logFC represents the increase in a given gene when *SLC6A8* expression doubles. To make it more meaningful and more similar to a standard logFC range from a group-based differential expression, we transformed the logFC coefficient to an interquartile range; in this scale, a logFC of 1 represents a 2-fold increase between the 25^th^ and 75^th^ percentile patients in view of their *SLC6A8* gene expression. GSEA was performed on the *SLC6A8* differential expression results using *clusterProfiler* R package^91^. Genes were ranked by their test statistics from the *limma* model and gene sets collection were drawn from seven MSigDB collections (version 2025.1) selected for their relevance for the current analysis (Hallmark, C2:CGP, KEGG, Reactome, WikiPathways, C5:GO:BP and ImmuneSigDB). Gene sets were filtered to those with 15 to 500 genes in the corresponding lists. The reported pathways reached significance after Benjamini-Hochberg FDR correction.

An erythroid differentiation signature was constructed by filtering gene sets from the MSigDB resource using a curated list of keywords targeting erythroid and heme-related terms (e.g. “erythroid”, “erythroblast”, “hemoglobin”, etc.) applied to both gene set names and descriptions. Gene sets with less than 5 genes or more than 500 genes present in the filtered expression matrix were removed. Per-sample enrichment scores for each retained gene set were calculated using the gene set variation analysis (GSVA) algorithm. A consensus erythroid score was then computed for each sample by calculating the average GSVA enrichment score of all retained gene sets. Both the erythroid score and the *SLC6A8* expression were adjusted against the same covariates used in the *limma* model to again display their association without confounding factors.

To rule out the possibility that the erythroid – *SLC6A8* association is driven by erythroid compositions in the BeatAML2/ TCGA samples, we inferred the proportion of cell-types in the samples and regressed out the proportion of erythroid lineage from the *SLC6A8* expression and from the erythroid signature score. The cell-type deconvolution was performed using CIBERSORTx (https://cibersortx.stanford.edu/)^47^. One BM and one peripheral blood reference single-cell RNA-seq dataset were used, depending on the origin of the samples to deconvolute. The BM reference dataset was the AML single-cell atlas of Whittle et al. (2026)^48^ containing 748’679 cells from multiples studies with healthy and AML BM samples. We used the broad cell type annotation as provided by the original authors. The peripheral blood reference is a combined reference from the 10x Genomics blood cells from Tabula Sapiens^92^, containing healthy samples, complemented by leukemic and leukemic-like cells from the BM atlas. Using CIBERSORTx, we built a signature matrix and imputed cell fractions for each reference, BM and PB samples run separately. The settings set to non-default were; disabling quantile normalization, using S-mode batch correction to account for technical differences in the single-cell and bulk data, and running 100 permutations for p-value estimation. All resulting deconvolution inferences were returned significant (p<0.01).

### Single-cell AML atlas analysis

Single-cell expression data were obtained from a published BM scRNA-seq atlas of AML from Whittle et al. (2026)^48^. The atlas integrated 748’679 cells from 20 published scRNA-seq studies, covering 159 AML patients at diagnosis and 51 healthy donors. We downloaded the processed AnnData object from figshare (https://doi.org/10.48420/27269946). We used the atlas’ normalized expression values of SLC6A8 and the published scVI UMAP coordinates as provided. Each cell was assigned to one of three categories, based on the atlas’s original annotations: 1) Erythroid, regrouping erythroid cell-types plus the early and late erythroid HSPC sub-populations, 2) leukemic, the LSC and leukemic-like HSPC sub-populations labels (HSC-like, Prog-like, GMPlike, etc.), 3) Other, all remaining cell-types.

### Code and data availability

Raw datasets have been deposited and will be publicly released upon publication in a peer-reviewed journal. Code used to support the findings is available at github.com/apostolovalab/slc6a8_aml_manuscript.

## References

1. Marine, J.C., Dawson, S.J., and Dawson, M.A. (2020). Non-genetic mechanisms of therapeutic resistance in cancer. Nat Rev Cancer 20, 743–756. 10.1038/s41568-020-00302-4.

2. Sharma, S.V., Lee, D.Y., Li, B., Quinlan, M.P., Takahashi, F., Maheswaran, S., McDermott, U., Azizian, N., Zou, L., Fischbach, M.A., et al. (2010). A chromatin-mediated reversible drug-tolerant state in cancer cell subpopulations. Cell 141, 69–80. 10.1016/j.cell.2010.02.027.

3. Shaffer, S.M., Dunagin, M.C., Torborg, S.R., Torre, E.A., Emert, B., Krepler, C., Beqiri, M., Sproesser, K., Brafford, P.A., Xiao, M., et al. (2017). Rare cell variability and drug-induced reprogramming as a mode of cancer drug resistance. Nature 546, 431–435. 10.1038/nature22794.

4. Faubert, B., Solmonson, A., and DeBerardinis, R.J. (2020). Metabolic reprogramming and cancer progression. Science 368. 10.1126/science.aaw5473.

5. Hangauer, M.J., Viswanathan, V.S., Ryan, M.J., Bole, D., Eaton, J.K., Matov, A., Galeas, J., Dhruv, H.D., Berens, M.E., Schreiber, S.L., et al. (2017). Drug-tolerant persister cancer cells are vulnerable to GPX4 inhibition. Nature 551, 247–250. 10.1038/nature24297.

6. Oren, Y., Tsabar, M., Cuoco, M.S., Amir-Zilberstein, L., Cabanos, H.F., Hutter, J.C., Hu, B., Thakore, P.I., Tabaka, M., Fulco, C.P., et al. (2021). Cycling cancer persister cells arise from lineages with distinct programs. Nature 596, 576–582. 10.1038/s41586-021-03796-6.

7. Munoz, D.P., Yannone, S.M., Daemen, A., Sun, Y., Vakar-Lopez, F., Kawahara, M., Freund, A.M., Rodier, F., Wu, J.D., Desprez, P.Y., et al. (2019). Targetable mechanisms driving immunoevasion of persistent senescent cells link chemotherapy-resistant cancer to aging. JCI Insight 5. 10.1172/jci.insight.124716.

8. Chaib, S., Lopez-Dominguez, J.A., Lalinde-Gutierrez, M., Prats, N., Marin, I., Boix, O., Garcia-Garijo, A., Meyer, K., Munoz, M.I., Aguilera, M., et al. (2024). The efficacy of chemotherapy is limited by intratumoral senescent cells expressing PD-L2. Nat Cancer 5, 448–462. 10.1038/s43018-023-00712-x.

9. Dohner, H., Wei, A.H., Appelbaum, F.R., Craddock, C., DiNardo, C.D., Dombret, H., Ebert, B.L., Fenaux, P., Godley, L.A., Hasserjian, R.P., et al. (2022). Diagnosis and management of AML in adults: 2022 recommendations from an international expert panel on behalf of the ELN. Blood 140, 1345–1377. 10.1182/blood.2022016867.

10. Shimony, S., Stahl, M., and Stone, R.M. (2025). Acute Myeloid Leukemia: 2025 Update on Diagnosis, Risk-Stratification, and Management. Am J Hematol 100, 860–891. 10.1002/ajh.27625.

11. Sweeney, C., and Vyas, P. (2019). The Graft-Versus-Leukemia Effect in AML. Front Oncol 9, 1217. 10.3389/fonc.2019.01217.

12. Dickinson, A.M., Norden, J., Li, S., Hromadnikova, I., Schmid, C., Schmetzer, H., and Jochem-Kolb, H. (2017). Graft-versus-Leukemia Effect Following Hematopoietic Stem Cell Transplantation for Leukemia. Front Immunol 8, 496. 10.3389/fimmu.2017.00496.

13. Spellman, S.R., Xu, K., Oloyede, T., Ahn, K.W., Akhtar, O., Bolon, Y.T., Broglie, L., Bloomquist, J., Bupp, C., Chen, M., et al. (2025). Current Activity Trends and Outcomes in Hematopoietic Cell Transplantation and Cellular Therapy – A Report from the CIBMTR. Transplant Cell Ther 31, 505–532. 10.1016/j.jtct.2025.05.014.

14. Bejanyan, N., Weisdorf, D.J., Logan, B.R., Wang, H.L., Devine, S.M., de Lima, M., Bunjes, D.W., and Zhang, M.J. (2015). Survival of patients with acute myeloid leukemia relapsing after allogeneic hematopoietic cell transplantation: a center for international blood and marrow transplant research study. Biol Blood Marrow Transplant 21, 454–459. 10.1016/j.bbmt.2014.11.007.

15. Horowitz, M., Schreiber, H., Elder, A., Heidenreich, O., Vormoor, J., Toffalori, C., Vago, L., and Kroger, N. (2018). Epidemiology and biology of relapse after stem cell transplantation. Bone Marrow Transplant 53, 1379–1389. 10.1038/s41409-018-0171-z.

16. Jones, C.L., Stevens, B.M., Pollyea, D.A., Culp-Hill, R., Reisz, J.A., Nemkov, T., Gehrke, S., Gamboni, F., Krug, A., Winters, A., et al. (2020). Nicotinamide Metabolism Mediates Resistance to Venetoclax in Relapsed Acute Myeloid Leukemia Stem Cells. Cell Stem Cell 27, 748–764 e744. 10.1016/j.stem.2020.07.021.

17. Farge, T., Saland, E., de Toni, F., Aroua, N., Hosseini, M., Perry, R., Bosc, C., Sugita, M., Stuani, L., Fraisse, M., et al. (2017). Chemotherapy-Resistant Human Acute Myeloid Leukemia Cells Are Not Enriched for Leukemic Stem Cells but Require Oxidative Metabolism. Cancer Discov 7, 716–735. 10.1158/2159-8290.CD-16-0441.

18. Stevens, B.M., Jones, C.L., Pollyea, D.A., Culp-Hill, R., D’Alessandro, A., Winters, A., Krug, A., Abbott, D., Goosman, M., Pei, S., et al. (2020). Fatty acid metabolism underlies venetoclax resistance in acute myeloid leukemia stem cells. Nat Cancer 1, 1176–1187. 10.1038/s43018-020-00126-z.

19. van Gastel, N., Spinelli, J.B., Sharda, A., Schajnovitz, A., Baryawno, N., Rhee, C., Oki, T., Grace, E., Soled, H.J., Milosevic, J., et al. (2020). Induction of a Timed Metabolic Collapse to Overcome Cancer Chemoresistance. Cell Metab 32, 391–403 e396. 10.1016/j.cmet.2020.07.009.

20. Gallipoli, P., Giotopoulos, G., Tzelepis, K., Costa, A.S.H., Vohra, S., Medina-Perez, P., Basheer, F., Marando, L., Di Lisio, L., Dias, J.M.L., et al. (2018). Glutaminolysis is a metabolic dependency in FLT3(ITD) acute myeloid leukemia unmasked by FLT3 tyrosine kinase inhibition. Blood 131, 1639–1653. 10.1182/blood-2017-12-820035.

21. Burk, A.C., and Apostolova, P. (2024). Metabolic instruction of the graft-versus-leukemia immunity. Front Immunol 15, 1347492. 10.3389/fimmu.2024.1347492.

22. Uhl, F.M., Chen, S., O’Sullivan, D., Edwards-Hicks, J., Richter, G., Haring, E., Andrieux, G., Halbach, S., Apostolova, P., Buscher, J., et al. (2020). Metabolic reprogramming of donor T cells enhances graft-versus-leukemia effects in mice and humans. Sci Transl Med 12. 10.1126/scitranslmed.abb8969.

23. Mussai, F., De Santo, C., Abu-Dayyeh, I., Booth, S., Quek, L., McEwen-Smith, R.M., Qureshi, A., Dazzi, F., Vyas, P., and Cerundolo, V. (2013). Acute myeloid leukemia creates an arginase-dependent immunosuppressive microenvironment. Blood 122, 749–758. 10.1182/blood-2013-01-480129.

24. Mussai, F., Egan, S., Higginbotham-Jones, J., Perry, T., Beggs, A., Odintsova, E., Loke, J., Pratt, G., U, K.P., Lo, A., et al. (2015). Arginine dependence of acute myeloid leukemia blast proliferation: a novel therapeutic target. Blood 125, 2386–2396. 10.1182/blood-2014-09-600643.

25. Zorko, N.A., Bernot, K.M., Whitman, S.P., Siebenaler, R.F., Ahmed, E.H., Marcucci, G.G., Yanes, D.A., McConnell, K.K., Mao, C., Kalu, C., et al. (2012). Mll partial tandem duplication and Flt3 internal tandem duplication in a double knock-in mouse recapitulates features of counterpart human acute myeloid leukemias. Blood 120, 1130–1136. 10.1182/blood-2012-03-415067.

26. Fleming, R.A., Capizzi, R.L., Rosner, G.L., Oliver, L.K., Smith, S.J., Schiffer, C.A., Silver, R.T., Peterson, B.A., Weiss, R.B., Omura, G.A., and, et al. (1995). Clinical pharmacology of cytarabine in patients with acute myeloid leukemia: a cancer and leukemia group B study. Cancer Chemother Pharmacol 36, 425–430. 10.1007/BF00686192.

27. Hande, K.R., Stein, R.S., McDonough, D.A., Greco, F.A., and Wolff, S.N. (1982). Effects of high-dose cytarabine. Clin Pharmacol Ther 31, 669–674. 10.1038/clpt.1982.93.

28. Derissen, E.J., Beijnen, J.H., and Schellens, J.H. (2013). Concise drug review: azacitidine and decitabine. Oncologist 18, 619–624. 10.1634/theoncologist.2012-0465.

29. Paul, C., Baurain, R., Gahrton, G., and Peterson, C. (1980). Determination of daunorubicin and its main metabolites in plasma, urine and leukaemic cells in patients with acute myeloblastic leukaemia. Cancer Lett 9, 263–269. 10.1016/0304-3835(80)90016-6.

30. Pei, S., Minhajuddin, M., Callahan, K.P., Balys, M., Ashton, J.M., Neering, S.J., Lagadinou, E.D., Corbett, C., Ye, H., Liesveld, J.L., et al. (2013). Targeting aberrant glutathione metabolism to eradicate human acute myelogenous leukemia cells. J Biol Chem 288, 33542–33558. 10.1074/jbc.M113.511170.

31. Jones, C.L., Stevens, B.M., D’Alessandro, A., Culp-Hill, R., Reisz, J.A., Pei, S., Gustafson, A., Khan, N., DeGregori, J., Pollyea, D.A., and Jordan, C.T. (2019). Cysteine depletion targets leukemia stem cells through inhibition of electron transport complex II. Blood 134, 389–394. 10.1182/blood.2019898114.

32. Moreno Vanegas, Y., and Badar, T. (2022). Clinical Utility of Azacitidine in the Management of Acute Myeloid Leukemia: Update on Patient Selection and Reported Outcomes. Cancer Manag Res 14, 3527–3538. 10.2147/CMAR.S271442.

33. Mopin, A., Driss, V., and Brinster, C. (2016). A Detailed Protocol for Characterizing the Murine C1498 Cell Line and its Associated Leukemia Mouse Model. J Vis Exp. 10.3791/54270.

34. Dai, W., Vinnakota, S., Qian, X., Kunze, D.L., and Sarkar, H.K. (1999). Molecular characterization of the human CRT-1 creatine transporter expressed in Xenopus oocytes. Arch Biochem Biophys 361, 75–84. 10.1006/abbi.1998.0959.

35. Sora, I., Richman, J., Santoro, G., Wei, H., Wang, Y., Vanderah, T., Horvath, R., Nguyen, M., Waite, S., Roeske, W.R., and, et al. (1994). The cloning and expression of a human creatine transporter. Biochem Biophys Res Commun 204, 419–427. 10.1006/bbrc.1994.2475.

36. Tachikawa, M., Fujinawa, J., Takahashi, M., Kasai, Y., Fukaya, M., Sakai, K., Yamazaki, M., Tomi, M., Watanabe, M., Sakimura, K., et al. (2008). Expression and possible role of creatine transporter in the brain and at the blood-cerebrospinal fluid barrier as a transporting protein of guanidinoacetate, an endogenous convulsant. J Neurochem 107, 768–778. 10.1111/j.1471-4159.2008.05652.x.

37. Nagana Gowda, G.A., Pascua, V., Killion, C.E., Paranji, R.K., and Raftery, D. (2023). Labile Metabolite Profiling in Human Blood Using Phosphorus NMR Spectroscopy. Anal Chem 95, 15033–15041. 10.1021/acs.analchem.3c03040.

38. Wallimann, T., Tokarska-Schlattner, M., and Schlattner, U. (2011). The creatine kinase system and pleiotropic effects of creatine. Amino Acids 40, 1271–1296. 10.1007/s00726-011-0877-3.

39. Haas, R.C., and Strauss, A.W. (1990). Separate nuclear genes encode sarcomere-specific and ubiquitous human mitochondrial creatine kinase isoenzymes. J Biol Chem 265, 6921–6927.

40. Hossle, J.P., Schlegel, J., Wegmann, G., Wyss, M., Bohlen, P., Eppenberger, H.M., Wallimann, T., and Perriard, J.C. (1988). Distinct tissue specific mitochondrial creatine kinases from chicken brain and striated muscle with a conserved CK framework. Biochem Biophys Res Commun 151, 408–416. 10.1016/0006-291x(88)90608-0.

41. Wallimann, T., Wyss, M., Brdiczka, D., Nicolay, K., and Eppenberger, H.M. (1992). Intracellular compartmentation, structure and function of creatine kinase isoenzymes in tissues with high and fluctuating energy demands: the ‘phosphocreatine circuit’ for cellular energy homeostasis. Biochem J 281 ( *Pt 1*), 21–40. 10.1042/bj2810021.

42. Di Biase, S., Ma, X., Wang, X., Yu, J., Wang, Y.C., Smith, D.J., Zhou, Y., Li, Z., Kim, Y.J., Clarke, N., et al. (2019). Creatine uptake regulates CD8 T cell antitumor immunity. J Exp Med 216, 2869–2882. 10.1084/jem.20182044.

43. Samborska, B., Roy, D.G., Rahbani, J.F., Hussain, M.F., Ma, E.H., Jones, R.G., and Kazak, L. (2022). Creatine transport and creatine kinase activity is required for CD8(+) T cell immunity. Cell Rep 38, 110446. 10.1016/j.celrep.2022.110446.

44. Kurth, I., Yamaguchi, N., Andreu-Agullo, C., Tian, H.S., Sridhar, S., Takeda, S., Gonsalves, F.C., Loo, J.M., Barlas, A., Manova-Todorova, K., et al. (2021). Therapeutic targeting of SLC6A8 creatine transporter suppresses colon cancer progression and modulates human creatine levels. Sci Adv 7, eabi7511. 10.1126/sciadv.abi7511.

45. Bottomly, D., Long, N., Schultz, A.R., Kurtz, S.E., Tognon, C.E., Johnson, K., Abel, M., Agarwal, A., Avaylon, S., Benton, E., et al. (2022). Integrative analysis of drug response and clinical outcome in acute myeloid leukemia. Cancer Cell 40, 850–864 e859. 10.1016/j.ccell.2022.07.002.

46. Cancer Genome Atlas Research, N., Ley, T.J., Miller, C., Ding, L., Raphael, B.J., Mungall, A.J., Robertson, A., Hoadley, K., Triche, T.J., Jr., Laird, P.W., et al. (2013). Genomic and epigenomic landscapes of adult de novo acute myeloid leukemia. N Engl J Med 368, 2059–2074. 10.1056/NEJMoa1301689.

47. Newman, A.M., Steen, C.B., Liu, C.L., Gentles, A.J., Chaudhuri, A.A., Scherer, F., Khodadoust, M.S., Esfahani, M.S., Luca, B.A., Steiner, D., et al. (2019). Determining cell type abundance and expression from bulk tissues with digital cytometry. Nat Biotechnol 37, 773–782. 10.1038/s41587-019-0114-2.

48. Whittle, J., Meyer, S., Lacaud, G., Murtuza-Baker, S., and Iqbal, M. (2026). Single-cell atlas of AML reveals age-related gene regulatory networks in t(8;21) AML. Elife 14. 10.7554/eLife.104978.

49. Zhang, Y., Newsom, K.J., Zhang, M., Kelley, J.S., and Starostik, P. (2022). GATM-Mediated Creatine Biosynthesis Enables Maintenance of FLT3-ITD-Mutant Acute Myeloid Leukemia. Mol Cancer Res 20, 293–304. 10.1158/1541-7786.MCR-21-0314.

50. Loo, J.M., Scherl, A., Nguyen, A., Man, F.Y., Weinberg, E., Zeng, Z., Saltz, L., Paty, P.B., and Tavazoie, S.F. (2015). Extracellular metabolic energetics can promote cancer progression. Cell 160, 393–406. 10.1016/j.cell.2014.12.018.

51. Patel, R., Ford, C.A., Rodgers, L., Rushworth, L.K., Fleming, J., Mui, E., Zhang, T., Watson, D., Lynch, V., Mackay, G., et al. (2022). Cyclocreatine Suppresses Creatine Metabolism and Impairs Prostate Cancer Progression. Cancer Res 82, 2565–2575. 10.1158/0008-5472.CAN-21-1301.

52. Rashidi, A., Billingham, L.K., Zolp, A., Chia, T.Y., Silvers, C., Katz, J.L., Park, C.H., Delay, S., Boland, L., Geng, Y., et al. (2024). Myeloid cell-derived creatine in the hypoxic niche promotes glioblastoma growth. Cell Metab 36, 62–77 e68. 10.1016/j.cmet.2023.11.013.

53. Bao, Y., Qiao, J., Gong, W., Zhang, R., Zhou, Y., Xie, Y., Xie, Y., He, J., and Yin, T. (2024). Spatial metabolomics highlights metabolic reprogramming in acute myeloid leukemia mice through creatine pathway. Acta Pharm Sin B 14, 4461–4477. 10.1016/j.apsb.2024.07.004.

54. Zhang, L., Zhu, Z., Yan, H., Wang, W., Wu, Z., Zhang, F., Zhang, Q., Shi, G., Du, J., Cai, H., et al. (2021). Creatine promotes cancer metastasis through activation of Smad2/3. Cell Metab 33, 1111–1123 e1114. 10.1016/j.cmet.2021.03.009.

55. Li, Q., Liu, M., Sun, Y., Jin, T., Zhu, P., Wan, X., Hou, Y., and Tu, G. (2021). SLC6A8-mediated intracellular creatine accumulation enhances hypoxic breast cancer cell survival via ameliorating oxidative stress. J Exp Clin Cancer Res 40, 168. 10.1186/s13046-021-01933-7.

56. Zhou, X., Wang, G., Wu, Y., Wu, M., Zhai, X., Tian, C., Prochownik, E.V., Jiang, C., and Li, Y. (2025). SLC6A8-mediated creatine uptake suppresses ERK2-FSP1 signaling and induces ferroptosis in colorectal cancer. Cell Rep 44, 116139. 10.1016/j.celrep.2025.116139.

57. Fenouille, N., Bassil, C.F., Ben-Sahra, I., Benajiba, L., Alexe, G., Ramos, A., Pikman, Y., Conway, A.S., Burgess, M.R., Li, Q., et al. (2017). The creatine kinase pathway is a metabolic vulnerability in EVI1-positive acute myeloid leukemia. Nat Med 23, 301–313. 10.1038/nm.4283.

58. Benajiba, L., Alexe, G., Su, A., Raffoux, E., Soulier, J., Hemann, M.T., Hermine, O., Itzykson, R., Stegmaier, K., and Puissant, A. (2019). Creatine kinase pathway inhibition alters GSK3 and WNT signaling in EVI1-positive AML. Leukemia 33, 800–804. 10.1038/s41375-018-0291-x.

59. Darabedian, N., Ji, W., Fan, M., Lin, S., Seo, H.S., Vinogradova, E.V., Yaron, T.M., Mills, E.L., Xiao, H., Senkane, K., et al. (2023). Depletion of creatine phosphagen energetics with a covalent creatine kinase inhibitor. Nat Chem Biol 19, 815–824. 10.1038/s41589-023-01273-x.

60. Ji, L., Zhao, X., Zhang, B., Kang, L., Song, W., Zhao, B., Xie, W., Chen, L., and Hu, X. (2019). Slc6a8-Mediated Creatine Uptake and Accumulation Reprogram Macrophage Polarization via Regulating Cytokine Responses. Immunity 51, 272–284 e277. 10.1016/j.immuni.2019.06.007.

61. Chen, S., Ma, X., Liu, Y., Zhong, Z., Wei, C., Li, M., and Zhu, X. (2024). Creatine Promotes Endometriosis by Inducing Ferroptosis Resistance via Suppression of PrP. Adv Sci (Weinh) 11, e2403517. 10.1002/advs.202403517.

62. Peng, M., Ren, J., Jing, Y., Jiang, X., Xiao, Q., Huang, J., Tao, Y., Lei, L., Wang, X., Yang, Z., et al. (2021). Tumour-derived small extracellular vesicles suppress CD8+ T cell immune function by inhibiting SLC6A8-mediated creatine import in NPM1-mutated acute myeloid leukaemia. J Extracell Vesicles 10, e12168. 10.1002/jev2.12168.

63. Edwards-Hicks, J., Apostolova, P., Buescher, J.M., Maib, H., Stanczak, M.A., Corrado, M., Klein Geltink, R.I., Maccari, M.E., Villa, M., Carrizo, G.E., et al. (2023). Phosphoinositide acyl chain saturation drives CD8(+) effector T cell signaling and function. Nat Immunol 24, 516–530. 10.1038/s41590-023-01419-y.

64. Buenrostro, J.D., Giresi, P.G., Zaba, L.C., Chang, H.Y., and Greenleaf, W.J. (2013). Transposition of native chromatin for fast and sensitive epigenomic profiling of open chromatin, DNA-binding proteins and nucleosome position. Nat Methods 10, 1213–1218. 10.1038/nmeth.2688.

65. Team, R.C. (2020). R: A Language and Environment for Statistical Computing. R Foundation for Statistical Computing, Vienna, Austria.

66. Do, K.T., Wahl, S., Raffler, J., Molnos, S., Laimighofer, M., Adamski, J., Suhre, K., Strauch, K., Peters, A., Gieger, C., et al. (2018). Characterization of missing values in untargeted MS-based metabolomics data and evaluation of missing data handling strategies. Metabolomics 14, 128. 10.1007/s11306-018-1420-2.

67. Armitage, E.G., Godzien, J., Alonso-Herranz, V., Lopez-Gonzalvez, A., and Barbas, C. (2015). Missing value imputation strategies for metabolomics data. Electrophoresis 36, 3050–3060. 10.1002/elps.201500352.

68. Wei, R., Wang, J., Su, M., Jia, E., Chen, S., Chen, T., and Ni, Y. (2018). Missing Value Imputation Approach for Mass Spectrometry-based Metabolomics Data. Sci Rep 8, 663. 10.1038/s41598-017-19120-0.

69. van den Berg, R.A., Hoefsloot, H.C., Westerhuis, J.A., Smilde, A.K., and van der Werf, M.J. (2006). Centering, scaling, and transformations: improving the biological information content of metabolomics data. BMC Genomics 7, 142. 10.1186/1471-2164-7-142.

70. Gu, Z., Eils, R., and Schlesner, M. (2016). Complex heatmaps reveal patterns and correlations in multidimensional genomic data. Bioinformatics 32, 2847–2849.

71. 10.1093/bioinformatics/btw313.

71. Conway, J.R.L., A., Gehlenborg, N. (2017). UpSetR: an R package for the visualization of intersecting sets and their properties. Bioinformatics 33, 2938–2940.

72. Chen, S. (2023). Ultrafast one-pass FASTQ data preprocessing, quality control, and deduplication using fastp. Imeta 2, e107. 10.1002/imt2.107.

73. Dobin, A., Davis, C.A., Schlesinger, F., Drenkow, J., Zaleski, C., Jha, S., Batut, P., Chaisson, M., and Gingeras, T.R. (2013). STAR: ultrafast universal RNA-seq aligner. Bioinformatics 29, 15–21. 10.1093/bioinformatics/bts635.

74. Liu, D. (2019). Algorithms for efficiently collapsing reads with Unique Molecular Identifiers. PeerJ 7, e8275. 10.7717/peerj.8275.

75. Liao, Y., Smyth, G.K., and Shi, W. (2014). featureCounts: an efficient general purpose program for assigning sequence reads to genomic features. Bioinformatics 30, 923–930. 10.1093/bioinformatics/btt656.

76. Robinson, M.D., McCarthy, D.J., and Smyth, G.K. (2010). edgeR: a Bioconductor package for differential expression analysis of digital gene expression data. Bioinformatics 26, 139–140. 10.1093/bioinformatics/btp616.

77. Law, C.W., Chen, Y., Shi, W., and Smyth, G.K. (2014). voom: Precision weights unlock linear model analysis tools for RNA-seq read counts. Genome Biol 15, R29. 10.1186/gb-2014-15-2-r29.

78. Wu, D., and Smyth, G.K. (2012). Camera: a competitive gene set test accounting for inter-gene correlation. Nucleic Acids Res 40, e133. 10.1093/nar/gks461.

79. Badia, I.M.P., Velez Santiago, J., Braunger, J., Geiss, C., Dimitrov, D., Muller-Dott, S., Taus, P., Dugourd, A., Holland, C.H., Ramirez Flores, R.O., and Saez-Rodriguez, J. (2022). decoupleR: ensemble of computational methods to infer biological activities from omics data. Bioinform Adv 2, vbac016. 10.1093/bioadv/vbac016.

80. Muller-Dott, S., Tsirvouli, E., Vazquez, M., Ramirez Flores, R.O., Badia, I.M.P., Fallegger, R., Turei, D., Laegreid, A., and Saez-Rodriguez, J. (2023). Expanding the coverage of regulons from high-confidence prior knowledge for accurate estimation of transcription factor activities. Nucleic Acids Res 51, 10934–10949. 10.1093/nar/gkad841.

81. Zou, Z., Ohta, T., Kasukawa, T., and Oki, S. (2026). ChIP-Atlas 2025 update: 10-year anniversary of a data-mining platform for exploring epigenomic landscape. Nucleic Acids Res 54, W77–W84. 10.1093/nar/gkag378.

82. Lo Surdo, P., Iannuccelli, M., Karis, K., Meo, E., Omidi, P., Tosoni, M., Graziosi, S., Panni, S., Fiorillo, M.T., Licata, L., et al. (2026). SIGNOR 4.0: the 2025 update with focus on phosphorylation data. Nucleic Acids Res 54, D682–D690. 10.1093/nar/gkaf1237.

84. Ewels, P.A., Peltzer, A., Fillinger, S., Patel, H., Alneberg, J., Wilm, A., Garcia, M.U., Di Tommaso, P., and Nahnsen, S. (2020). The nf-core framework for community-curated bioinformatics pipelines. Nat Biotechnol 38, 276–278. 10.1038/s41587-020-0439-x.

85. Di Tommaso, P., Chatzou, M., Floden, E.W., Barja, P.P., Palumbo, E., and Notredame, C. (2017). Nextflow enables reproducible computational workflows. Nat Biotechnol 35, 316–319. 10.1038/nbt.3820.

85. Frankish, A., Diekhans, M., Ferreira, A.M., Johnson, R., Jungreis, I., Loveland, J., Mudge, J.M., Sisu, C., Wright, J., Armstrong, J., et al. (2019). GENCODE reference annotation for the human and mouse genomes. Nucleic Acids Res 47, D766–D773. 10.1093/nar/gky955.

86. Stark, R.B., G. (2011). DiffBind: differential binding analysis of ChIP-Seq peak data.

87. Robinson, M.D., and Oshlack, A. (2010). A scaling normalization method for differential expression analysis of RNA-seq data. Genome Biol 11, R25. 10.1186/gb-2010-11-3-r25.

88. Wang, Q., Li, M., Wu, T., Zhan, L., Li, L., Chen, M., Xie, W., Xie, Z., Hu, E., Xu, S., and Yu, G. (2022). Exploring Epigenomic Datasets by ChIPseeker. Curr Protoc 2, e585. 10.1002/cpz1.585.

89. Machlab, D., Burger, L., Soneson, C., Rijli, F.M., Schubeler, D., and Stadler, M.B. (2022). monaLisa: an R/Bioconductor package for identifying regulatory motifs. Bioinformatics 38, 2624–2625. 10.1093/bioinformatics/btac102.

90. Rauluseviciute, I., Riudavets-Puig, R., Blanc-Mathieu, R., Castro-Mondragon, J.A., Ferenc, K., Kumar, V., Lemma, R.B., Lucas, J., Cheneby, J., Baranasic, D., et al. (2024). JASPAR 2024: 20th anniversary of the open-access database of transcription factor binding profiles. Nucleic Acids Res 52, D174–D182. 10.1093/nar/gkad1059.

91. Yu, G., Wang, L.G., Han, Y., and He, Q.Y. (2012). clusterProfiler: an R package for comparing biological themes among gene clusters. OMICS 16, 284–287. 10.1089/omi.2011.0118.

93. Tabula Sapiens, C., Jones, R.C., Karkanias, J., Krasnow, M.A., Pisco, A.O., Quake, S.R., Salzman, J., Yosef, N., Bulthaup, B., Brown, P., et al. (2022). The Tabula Sapiens: A multiple-organ, single-cell transcriptomic atlas of humans. Science 376,eabl4896. 10.1126/science.abl4896.

